# Functional interrogation uncovers a critical role for a high-plasticity cell state in lung adenocarcinoma

**DOI:** 10.1101/2025.10.03.680232

**Authors:** Jason E. Chan, Chun-Hao Pan, Jonathan Rub, Klavdija Krause, Emma Brown, Gary Guzman, Zeda Zhang, Hannah Styers, Griffin Hartmann, Zhuxuan Li, Xueqian Zhuang, Scott W. Lowe, Doron Betel, Yan Yan, Tuomas Tammela

**Affiliations:** Cancer Biology and Genetics Program, Sloan Kettering Institute, Memorial Sloan Kettering Cancer Center, New York, New York 10065, USA; Division of Solid Tumor Oncology, Department of Medicine, Memorial Sloan Kettering Cancer Center, New York, New York 10065, USA; Tri-I Program in Computational Biology & Medicine, Weill Cornell Medicine, New York, New York 10065, USA; BCMB Allied Program, Weill Cornell Graduate School of Medical Science, Weill Cornell Medicine, New York, New York 10065, USA; Howard Hughes Medical Institute, Chevy Chase, MD, USA; Division of Hematology & Medical Oncology, Department of Medicine, Weill Cornell Medicine, New York, New York 10065, USA; Institute for Computational Biomedicine, Weill Cornell Medicine, New York, New York 10065, USA; College of Biomedicine and Health and College of Life Science and Technology, Huazhong Agricultural University, Wuhan, Hubei 430070, China; Hubei Hongshan Laboratory, Wuhan, Hubei 430070, China

## Abstract

Plasticity—the ability of cells to undergo phenotypic transitions—drives cancer progression and therapy resistance^1–3^. To date, strategies targeting cancer plasticity have not advanced to the clinic due to a lack of fundamental understanding of the underlying mechanisms. Recent studies have suggested that plasticity in solid tumors is concentrated in a minority subset of cancer cells^4–6^, yet functional studies interrogating this high plasticity cell state (HPCS) *in situ* are lacking. Here, we developed mouse models enabling detection, longitudinal lineage tracing, and ablation of the HPCS in autochthonous lung tumors *in vivo*. Using lineage tracing, we uncover the HPCS cells are dedifferentiated but possess high capacity for cell state transitions, giving rise to both early neoplastic (differentiated) and advanced lung cancer cell states *in situ*. Longitudinal lineage tracing using secreted luciferases reveals HPCS-derived cells harbor high capacity for growth when compared to bulk cancer cells or another defined cancer cell state with features of differentiated lung epithelium. Suicide gene-mediated ablation of the HPCS in early neoplasias abrogates tumor progression. Ablating HPCS cells in established tumors by suicide gene or HPCS-directed CAR T cells robustly reduces tumor burden, whereas ablation of a differentiated lung cancer cell state had no effect. We further demonstrate that the HPCS gives rise to therapy-resistant cell states, whereas ablation of the HPCS abrogates resistance to chemotherapy and oncoprotein-targeted therapy. Interestingly, an HPCS-like state is ubiquitous in regenerating epithelia and in carcinomas of multiple other tissues, revealing a convergence of plasticity programs. Our work establishes the HPCS as a critical hub enabling reciprocal transitions between cancer cell states, including acquisition of states adapted to cancer therapies. Targeting the HPCS in lung cancer and in other carcinomas may suppress cancer progression and eradicate treatment resistance.

## Introduction

Plasticity promotes tumor progression by enabling cancer cells to acquire states with a high capacity to proliferate, metastasize, and adapt to stress^1,2^. Moreover, plasticity promotes resistance to chemotherapy^4,7^ and oncoprotein-targeted therapy^8–10^ in multiple cancer types, allowing cancer cells to acquire new cell states adapted to withstand therapeutic pressure via nongenetic mechanisms^1,2^. In addition to directly enabling adaptation to therapies, plasticity is a requisite for the emergence and maintenance of intra-tumoral heterogeneity, which increases the likelihood of cancer cell states harboring intrinsic resistance to therapy. Thus, plasticity remains one of the most fundamental problems in cancer biology and one of the foremost challenges in clinical cancer management today. Despite the increasingly recognized importance of cellular plasticity for tumor progression and treatment resistance, it is not apparent, neither conceptually nor practically, how targeting cancer plasticity would be best achieved—i.e., should the focus be on subsets of cells or rather on specific molecular programs.

The application of single-cell genomics and associated computational approaches over the past decade has enabled the unsupervised mapping of malignant cell states at an unprecedented scale and resolution^11,12^. Yet most experimental and clinically based single-cell analyses remain descriptive and are focused on a static measurement, largely due to gaps in available technologies. Although limited to static snapshots, single-cell mRNA sequencing (scRNA-seq) studies have identified candidate transitional states in multiple solid tumor types^4,11–16^. Collectively, these studies suggest plasticity is concentrated in specific subsets of cancer cells, warranting their functional interrogation in time-dynamic experiments. As plasticity in cancer is fundamentally a temporal problem, new experimental strategies and model systems enabling elucidation of the dynamic nature of plastic transitions are needed.

Lung adenocarcinoma (LUAD) is a prototype of a common, lethal, and therapy-resistant solid tumor^17,18^. Important insights into the biology of human LUAD have emerged from the use of genetically engineered mouse models (GEMMs). In the most commonly used LUAD GEMMs, viral expression of *Cre* or *Flp* recombinase in lung epithelial cells leads to somatic activation of oncogenic **<u>K</u>**RAS(G12D) and deletion of **<u>p</u>**53 (***KP***)^19–21^. The *KP* model incorporates *de novo* LUAD development in the relevant tissue site, recapitulating key molecular and histopathological features of the human disease, including responses to chemotherapy and KRAS-targeted therapy^9,22^.

We recently applied scRNA-seq to construct a map of LUAD evolution from the alveolar type 2 (AT2) cell of origin to advanced adenocarcinoma in the *KP* model^4^. This analysis identified a previously unknown cancer cell state, which emerges in early stages of lung tumorigenesis and is maintained in LUAD tumors throughout cancer progression. This cell state harbors transcriptomic features that are drastically different from AT2 cells—most notably the concurrent expression of gene programs associated with a wide range of cellular identities ranging from trophoblast stem cells, chondroblasts, and endothelial cells to lung and kidney epithelium^4^. Intriguingly, these transcriptomic features are distinct from published gene expression signatures of adult tissue stem cells or cancer stem cells (CSCs)^4^. Computational modeling of cancer cell state differentiation trajectories implicated this cell state as a key transition point in LUAD progression^4^. Given these attributes, we nominated this cellular subset as a candidate high-plasticity cell state (HPCS). Important recent work uncovered a cell state analogous to the HPCS in human early-stage lung neoplasias^5^, underscoring the utility of the *KP* model for studying the HPCS. Together, these findings suggest plasticity in lung cancer is concentrated in the HPCS subset, motivating functional interrogation of this state and its potential causal role in distinct steps of LUAD progression and in the context of therapy-associated transitions to drug-resistant cell states.

## Results

### Development of novel multifunctional reporter systems enabling visualization, ablation, and longitudinal lineage tracing of the HPCS *in situ* in tumors

To functionally interrogate the HPCS *in situ* at distinct stages of LUAD evolution, we developed two reporter systems allowing us to visualize, isolate, lineage-trace, and ablate the HPCS cells. We identified *Slc4a11* as the most specific gene marking the HPCS in autochthonous mouse *KP* LUAD tumors (**Fig. 1a-b; Extended Data Fig. 1a; Supplementary Table 1**)^4^. *Slc4a11* encodes a sodium and hydroxyl-coupled borate cotransporter, whose expression is restricted to corneal endothelial cells, inner ear, and medullary kidney epithelium in adults^23–25^. To mark, lineage-trace, and ablate the *Slc4a11^+^* HPCS cells, we assembled a cDNA reporter cassette comprising ***<u>m</u>****Scarlet* bright red monomeric fluorescent protein^26^, tamoxifen-activatable Cre recombinase (***<u>C</u>****reERT2*)^27^, and human diphtheria toxin (DT) receptor (***<u>D</u>****TR*)^28^ (***MCD*** cassette). We knocked the *MCD* cassette into the *Slc4a11* locus in mouse embryonic stem cells (mESCs) harboring Flp recombinase-activatable KRAS(G12D) (*Kras^frt-stop-frt(FSF)-G12D/+^*)^21^ and p53 loss-of-function (*Trp53^frt/frt^*)^29^ alterations (***KPfrt***) and generated mESC-derived chimeras (**Extended Data Fig. 1b-d**). Unlike humans, mouse cells are not sensitive to DT, an extremely potent cytotoxic peptide, allowing us to selectively ablate *Slc4a11^+^* HPCS cells by systemic administration of DT to mice bearing *KPfrt; Slc4a11^MCD/+^* lung tumors^28^. To restrict the expression of the *MCD* cassette to *Slc4a11^+^* cancer cells, we introduced a frt-stop-frt (*FSF*) cassette^21^ upstream of the *MCD*. Upon removal of the *FSF* cassette, *MCD* is linked to the last coding exon of *Slc4a11* by a 2A peptide sequence^30^ (**Fig. 1c; Extended Data Fig. 1b**). This approach preserves activity of the endogenous gene and links the *MCD* reporter tightly to native gene-regulatory elements, increasing the fidelity of the reporter.

**Figure 1.**
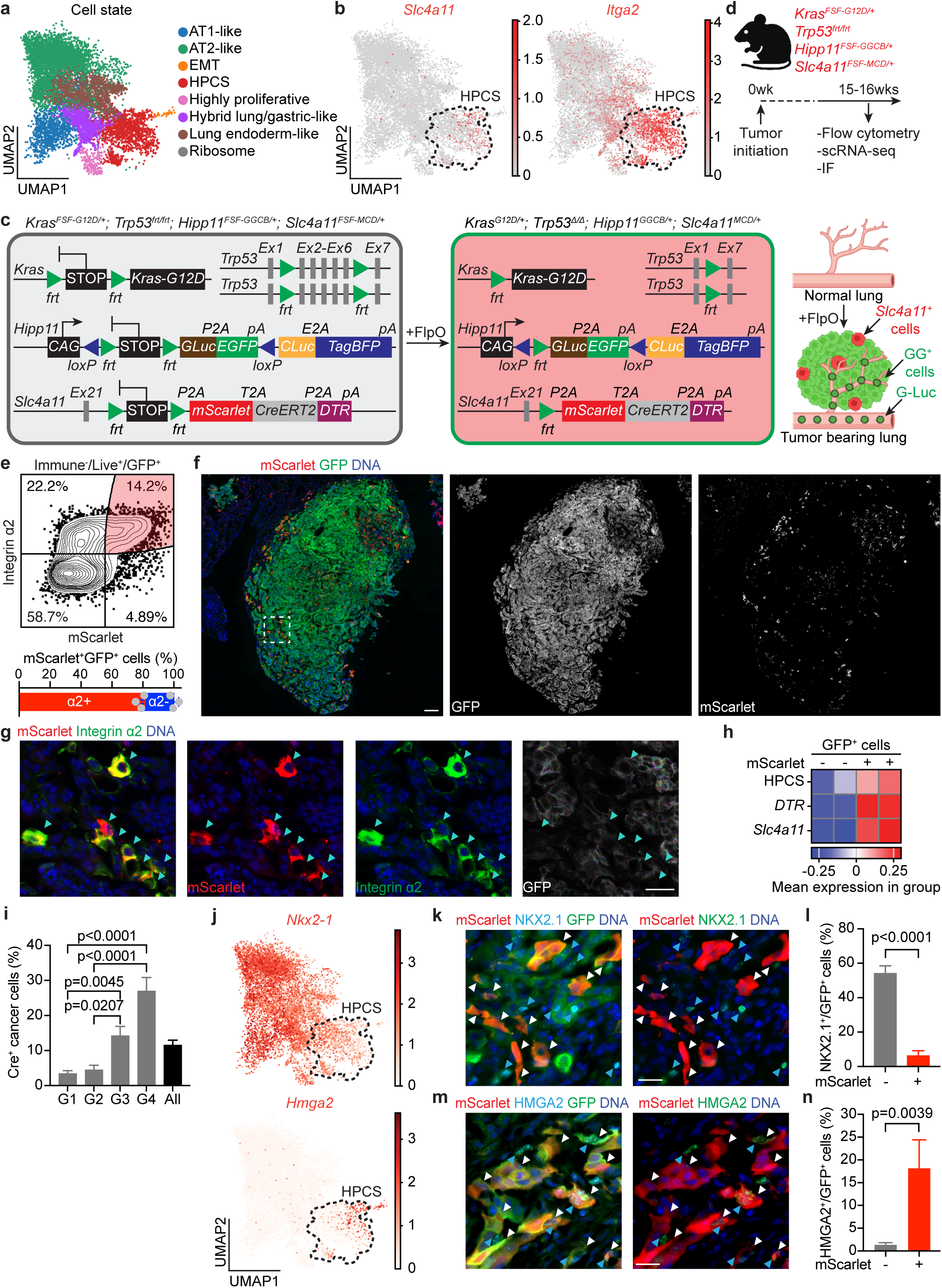
The *Slc4a11^MCD/+^*reporter system faithfully marks the HPCS *in vivo*. **a,** Single-cell transcriptomics showing eight annotated lung adenocarcinoma (LUAD) cell states that molecularly define LUAD tumors collected from six *KP* mice 15-16 weeks post-tumor initiation (PTI). **b,** Distribution of *Slc4a11* (*left*) and *Itga2* (*right*) gene expression. **c,** *Left*: Schematic of *Kras^FSF-G12D/+^; Trp53^frt/frt^; Hipp11^FSF-GGCB/+^; Slc4a11^FSF-MCD/+^* alleles before and after FlpO-mediated recombination. *Right*: Delivery of FlpO recombinase in the lung results in lung tumors expressing E<u>G</u>FP and *Gaussia* secreted luciferase, <u>G</u>-Luc (GG^+^, green). *Slc4a11*^+^ tumor cells are marked by <u>m</u>Scarlet, <u>C</u>reER^T2^ and <u>D</u>TR (MCD, red). **d**, Experimental design to validate the *Slc4a11^FSF-MCD/+^* reporter. Mice were intubated with PGK-FlpO lentivirus and LUAD tumors were harvested 15-16 weeks PTI for flow cytometry, single-cell RNA sequencing (scRNA-seq), and immunofluorescence (IF). **e**, *Top*: Representative flow cytometry plot showing co-expression of mScarlet and integrin α2 in GFP^+^ tumor cells (highlighted region). Cells were gated as CD45^-^/CD31^-^/CD11b^-^/CD11c^-^/F4/80^-^/TER-119^-^/Helix NP NIR^-^ (live)/GFP^+^ cells. *Bottom*: Average percentage of integrin α2^+^ cells in the mScarlet^+^/GFP^+^ cell pool gated as above. N = 3 mice. **f**, IF showing mScarlet^+^ cells (red) within a GFP^+^ LUAD tumor (green). Scale bar: 100 µm. **g**, Co-localization of mScarlet with integrin α2 (turquoise arrowheads) in GFP^+^ cancer cells. Images are magnified from the tumor region marked by dashed rectangles are shown in (**f**). Scale bar: 20 µm. **h,** Heatmap of expression values for the HPCS gene expression program, the *Slc4a11^MCD^* reporter cassette (as measured by *DTR* expression), and the native *Slc4a11* transcript in sorted mScarlet^-^/GFP^+^ and mScarlet^+^/GFP^+^ LUAD cancer cells. **i**, Percentage of Cre^+^ (HPCS) cancer cells across distinct histopathological grades in autochthonous *KPfrt; Slc4a11^MCD/+^* lung tumors. N = 28 tumors from 5 mice. One-way ANOVA. **j**, Distribution of *Nkx2-1* (*top*) and *Hmga2* (*bottom*) gene expression across the LUAD cell states shown in (**a**). **k**, Representative images showing mScarlet and NKX2.1 expression in GFP^+^ LUAD cells in *KPfrt*; *Hipp11^GGCB/+^*; *Slc4a11^MCD/+^* lung tumors at 16 weeks PTI. Note mutual exclusivity of mScarlet (white arrowheads) and NKX2.1 (blue arrowheads). Scale bar: 20 µm. **l,** Percentage of NKX2.1^+^ cells in mScarlet^-^/GFP^+^ vs. mScarlet^+^/GFP^+^ LUAD cell subsets. N = 53 tumors from 5 mice. Mann-Whitney U test. **m,** Representative images showing mScarlet and HMGA2 expression in GFP^+^ LUAD cells in *KPfrt*; *Hipp11^GGCB/+^*; *Slc4a11^MCD/+^*lung tumors at 16 weeks PTI. Note co-expression of both mScarlet (white arrowheads) and HMGA2 (blue arrowheads) proteins in a subset of LUAD cells. Scale bar: 20 µm. **n**, Percentage of HMGA2^+^ cells in mScarlet^-^/GFP^+^ vs. mScarlet^+^/GFP^+^ LUAD cell subsets. N = 21 tumors from 5 mice. Mann-Whitney U test. Error bars show SEM.

We also developed a reporter system marking cells undergoing Flp and/or Cre recombinase activity, enabling longitudinal monitoring of the growth potential of the HPCS and other cell states *in vivo*. Our system is comprised of a Flp-inducible *Gaussia* luciferase (**<u>G</u>-Luc**^31^) linked to E**<u>G</u>**FP^32^ (***GG***) and Cre-inducible *Cypridina* luciferase (**<u>C</u>-Luc**^33^) linked to Tag**<u>B</u>**FP^34^ (***CB***) (**Fig. 1c**). Both luciferases are naturally secreted, enabling their detection from a small volume of plasma using specific substrates^35^. This allows non-terminal longitudinal sampling in the same animal to investigate the long-term proliferative potential of lineage-traced cells *in vivo*. The “*GGCB*” cassette was knocked into the ubiquitously active *Hipp11* locus^36^ in *KPfrt* mESCs, followed by generation of mESC-derived chimeras (**Extended Data Fig. 2a-c**). To validate the functionality of the *GGCB* cassette, we performed *ex vivo* transformation of AT2 cells^37^ isolated from a *KPfrt; Hipp11^FSF-GGCB/+^* chimeric mouse using lentiviral vectors encoding either codon-optimized Flp recombinase (*FlpO*) alone or *FlpO* linked to *CreERT2*. In these experiments, FlpO activates oncogenic KRAS(G12D), deletes p53, and initiates expression of the *GG* cassette, whereas subsequent activation of CreER^T2^ with 4-hydroxytamoxifen (4-OHT) results in a switch from *GG* to *CB* (**Extended Data Fig. 2d**). G-Luc activity was increased 13 days post-transformation in all conditions (**Extended Data Fig. 2e**, upper panel), whereas C-Luc activity was only observed following 4-OHT stimulation in organoids transduced with the vector encoding both *FlpO* and *CreERT2* (**Extended Data Fig. 2e**, lower panel). In addition, we performed flow cytometry and fluorescence imaging analyses on organoids under these four conditions. We found that EGFP was expressed at baseline following *FlpO* and a switch to TagBFP was only observed upon 4-OHT exposure in the organoids transduced with both *FlpO* and *CreERT2* (**Extended Data Fig. 2f-g**). Similar results were obtained in subcutaneous transplants (**Extended Data Fig. 3a-c**) and autochthonous lung tumors (**Extended Data Fig. 3d-g**) *in vivo*, both by detection of G-Luc and C-Luc from repeated blood samples and by fluorescence imaging of tumors at endpoint. Taken together, these results demonstrate the *Hipp11^FSF-GGCB/+^* allele is a specific and sensitive reporter of Flp and Cre recombinase activity, which can be utilized to longitudinally track growth of tumors initiated by Flp recombinase activity and growth potential of cell states marked by Cre recombinase activity.

### The Slc4a11^MCD/+^ reporter system faithfully marks the HPCS in situ

To interrogate the HPCS in autochthonous lung tumors *in situ*, we next crossed the *Hipp11^FSF-GGCB/+^* and the *Slc4a11^FSF-MCD/+^* alleles together to obtain *KPfrt; Hipp11^GGCB/+^; Slc4a11^MCD/+^* mice. In these mice, intra-tracheal intubation of viral FlpO recombinase induces *GG^+^ KP* tumors where the *Slc4a11^+^* HPCS cells are marked by mScarlet (*MCD*) expression (**Fig. 1c**). To determine whether the activated *Slc4a11^MCD^* allele faithfully marks the HPCS in autochthonous LUAD, we initiated lung tumors in *KPfrt; Hipp11^FSF-GGCB/+^; Slc4a11^FSF-MCD/+^* mice using lentiviral FlpO recombinase and harvested lung tumors at 15 to 16 weeks post-tumor initiation (PTI) (**Fig. 1d**). Flow cytometry revealed the average proportion of mScarlet^+^ cells to be 16.98 ± 4.29% of the total GFP^+^ pool of cancer cells (**Fig. 1e**, *top*), closely mirroring the proportion of HPCS cells detected by scRNA-seq (13.1 ± 3.80%) (**Fig. 1a, b**)^4^. The vast majority (78.41 ± 1.66%) of the mScarlet^+^ cells expressed integrin α2 (**Fig. 1e**, *bottom*; **Extended Data Fig. 4a**), another sensitive, but less specific marker of the HPCS (**Fig. 1b; Supplementary Table 1**)^4^. Spatial analysis in tissue sections revealed mScarlet^+^ cells form a subset of *KP* LUAD tumors *in situ,* localizing as small clusters or single cells throughout the lung tumors (**Fig. 1f**). The mScarlet^+^ cells express integrin α2 (**Fig. 1g**), corroborating our flow cytometry results. Robust enrichment of the HPCS gene expression signature^4^ and *Slc4a11* expression was observed in isolated mScarlet^+^ cells (hereafter referred to as HPCS cells) when compared to mScarlet^-^ cells (**Fig. 1h**). Furthermore, scRNA-seq analysis of GFP^+^ tumor cells revealed concordant expression of the *Slc4a11^MCD^* allele (marked by *DTR)* with *Slc4a11* and *Itga2* endogenous transcripts within cancer cells (**Extended Data Fig. 4b**). Our analysis of the scRNA-seq data revealed the specificity of the *Slc4a11^MCD^* reporter cassette to be 84.25-99.82% for the HPCS in autochthonous *KPfrt; Hipp11^GGCB/+^; Slc4a11^MCD/+^* lung tumors (**Extended Data Fig. 4; Supplementary Table 2**). In sum, these results establish the *Slc4a11^MCD^* allele as a faithful reporter of the HPCS *in situ*.

We next leveraged the *Slc4a11^MCD/+^* reporter to molecularly and functionally characterize the HPCS. Lung cancer progression is defined by loss of lung epithelial identity, marked by downregulation of the lung epithelial master regulator NKX2.1 and acquisition of programs associated with the embryonic foregut, marked by induction of HMGA2^38,39^. We found the density of the malignant cells in the HPCS increased with histopathological grade (**Fig. 1i**), which was associated with a reduced density of NKX2.1-positive cells and an increase in HMGA2-positive cells (**Extended Data Fig. 5**)^38,39^. A more granular analysis using scRNA-seq and immunofluorescence revealed *Nkx2-1* expression is associated with early neoplastic cell states with an alveolar or lung endoderm identity (**Fig. 1j**, *top*), whereas the HPCS exhibits reduced *Nkx2-1* expression and an induction of *Hmga2* (**Fig. 1j**, *bottom*; **Fig. 1k-n**). A computational time series analysis of our original longitudinal scRNA-seq data spanning early neoplasias and advanced adenocarcinomas^4^ corroborated that the *Nkx2-1*-to-*Hmga2* switch occurs in the HPCS (marked by *Slc4a11* and *Plaur*) (**Extended Data Fig. 6**). Furthermore, this analysis suggests two long-term trajectories of lung cancer evolution emanating from the HPCS, one that maintains *Hmga2* expression and eventually undergoes epithelial-mesenchymal transition (EMT) and another that re-acquires *Nkx2-1* expression, loses *Hmga2*, and ends up in a proximal ciliated lung epithelial-like state (**Extended Data Fig. 6**). These data suggest the HPCS functions as a key transition between the alveolar states in early neoplasias and the cell states that emerge later in lung cancer progression.

### The HPCS is a hub for cell state transitions in LUAD

To functionally interrogate the transition potential—i.e. plasticity—of the HPCS *in situ*, we lineage-traced the HPCS in autochthonous *KPfrt; Rosa26^mTmG/+^; Slc4a11^MCD/+^* lung adenomas and in established lung adenocarcinomas at 6 or 12 weeks PTI, respectively, with a single pulse of tamoxifen (**Fig. 2a**). Flow cytometry analysis revealed faithful labeling of the HPCS cells at 3 days post-labeling (**Fig. 2b**), the time required for washout of tamoxifen and its active metabolite 4-OHT. After 14 days of tracing, we found the percentage of cells actively in the HPCS (mScarlet^+^) in the total pool of traced (GFP^+^) cells decreased precipitously compared to the 3-day baseline at both the 6➔8 and 12➔14 week tracing windows (**Fig. 2b**), indicating that the majority of the traced cells exited the HPCS and acquired new fates during the 14-day trace. Notably, the fraction of HPCS cells over total cancer cells (mScarlet^+^/EpCAM^+^) remained stable over time, but the fraction of traced cells over total EpCAM^+^ cells—enriched for malignant cells—in microdissected tumors expanded over time, indicative of clonal expansion (**Extended Data Fig. 7a-d**). ScRNA-seq analysis of sorted traced (GFP^+^) cells at 3 days post-tamoxifen in both early (6-week) and advanced (12-week) tumors showed that >93% and >67% of the traced cells occupied the HPCS, respectively (**Fig. 2d, e, g, h; Extended Data Fig. 7e, f**). These data indicate the *Slc4a11^MCD/+^* reporter system affords lineage tracing of the HPCS at high specificity, although transitions out of the HPCS may occur more rapidly in adenocarcinomas, at the 12➔14 week tracing window (**Fig. 2b, g, h; Extended Data Fig. 7e, f**).

**Figure 2.**
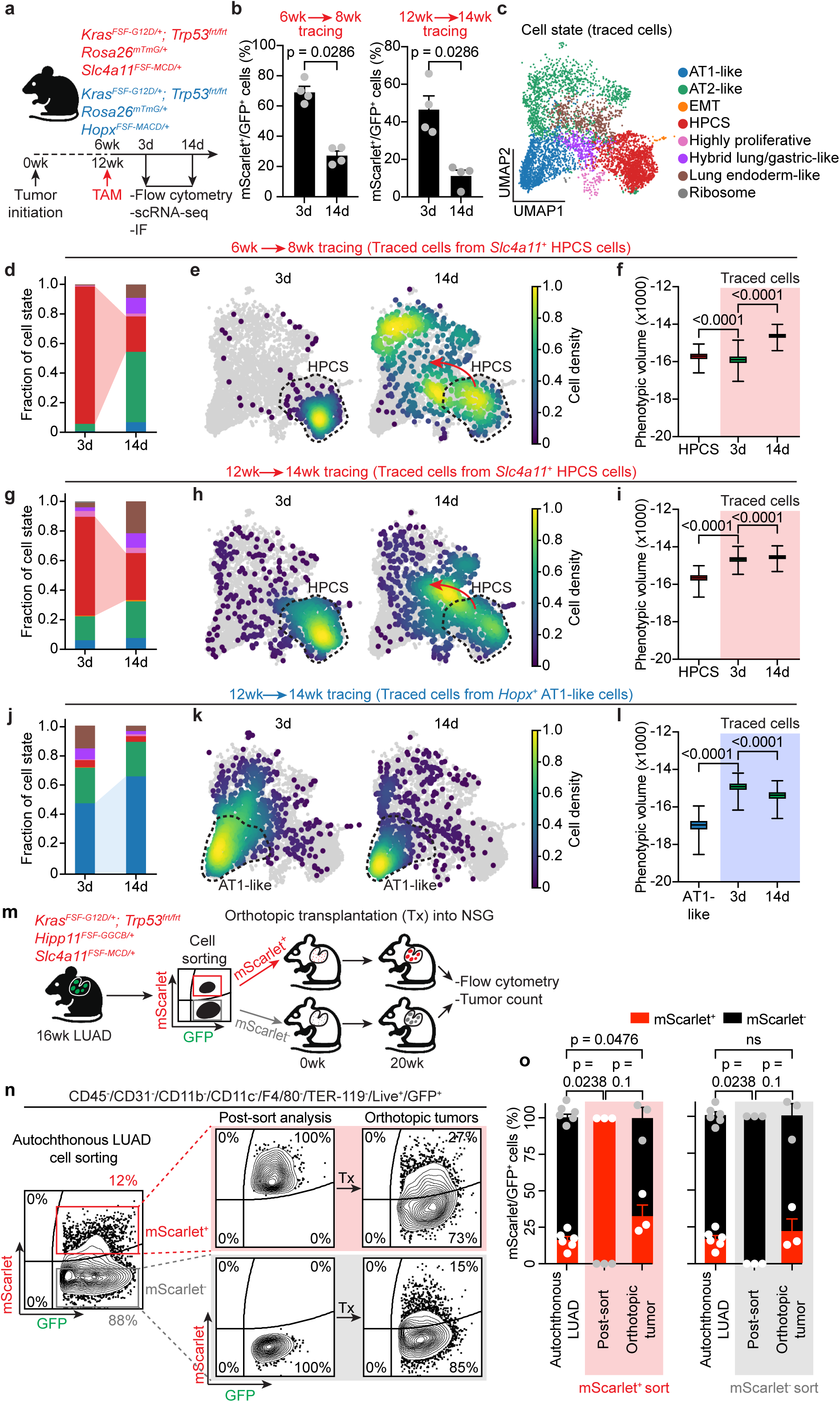
The HPCS is a hub for cell state transitions *in vivo*. **a**, Experimental design for lineage tracing HPCS in *KPfrt; Rosa26^mTmG/+^; Slc4a11^FSF-MCD/+^* mice and AT-1 like cells in *KPfrt; Rosa26^mTmG/+^; Hopx^FSF-MACD/+^* mice harboring autochthonous LUAD tumors. **b,** mScarlet^+^/GFP^+^ (HPCS/all traced) cells at week 6 (*left*) or 12 (*right*) PTI after 3d or 14d of tracing. N = 4 mice per time point. Mann Whitney U test. **c**, Cell states in traced GFP^+^ LUAD tumor cells [CD45^-^/CD31^-^/CD11b^-^/CD11c^-^/F4/80^-^/TER-119^-^/Helix NP NIR^-^(Live^+^)/GFP^+^ cells] harvested at 6 to 8 weeks or 12 to 14 weeks PTI, defined using scRNA-seq. **d-f,** Transcriptomes of HPCS-derived cells traced at 6 weeks PTI harvested at 3d or 14d following tracing. **d**, Cell state identities of HPCS-derived traced cells at 3d and 14d after lineage tracing, colored as in (**c**). **e,** Density of traced cells in UMAP space after 3d or 14d of tracing. The HPCS is outlined. Red arrow indicates trans-differentiation of HPCS cells into other LUAD cell states. **f**, Quantification of cell state diversity as measured by the phenotypic volumes of cells in the HPCS compared to traced LUAD cells harvested after 3d or 14d of tracing. Box plots are min-max. One-way ANOVA. **g-i**, Transcriptomes of HPCS-derived cells traced at 12 weeks PTI harvested at 3d or 14d following tracing. **g**, Cell state identities of HPCS-derived traced cells at 3d and 14d after lineage tracing, colored as in (**c**). **h**, Density of traced cells in UMAP space after 3d or 14d of tracing. The HPCS is outlined. Red arrow indicates trans-differentiation of HPCS cells into other LUAD cell states. **i**, Phenotypic volume of cells in the HPCS compared to traced LUAD cells harvested after 3d and 14d of tracing. Box plots are min-max. One-way ANOVA. **j-l**, Cell state identities of cells derived from *Hopx^+^* AT1-like cells traced at 12 weeks PTI and harvested at 3d and 14d after lineage tracing, colored as in (**c**). **j**, Cell state identities of AT1-like-derived traced cells at 3d and 14d after lineage tracing, colored as in (**c**). **k,** Density of traced cells in UMAP space after 3d or 14d of tracing. The AT1-like state is outlined. **l**, Phenotypic volume of cells in the AT1-like state compared to traced LUAD cells harvested after 3d and 14d of tracing. Box plots are min-max. One-way ANOVA. **m**, Experimental design of orthotopic transplantation assays to assess bidirectional plasticity between HPCS and non-HPCS states. See text. Tx: transplantation. **n,** Flow cytometry analysis of mScarlet and GFP expression in primary autochthonous LUAD tumors (*left*), sorted mScarlet^+^/GFP^+^ and mScarlet^−^/GFP^+^ populations (post-sort analysis, *middle*), and orthotopically transplanted tumors (*right*). Cells were gated as CD45^−^/CD31^−^/CD11b^−^/CD11c^−^/F4/80^−^/TER-119^−^/DAPI^−^ (live)/GFP^+^ cells. **o,** Proportion of mScarlet^+^ and mScarlet^-^ cells within GFP^+^ populations in orthotopically transplanted tumors derived from either mScarlet^+^/GFP^+^ (*left*) or mScarlet^−^/GFP^+^ (*right*) cells. The proportion from autochthonous LUAD and sorted cells prior to transplantation (mScarlet^+^/GFP^+^ and mScarlet^−^/GFP^+^) were included as reference. N = 3-5 mice per group. Mann Whitney U test. Error bars are SEM.

To identify the fates acquired by the cells derived from the HPCS, we performed scRNA-seq on the total pool of traced cells at 3 and 14 days post-tracing in adenomas (6 weeks PTI) and in established adenocarcinomas (12 weeks PTI) (**Fig. 2a, c-i**). Remarkably, analysis of the transcriptomes of traced cells revealed the HPCS gives rise to all cancer cell states observed in both adenomas (6→8 weeks) and the adenocarcinomas (12→14 weeks) (**Fig. 2d, e, g, h; Extended Data Fig. 7e, f**), indicating the HPCS is critical for the maintenance of intra-tumoral heterogeneity and cell state composition in LUAD tumors. In line with this finding, a significant fraction of the HPCS-derived cells acquired expression of the differentiated lung epithelial markers *Nkx2-1/*NKX2.1, *Sftpc*/SPC, and *Hopx* and lost expression of *Hmga2*/HMGA2 in both adenomas and adenocarcinomas at 14 days following lineage tracing compared to the 3-day baseline (**Extended Data Fig. 8a-d**). Taken together, these data indicate the HPCS functions as a hub for cell state transitions between the early neoplastic, alveolar-like cell states and the advanced, endoderm-like states. Surprisingly, these transitions occur via the HPCS in both “forward” and “reverse” directions along the axis of cancer progression (**Extended Data Fig. 6**), towards both the alveolar and endoderm-like states, even in advanced adenocarcinomas. This acquisition of cellular diversity by the HPCS-derived cells in both adenomas and adenocarcinomas translated into robust, statistically significant increases in phenotypic volume, a quantitative measure of the diversity of cellular phenotypes within cell populations^40^ (**Fig. 2f, i**). Notably, the phenotypic volumes of the traced cells at 14 days post tamoxifen exposure are significantly higher than the subset of HPCS cells alone or the 3-day labeling baseline (**Fig. 2f, i**), suggesting that some traced cells exit the HPCS already during the 3-day tracing period, especially in the advanced tumors in the 12→14 week tracing window. These findings thus implicate the HPCS as a central driver of malignant cell state diversity in lung cancer.

To compare the plasticity of the HPCS to another defined, differentiated cancer cell state, we generated a novel reporter allele by knocking a *FSF-mScarlet-AkaLuc-CreER-DTR* (*FSF-MACD*) cassette into the *Hopx* locus in *KPfrt* mESCs (**Extended Data Fig. 9a-c)**. *Hopx* marks AT1 cells in the normal lung and is enriched in the AT1-like cell state in LUAD tumors^4,9^, although we note that its specificity to the AT1-like state is lower (82.2%) than *Slc4a11*’s for the HPCS (99.7%), as *Hopx* is also expressed in a subset of AT2-like cancer cells (**Extended Data Fig. 9d-j; Supplementary Table 2; Supplementary Table 3**). We first generated *KPfrt; Hipp11^FSF-GGCB/+^; Hopx^FSF-MACD/+^* mice and induced autochthonous tumors using lentiviral FlpO recombinase (**Extended Data Fig. 9d**). In these mice, LUAD cells expressing *Hopx* can be tracked by *in vivo* imaging of the super-bright luciferase AkaLuc^41^ (**Extended Data Fig. 9e**), in addition to mScarlet fluorescence. An analysis of microdissected tumors at 16 weeks PTI revealed robust tumor-specific mScarlet fluorescence, which is strongly enriched for *Hopx* mRNA and HOPX protein as well as the AT1-like state transcriptomic signature (**Extended Data Fig. 9f-j**). Thus, the *KPfrt; Hipp11^GGCB/+^; Hopx^MACD/+^*model enables faithful tracking of the AT1-like cell state *in situ* in LUAD tumors.

To measure plasticity, or differentiation potential, of the AT1-like cells we performed similar lineage tracing studies as for the HPCS in established autochthonous *KPfrt; Rosa26^mTmG/+^; Hopx^MACD/+^*adenocarcinomas (**Fig. 2a; Extended Data Fig. 10a**). We found the fraction of mScarlet^+^ cells remained constant in both the traced fraction and the total pool of cancer cells in the *KPfrt; Rosa26^mTmG/+^; Hopx^MACD/+^* adenocarcinomas (**Extended Data Fig. 10b, c**). Further, the fraction of traced AT1-like cells in the total pool of cancer cells declined over time (**Extended Data Fig. 10d**), suggesting clonal contraction. These results are markedly different from the HPCS tracing experiments, which showed rapid exit of the reporter-positive cells from the traced fraction and clonal expansion in adenocarcinomas (**Fig. 2b**; **Extended Data Fig. 7d**). ScRNA-seq analysis of the lineage-traced AT1-like cells 3 days following a single pulse of tamoxifen (baseline) showed 47.2 ± 7.73% of the traced cells reside in the AT1-like cell state, indicating relatively specific labeling of the AT1-like state, although, as expected, some labeling of AT2-like cells was also observed (**Fig. 2j, k; Extended Data Fig. 10e, f**). In notable contrast to the HPCS tracing experiments, the traced AT1-like cells remained in the AT1-like state and the phenotypic volume of the traced cells in fact contracted over the 12→14 week tracing window (**Fig. 2j-l; Extended Data Fig. 10e, f**). These results are consistent with a model where high plasticity is concentrated in the HPCS, which gives rise to cancer cell states with fixed phenotypes and low plasticity.

### The HPCS can be acquired by non-HPCS cells

Our results indicate that the fraction of the malignant cells in the HPCS remains stable during the 6→8 and 12→14 week tracing windows (**Extended Data Fig. 7a, c**), and, in fact, increases with higher histopathological grade (**Fig. 1i**). However, our tracing results indicate the traced *Slc4a11^+^*cells rapidly exit the HPCS (**Fig. 2b, d-i**). This suggested that in order to maintain the fraction of HPCS cells stable over time, non-HPCS may harbor the capacity to acquire the HPCS. To test this hypothesis, we transplanted primary non-HPCS (mScarlet^-^/GFP^+^) cells from autochthonous *KPfrt; Hipp11^GGCB/+^; Slc4a11^MCD/+^* lung tumors orthotopically into lungs of recipient mice; pure HPCS (mScarlet^+^/GFP^+^) cells were transplanted as a control (**Fig. 2m-o**). In line with a high growth potential (**Extended Data Fig. 7b, d**), the HPCS cells were significantly more (∼9x) efficient at forming tumors than the non-HPCS cells (**Extended Data Fig. 11a**). Interestingly, tumors arising from the non-HPCS population exhibited a very similar fraction of mScarlet^+^ cells as the autochthonous parent tumors (**Fig. 2n, o**), indicating that non-HPCS cells can acquire the HPCS. Similar results were obtained in subcutaneous serial allograft experiments (**Extended Data Fig. 11b-d**). Collectively, these data demonstrate that the HPCS is not a stem-like state that can only arise via self-renewal of pre-existing HPCS cells, but rather dynamically interconverts between non-HPCS states. This bidirectional plasticity likely enables the maintenance and expansion of the HPCS during cancer progression.

### The HPCS drives tumor growth and progression

To interrogate growth potential of the HPCS *in situ*, we leveraged the *KPfrt; Hipp11^FSF-GGCB/+^; Slc4a11^FSF-MCD/+^*model, where the relative contribution of the HPCS cells vs. other cancer cell states to tumor growth can be longitudinally recorded by measuring the ratio of traced (CB^+^) and non-traced (GG^+^) cancer cells through repeated measurement of the C-Luc/G-Luc ratio in blood samples (**Fig. 3a**). As controls, we used *KPfrt; Hipp11^FSF-GGCB/+^; Rosa26^FSF-CreERT2/+^*mice to randomly label cancer cells (using a low dose of tamoxifen, as before^42^) and the *KPfrt; Hipp11^FSF-GGCB/+^; Hopx^FSF-MACD/+^* mice to label the AT1-like cell state (**Fig. 3b**). Longitudinal lineage tracing via secreted luciferases in established LUAD at 14 ➔ 16 weeks post-tumor initiation revealed that cancer cells derived from the HPCS harbor significantly higher growth potential and capacity for clonal expansion when compared to derivatives of randomly labeled cells (**Fig. 3c**). In contrast, derivatives of the differentiated *Hopx^+^* AT1-like cell state harbor low growth potential (**Fig. 3c**), suggesting the AT1-like cell state is outcompeted in LUAD tumors. These findings are consistent with the flow cytometry analyses showing clonal expansion of HPCS-derived traced cells and contraction of the AT1-like-derived traced cells (**Extended Data Fig. 7b, d; Extended Data Fig. 10d**) as well as the HPCS vs. non-HPCS allotransplantation experiments (**Fig. 2m-o**; **Extended Data Fig. 11b-d**). Taken together, these findings demonstrate that the HPCS exhibits robust growth potential in established LUAD.

**Figure 3.**
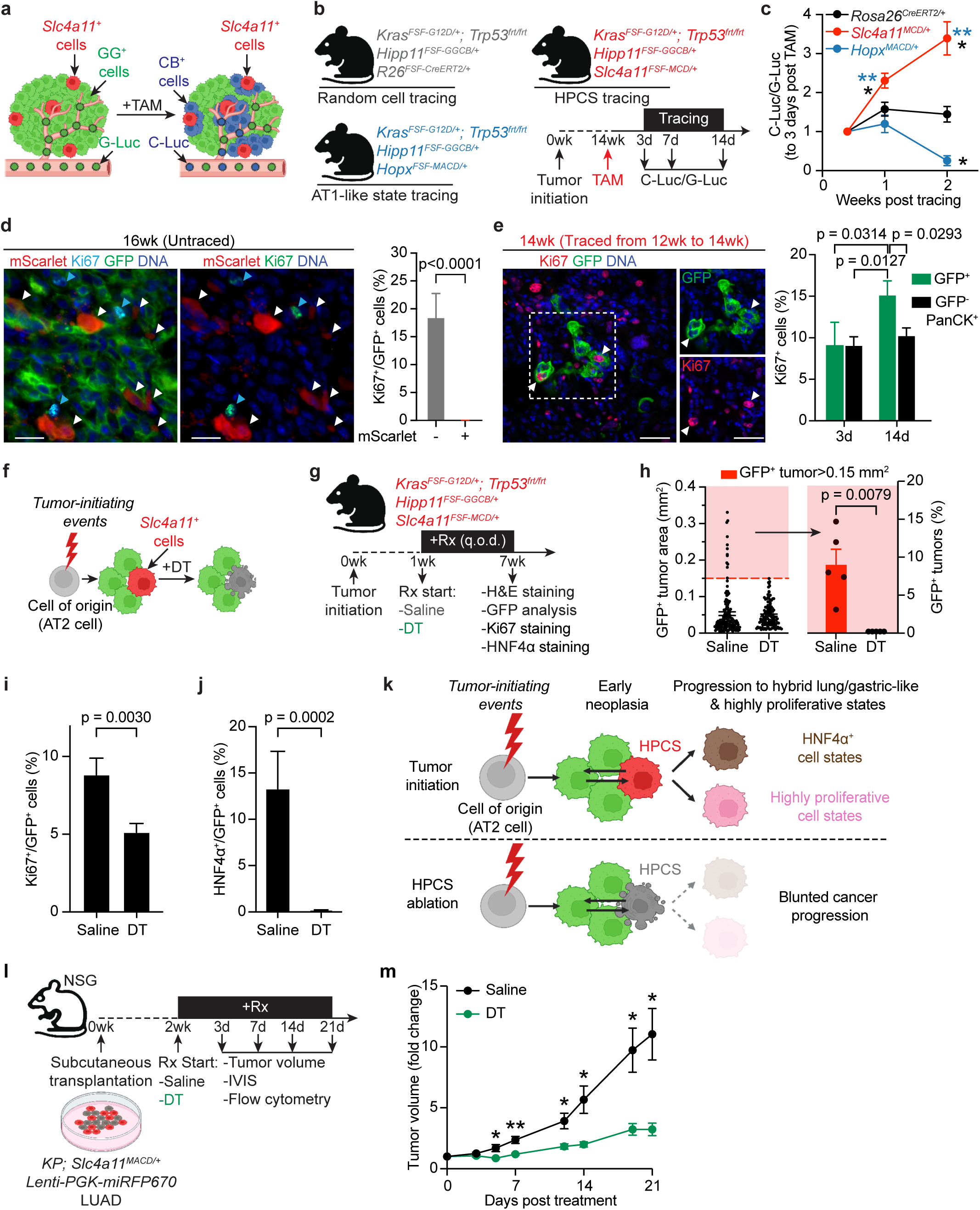
The HPCS drives tumor growth and is essential for LUAD maintenance. **a**, LUAD with *Hipp11^GGCB/+^; Slc4a11^MCD/+^* reporter before (*left*) and after (*right*) tamoxifen (TAM) treatment. TAM administration induces conversion of *GG* (*Gaussia* luciferase, **<u>G</u>**-Luc and E**<u>G</u>**FP) to *CB* (*Cypridina* luciferase, **<u>C</u>**-Luc and Tag**<u>B</u>**FP) in *Slc4a11^+^* LUAD cells. The growth potential of the traced cells can be longitudinally monitored by detection of the relative ratios of G-Luc and C-Luc in the blood of the mice. **b,** Lineage tracing strategy comparing the growth potential of randomly labeled LUAD cells (Random cell tracing, *Rosa26^CreERT2/+^*) vs. the HPCS (HPCS tracing, *Slc4a11^MCD/+^*) and the AT1-like cell state (AT1-like tracing, *Hopx^MACD/+^*) at 14 weeks PTI. Luciferase activity was measured at 3, 7, and 14 days following TAM. **c,** Relative C-Luc/G-Luc activity ratio in the mouse blood normalized to day 3 (3d) across experimental groups: HPCS tracing (*Slc4a11^MCD/+^*, n = 5 mice), random cell tracing (*Rosa26^CreERT2/+^*, n = 4 mice), and AT1-like cell tracing (*Hopx^MACD/+^*, n = 4 mice). Two-way ANOVA. **d,** *Left*: Co-staining of mScarlet^+^ (red), Ki67^+^ (turquoise), and GFP^+^ (green) cancer cells in *KPfrt; Hipp11^GGCB/+^; Slc4a11^MCD/+^* LUAD tumors at 16 weeks PTI (untraced, Fig. 1d). Note that mScarlet (white arrowheads) and Ki67 (blue arrowheads) signals are mutually exclusive. *Right*: Percentage of Ki67^+^cells in mScarlet^-^/GFP^+^ or mScarlet^+^/GFP^+^ LUAD cell subsets. Scale bar: 20 µm. N = 49 tumors from 3 mice. Mann-Whitney U test. **e,** *Left*: Co-staining of Ki67 (red) and GFP (green) in *KPfrt; Rosa26^mTmG/+^; Slc4a11^MCD/+^* LUAD tumors at 16 weeks PTI (traced from 12 to 14 week, Fig. 2a). Demarcated area shown as individual insets to the right. Scale bar: 20 µm. *Right*: Percentage of Ki67^+^ cells in HPCS-derived (GFP^+^) cells at 3d (n = 17 tumors from 3 mice) or 14d (n = 30 tumors from 3 mice) following lineage tracing. GFP^-^/PanCK^+^ (non-HPCS-derived) cells are shown as a comparator (n = 18 tumors from 3 mice for 3d and n = 27 tumors from 3 mice for 14d time points). Two-way ANOVA. **f-g,** Experimental design to examine the effect of ablating *Slc4a11^+^* HPCS cells in the early neoplastic lesions. *KPfrt; Hipp11^FSF-GGCB/+^; Slc4a11^FSF-MCD/+^* mice were administered saline or DT (25 μg/kg, q.o.d.) starting at 1 week PTI and analyzed after 6 weeks on treatment, at 7 weeks PTI. **h,** *Left*: Tumor size shown as GFP*^+^* tumor area in saline vs. DT groups; saline: n = 147 tumors from 5 mice; DT: n = 128 tumors from 6 mice. *Right*: Percentage of GFP*^+^*tumors >0.15 mm^2^ in cross-sectional area, highlighted in red from the *left*. N = 5 mice per group. Mann-Whitney U test. **i,** Ki67^+^/GFP^+^ cells as a percentage of total GFP^+^ cancer cells in saline vs. DT groups. N = 21 tumors from 3 mice in each group. Mann-Whitney U test. **j,** HNF4α^+^/GFP^+^ cells as a percentage of GFP^+^ cancer cells in saline vs. DT groups. N = 10 tumors from 3 mice in each group. Mann-Whitney U test. **k**, Summary of the effect of *Slc4a11^+^* HPCS cell ablation on subsequent cell states in cancer progression as evaluated by HNF4α and Ki67 staining. **l,** Experimental design to test the effects of ablating *Slc4a11^+^* HPCS cells on tumor growth in subcutaneous *KPfrt; Slc4a11^MACD/+^* LUAD reporter allografts. Two weeks after transplantation, allograft-bearing mice were administered saline or DT (25 μg/kg, q.o.d.). Tumor volume measurement, AkaLuc bioluminescence signal detection, and flow cytometry analysis were performed at indicated time point. IVIS: bioluminescence *in vivo* imaging system. **m**, Allograft volume in mice administered either saline or DT. N = 16 tumors from 8 mice (saline) and n = 30 tumors from 15 mice (DT). Welch’s t test. Error bars are SEM.

Given the high growth potential of the HPCS, we hypothesized that either the HPCS itself or a substantial proportion of its derivative cell states must possess high proliferative capacity. To test this, we first examined the proliferative index of the HPCS using scRNA-seq, Ki67 immunostaining, and EdU/BrdU incorporation. Surprisingly, we found the HPCS is highly quiescent (**Fig. 3d; Extended Data Fig. 12a-e**). In marked contrast, lineage-traced LUAD cells that had exited the HPCS (mScarlet^-^/GFP^+^) showed high proliferative capacity as early as 3 days post-tracing (9.13 ± 2.73% SEM Ki67^+^), which further increased by 14 days (15.11 ± 1.76% SEM Ki67^+^) (**Fig. 3e**). The proliferation rate in the non-HPCS derivatives—i.e., the non-traced cells (GFP^-^/PanCK^+^)—remained constant over the 2-week lineage tracing window (3d: 9.04 ± 1.08% SEM vs 14d: 10.21 ± 0.98% SEM, **Fig. 3e**). In striking contrast, the percentage of Ki67^+^ traced cells at the 14 day timepoint is significantly higher than the percentage of Ki67^+^ untraced cells, indicating that the HPCS gives rise to cells with a higher proliferative capacity than an average non-HPCS cell. This explains the high clonal expansion capacity of the HPCS. The proportion of EdU^+^/BrdU^+^ double-positive cells in the traced fraction at 14 days, compared to either the traced cells at the 3-day timepoint or to non-traced cells, was proportional to the changes in EdU single-positive cells (**Extended Data Fig. 12c-f**). This indicates no significant difference in S phase duration between the HPCS derivatives and the non-HPCS derivatives. (**Extended Data Fig. 12f**). Of note, tamoxifen did not significantly confound our assessment of HPCS cell proliferation (**Extended Data Fig. 12g**). These data indicate that while the HPCS is largely quiescent, a subset of its derivatives rapidly enter the cell cycle upon exit from the HPCS, driving clonal expansion of the HPCS-derived cancer cells and tumor growth *in situ*.

Given that the HPCS emerges early during tumorigenesis^4^ and gives rise to advanced cell states, we hypothesized that it plays a critical role in early tumor progression. To address this hypothesis, we ablated the HPCS in early autochthonous *KPfrt; Hipp11^GGCB/+^; Slc4a11^MCD/+^* lung neoplasias by continuous systemic administration of DT starting at 1 week PTI and continuing to 7 weeks PTI (**Fig. 3f, g**). Consistent with the HPCS emerging after the initiation of early neoplasias, we did not detect a difference in the number of neoplastic nodules in the vehicle vs. DT groups (**Extended Data Fig. 13a**). However, we observed a dramatic reduction in overall neoplastic burden, which was largely driven by a significant decrease in the fraction of large (>0.15 mm^2^) neoplastic lesions in the DT group (**Fig. 3h; Extended Data Fig. 13b, c**). Interestingly, ablation of the HPCS blunted progression of grade 1 atypical adenomatous hyperplasias to grade 2 adenomas (**Extended Data Fig. 13d**) and suppressed the emergence cell states with high proliferation rates (**Fig. 3i**) or expression of HNF4α (**Fig. 3j; Extended Data Fig. 13e**), a marker of progression to a hybrid cell state with lung and gastric-like features that emerges after the HPCS in LUAD progression^43,44^ (**Extended Data Fig. 13f-h**). These results indicate that the HPCS is not required for tumor initiation but is critical for the early progression of lung neoplasias, giving rise to cell states with embryonal gene expression programs and/or a high mitotic index, which define advanced cancers (**Fig. 3k**).

As the HPCS possesses high capacity for transitions and a fraction of its derivative cell states harbor a high capacity for proliferation, we hypothesized it plays a central role in tumor maintenance. To test this, we ablated the HPCS cells in autochthonous *KPfrt; Hipp11^GGCB/+^; Slc4a11^MCD/+^* LUAD tumors at 14 weeks PTI via systemic administration of DT (**Extended Data Fig. 13i, j**). This intervention led to a rapid and pronounced ablation of mScarlet^+^ cells and a decrease in tumor burden at 7 days (**Extended Data Fig. 13k-m**). ScRNA-seq analysis of residual GFP^+^ cancer cells confirmed efficient ablation of *DTR^+^* cells, though we still observed *Slc4a11^+^*cells in residual tumors (**Extended Data Fig. 13n**). This suggested incomplete recombination of the *FSF* cassette in the *Slc4a11^MCD/+^* reporter allele in the residual tumors, allowing cancer cells to escape DT-mediated cytoablation. As such, our results likely underestimate the importance of the HPCS in the maintenance of LUAD tumors. To overcome this limitation, we generated a clonal *KP; Slc4a11^MACD/+^* LUAD cell line with confirmed excision of the *FSF* cassette (**Extended Data Fig. 14a-c**). As in the autochthonous tumors, the *Slc4a11^MACD/+^* reporter faithfully marked the HPCS in subcutaneous allografts (**Extended Data Fig. 14d-f**), enabling lineage tracing (**Extended Data Fig. 14g-i**) and ablation of the HPCS cells *in vivo* (**Extended Data Fig. 14j-l**). Ablation of the HPCS produced a notable anti-tumor response (**Fig. 3l, m**), which was accompanied by a reduction in cell state heterogeneity in the residual tumors (**Extended Data Fig. 14m, n**). In contrast, prolonged cytoablation of the AT1-like cell state in subcutaneous *KPfrt; Hopx^MACD/+^*allografts had no significant effect on tumor growth (**Extended Data Fig. 15a-c**), consistent with our results on longitudinal lineage tracing of the AT1-like state (**Fig. 3c**). Collectively, these results establish the HPCS as critical state for the progression and growth of early lung neoplasias and established adenocarcinomas.

### Drug-resistant cell states originate from the HPCS

Plasticity contributes significantly to failure of targeted and chemotherapies, allowing cancer cells to acquire states that are adapted to therapy^3,45,46^. We recently reported that KRAS oncoprotein-targeted therapy promotes acquisition of an AT1-like state that is resistant to KRAS inhibition^9^. Given its high transition potential, we hypothesized the HPCS may similarly serve as a source of therapy-adapted cell states in lung cancer therapy. To address this hypothesis, we applied our longitudinal dual luciferase lineage tracing system to investigate whether the HPCS gives rise to treatment-resistant cancer cell states and minimal residual disease in autochthonous *KPfrt; Hipp11^GGCB/+^; Slc4a11^MCD/+^* lung tumors in models of two standard-of-care therapies, cisplatin chemotherapy^22^ and allele-specific KRAS(G12D) oncoprotein-targeted therapy (MRTX1133)^9,47,48^ (**Fig. 4a**). Compared to the saline group, both MRTX1133 and cisplatin therapies produced a significant decrease in the non-HPCS cells—i.e. the non-traced cells that retain G-Luc—consistent with a robust therapeutic response to the two drugs (**Extended Data Fig. 16a**). In striking contrast, the HPCS-derived cells marked by C-Luc not only survive, but in fact continue to grow under cisplatin and MRTX1133 therapy (**Extended Data Fig. 16b**). These results indicate that the therapeutic response to either therapy occurs in the non-HPCS cells, whereas the derivatives of the HPCS cells continue to thrive under therapeutic pressure and thus become highly enriched in minimal residual disease (**Fig. 4b**).

**Figure 4.**
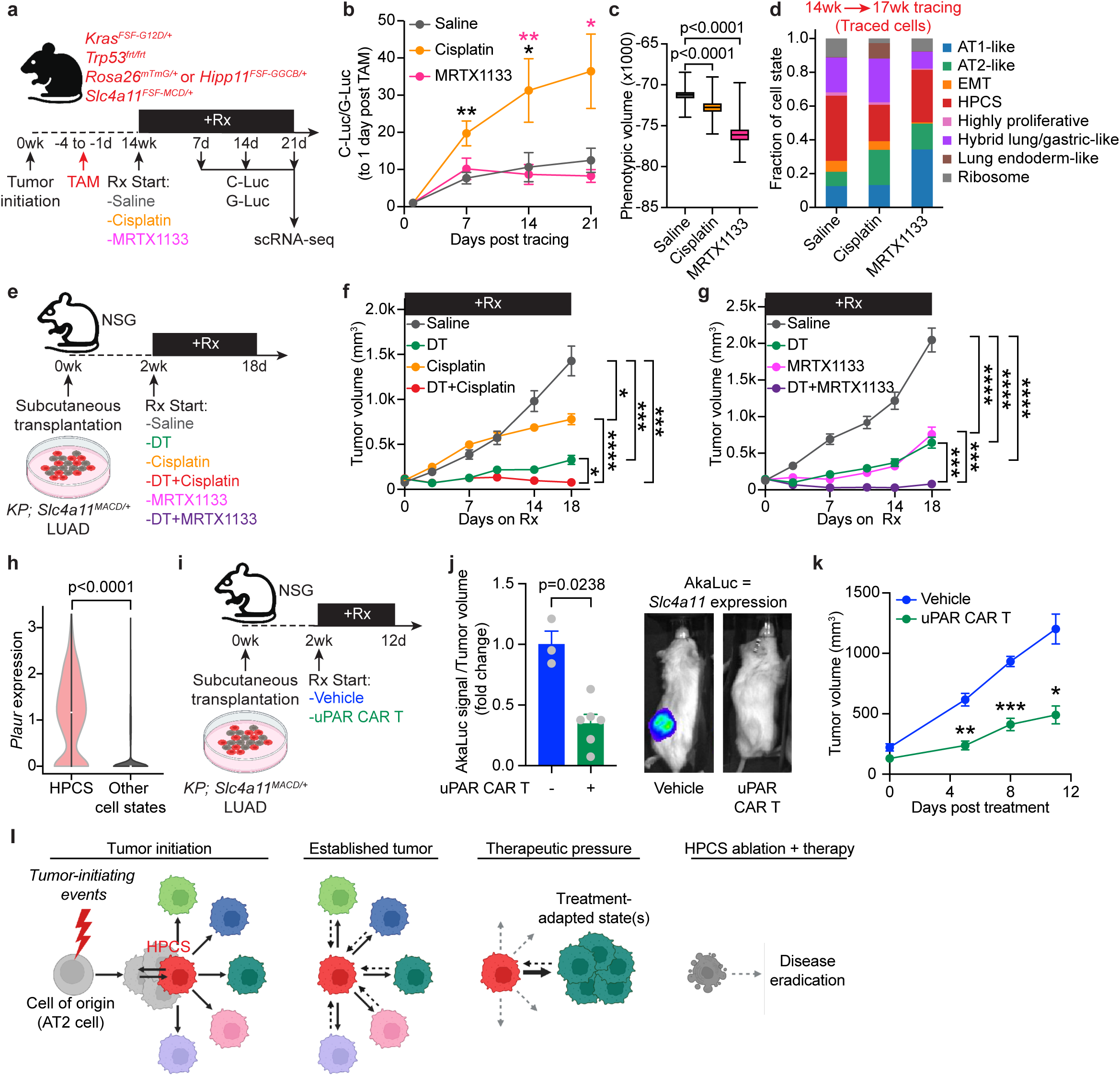
Drug-resistant cell states originate from the HPCS. **a,** Experimental design for assessing impacts of chemotherapy vs. KRAS(G12D)-targeted therapy on HPCS-derived cell states. Mice at 14 weeks PTI were administered one or two doses of tamoxifen (TAM; 200 mg/kg) before initiation of therapy. Mice bearing autochthonous *KPfrt; Hipp11^GGCB/+^; Slc4a11^MCD/+^* lung tumors were administered either saline, cisplatin (1.5 mg/kg, every 3 days), or MRTX1133 (30 mg/kg, b.i.d) for 3 weeks. G-Luc and C-Luc activities were measured at one day after the TAM pulse and subsequently every 7 days on therapy. Traced GFP^+^ LUAD cells [CD45^-^/CD31^-^/CD11b^-^ /CD11c^-^/F4/80^-^/TER-119^-^/Helix NP NIR^-^(Live^+^)/GFP^+^ cells] from autochthonous *KPfrt*; *Rosa26^mTmG/+^*; *Slc4a11^MCD/+^* lung tumors were isolated for scRNA-seq after 21 days on therapy. **b,** C-Luc/G-Luc activity ratio at indicated time points (normalized to day 1) in saline (n = 5 mice), cisplatin (n = 7 mice), or MRTX1133 (n = 8 mice) treatment groups. Welch’s t-test. **c,** Phenotypic volume of cells under the indicated therapies, traced over 3 weeks as in (**a**). N = 4 (saline), 6 (cisplatin), or 7 (MRTX1133) mice. Box and whiskers plotted as min-max. One-way ANOVA. **d,** Cell states of lineage-traced cells under the indicated therapies. **e,** Experimental design to test the effect of ablating *Slc4a11^+^* HPCS LUAD cells by DT (25 μg/kg, q.o.d.) in the context of either saline control, cisplatin chemotherapy (1.5 mg/kg, every 3 days), or KRAS(G12D)-targeted therapy (MRTX1133 30 mg/kg, b.i.d.) in subcutaneous *KPfrt; Slc4a11^MACD/+^* LUAD reporter allografts. **f-g,** Volume of subcutaneous *KPfrt; Slc4a11^MACD/+^* LUAD allografts subjected to the indicated therapies. N = 4-5 mice with 8-10 tumors per group. Mann-Whitney U test. **h,** Violin plot of *Plaur* gene expression in the HPCS vs. other cell states. Mann-Whitney U test. **i,** Experimental design to evaluate uPAR CAR T cells in subcutaneous *KP; Slc4a11^MACD/+^*LUAD reporter allografts. **j,** *Slc4a11*-AkaLuc bioluminescence signal intensity normalized to tumor volume in mice administered either vehicle or uPAR-directed CAR T cells 8 days post injection. N = 3 (vehicle) or 6 (uPAR CAR-T cells) mice. Mann-Whitney U test. **k,** Allograft volume in mice treated with vehicle or uPAR CAR T cells. n = 3 (vehicle) or 6 (uPAR CAR T cells) mice. Mann-Whitney U test. **l,** Schematic summary of key findings (see main text). Error bars are SEM.

We next investigated the fate of the HPCS under therapeutic pressure using scRNA-seq analysis of HPCS-derived lineage-traced cells in *KPfrt; Rosa26^mTmG/+^; Slc4a11^MCD/+^* tumors at 3 weeks on either therapy (**Fig. 4a**). We found both cisplatin and KRAS inhibition suppressed the phenotypic volume, i.e. heterogeneity, of the traced cells (**Fig. 4c**), indicting directed differentiation of the HPCS towards a more limited number of cell states under therapeutic pressure in residual disease when compared to unperturbed tumors. Cisplatin promoted differentiation towards the AT2-like and Hybrid lung/gastric-like fates, whereas KRAS inhibition drove AT1-like differentiation, consistent with our prior work^9^ (**Fig. 4d**). Interestingly, both therapies reduced the fraction of traced cells that remained in the HPCS. These findings suggested cytoablation of the HPCS combined with either chemotherapy or KRAS inhibition may abrogate treatment resistance. Indeed, combining DT-mediated ablation of the HPCS with either cisplatin or MRTX1133 produced a robust combinatorial therapeutic response, nearly eradicating tumors (**Fig. 4e-g**). In contrast, ablation of *Hopx^+^* AT1-like cells—either alone or in combination with cisplatin therapy—did not lead to therapeutic benefit (**Extended Data Fig. 16c, d**). These results indicate the HPCS is a critical transitional node that gives rise to therapy-resistant states and minimal residual disease.

Our findings underscore the therapeutic potential of HPCS cytoablation, motivating us to identify a surface marker that would enable us to direct cytotoxicity specifically to this subset of LUAD cells. We identified robust enrichment of *Plaur,* the receptor for urokinase-type plasminogen activator receptor (uPAR), in the HPCS (**Fig. 4h; Extended Data Fig. 4b; Extended Data Fig. 6b; Extended Data Fig. 16e**). Chimeric antigen receptor T (CAR T) cells targeting uPAR have been developed for the elimination of senescent cells in tissues^49^. We introduced these uPAR CAR T cells^49^ systemically into mice bearing autochthonous *KP; Rosa26^tdTomato/+^*lung tumors and observed eradication of both uPAR^+^ and integrin α2^+^ cancer cells (**Extended Data Fig. 16f-j**), indicative of effective HPCS cytoablation. Excitingly, adoptive transfer of the uPAR-directed CAR T cells into mice bearing subcutaneous *KPfrt; Slc4a11^MCD/+^* LUAD allografts produced a robust anti-tumor response (**Fig. 4i-k**). In sum, our results show the HPCS is a critical, targetable cancer cell state driving tumor progression and maintenance, intra-tumoral heterogeneity, and treatment resistance (**Fig. 4l**).

### An HPCS-like state marked by Slc4a11 and uPAR recurs across carcinomas and regenerating epithelia

Our work establishes the HPCS as a critical cell state driving lung cancer progression and resistance to therapy. Furthermore, our recent pan-cancer analysis of The Cancer Genome Atlas (TCGA) bulk RNA-seq data showed that patients enriched with expression of the HPCS program exhibit worse overall survival^4^. Based on these findings, we hypothesized the HPCS may reflect an essential cancer cell subset across multiple human cancers. Two recent, elegant meta-analyses of scRNA-seq data across a collection of human solid tumors uncovered recurrent cancer cell states, or “archetype states,” that are conserved across all tumors independently of lineage^50,51^. Interestingly, the HPCS signature showed the highest correlation with the *Stress* archetype state in both studies across multiple mouse and human carcinomas, including breast (BC), colorectal (CRC), head and neck (HNC), lung (LUAD), ovarian (OC), pancreas (PDAC), prostate (PCa), and squamous cell cutaneous (SCC) cancers (**Fig. 5a; Extended Data Fig. 17a**). Thus, the HPCS may be functionally important across multiple common human carcinomas.

**Figure 5.**
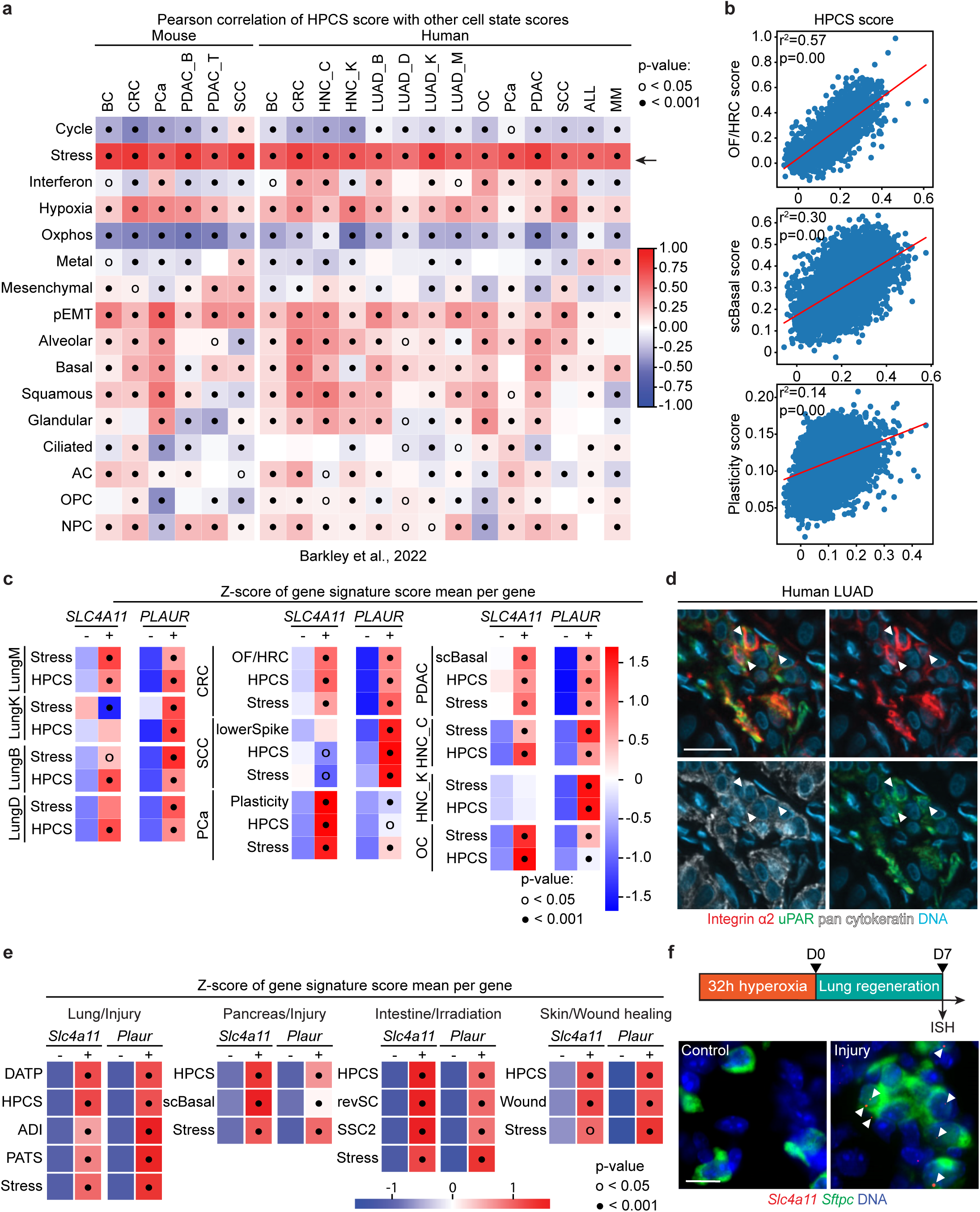
A HPCS-like state marked by *Slc4a11* and *Plaur* is ubiquitously present in multiple carcinomas and regenerating epithelia of other tissues. **a,** Heatmap of Pearson correlations calculated between the HPCS and recurrent pan-cancer cell states (AC: astrocyte-like cell state, OPC: oligodendrocyte progenitor cell-like, NPC: neural progenitor cell-like)^50^, divided by cancer type (BC: breast cancer, CRC: colorectal cancer, HNC: head and neck cancer, LUAD: lung adenocarcinoma, OC: ovarian cancer, PCa: prostate adenocarcinoma, PDAC: pancreatic ductal adenocarcinoma, SCC: squamous cell carcinoma of the skin, ALL: acute lymphoblastic leukemia, MM: multiple myeloma). Arrow indicates the *Stress*-associated cell state from ref. ^50^. A full list of studies is listed in **Supplementary Table 5**. **b,** Pearson correlation between the HPCS signature score (*x*-axis) and plastic cell states (*y*-axis) from either colorectal cancer (OF/HRC: oncofetal/core high relapse^52,54^) (*top*), and pancreatic ductal adenocarcinoma (scBasal^86^) (*middle*), or castration-resistant prostate cancer adenocarcinoma (Plasticity^16^) (*bottom*). Two-tailed p value calculated from the exact distribution of Pearson correlation coefficient. **c**, Heatmaps demonstrating mean gene signature score in either *SLC4A11* or *PLAUR* negative and positive cells scaled using a z-score generated per program to compare gene signatures across different cancer types in human scRNA-seq datasets (**Supplementary Table 5**). For each dataset gene pair, significance is calculated by comparing positive vs negative cells and depicted in the positive cell column. Four human LUAD datasets are shown [LungM^5^; LungK^87^; LungB^88^; LungD^89^] along with plastic cell states [OF/HRC^52,54^, lowerSpike^15^, Plasticity^16^] in other cancers [CRC^90^; SCC^91^; PCa^92^; PDAC^93^; HNC_C^94^, HNC_K^95^, OC^96^]. t-test. **d,** Immunofluorescence of human LUAD stained for (top) uPAR (green) and integrin α2 (red) or (bottom) pan-cytokeratin (white). Arrows point to uPAR and integrin α2 double positive cells. Scale bar: 25 μm. **e,** Heatmaps demonstrating mean gene signature score in either *Slc4a11* or *Plaur* negative and positive cells scaled using a z-score generated per program to compare gene signatures across different injury models in mouse single cell datasets (**Supplementary Table 5**). Heatmaps of signature scores represented as z-scores of the indicated regenerative programs in Lung (DATP: damage associated transient progenitor^55^; ADI: KRT8^+^ alveolar differentiation intermediate^57^; PATS: pre-alveolar type-1 transitional cell state^56^), Pancreas (scBasal cell state^86^), Intestine (revSC: revival stem cell^97^; SSC2: singly profiled stem cell cluster 2^98^) and Skin (Wound^99^). For each dataset gene pair, significance is calculated by comparing positive vs negative cells and depicted in the positive cell column. t-test. **f,** Representative multiplex FISH staining of *Slc4a11* (red) and *Sftpc* (green) in normal lungs vs. 7 days following hyperoxia-induced lung injury. Scale bar: 10 µm.

To further explore the possibility that the HPCS may be important across multiple carcinomas, we compared the HPCS gene expression signature to recently published scRNA-seq datasets identifying candidate plastic states in the context of drug resistance^15,16,52^ or in cancer progression^52,53^ across mouse carcinoma models. We observed significant correlation of the HPCS gene expression scores with the core high-relapse cell state (coreHRC)^52^ or “oncofetal” (OF)^54^ program previously shown to be critical for drug resistance and metastatic recurrence in CRC^16^; with the basal state associated with cancer progression in PDAC^53^; a plastic phenotype described in PCa cells linked to a transition to drug resistant neuroendocrine prostate cancer^16^, and a “lower spike cell state” associating with drug resistance in SCC^15^ (**Fig. 5b, c**). Intriguingly, in human tumors, *SLC4A11*, *PLAUR*, the HPCS signature, and the *Stress* program expression strongly correlated with these subsets of plastic cells proposed to have similar functions as the HPCS (**Fig. 5c**). We uncovered an association of the HPCS and uPAR via integrin α2 and uPAR immunofluorescence *in situ* in human LUAD (**Fig. 5d**). Finally, we identified a significant correlation between *Slc4a11* and *Plaur* expression and a “*KRT8*^+^ alveolar intermediate cell state” (“KAC”), predicted to be a lung adenocarcinoma precursor state in both mouse and human tumors in previous analyses^5^ (**Extended Data Fig. 17b**)—although we note *Krt8* is broadly expressed and not constrained to the HPCS (**Extended Data Fig. 17c**). The convergence of molecular programs in these independently reported carcinoma cell states suggests they may also be functionally convergent across cancer types.

Given the correlation of the HPCS with cellular stress, we speculated *Slc4a11* expression may be associated with regenerative programs in epithelial injury. Prior work has established that a transition state is induced in response to injury-associated stress in AT2 cells and that this response is required for alveolar regeneration^55–57^. We identified significant overlap of the HPCS signature, *Slc4a11*, and *Plaur* expression with this transition state in a scRNA-seq dataset comprising alveolar epithelial cells following lung injury (**Fig. 5e; Extended Data Fig. 18a**). In agreement with this analysis, we detected induction of *Slc4a11* expression in AT2 cells *in situ* in mouse lungs subjected to hyperoxia injury, but not in uninjured lung (**Fig. 5f**). Interestingly, we observed significant enrichment of *Slc4a11, Plaur*, the HPCS signature, and the *Stress* signature in other previously reported injury-associated, regenerative epithelial cell states in the pancreas, intestine, and skin (**Fig 5e; Extended Data Fig. 18b**). Thus, the HPCS may reflect co-option of a latent, stress-associated regenerative program by a subset of cancer cells that physiologically manifests upon epithelial tissue injury.

## Discussion

Using advanced multi-functional genetically engineered autochthonous lung cancer models, we demonstrate that a malignant high-plasticity cell state (HPCS) is critical for cancer progression, maintenance, and resistance to standard-of-care therapies. Functionally, the HPCS harbors high capacity for diverse cell state transitions, including giving rise to highly proliferative states. Thus, the HPCS shares functional features of both normal adult stem cells and CSCs, including robust growth and differentiation potential^58^. The ability of stem-like cancer cells to both self-renew and give rise to differentiated progeny is central to the CSC hypothesis^58^. In contrast to CSCs, we show that the HPCS is *acquired* by neoplastic cells early in tumorigenesis and by other cancer cell states in established adenocarcinomas. Thus, rather than homeostatic tissue stem cells or CSCs, the HPCS is molecularly and functionally similar to a plastic and transient regenerative cell state that is acquired by differentiated AT2 cells upon injury in response to paracrine signals^55–57^. Similar acquired plasticity has been observed in multiple contexts of epithelial regeneration^3,59,60^. Future work will focus on identifying specific niche-derived factors that induce the HPCS and reconstituting them in controlled, reductionist systems to induce the HPCS in non-plastic cancer cells—an approach conceptually similar to reprogramming of differentiated cells to pluripotency using defined factors by Yamanaka and colleagues^61^ or the reconstitution of intestinal stem cell niche factors in organoid culture by Sato, Clevers, and colleagues^62^.

Interestingly, the HPCS is maintained and, in fact, proportionally expands with advancing histopathological grade following its emergence in early neoplasias. Given that the physiological, regenerative correlates of the HPCS are induced by tissue injury lends itself to a model whereby the HPCS becomes induced by niche-derived signals associated with regeneration in a subset of cancer cells, facilitating cancer cell state transitions. In this model, the HPCS forms an intersection or transition hub between cell states—rather than residing at the apex of a hierarchy as postulated by the cancer stem cell hypothesis (**Fig. 4l**). Thus, our findings outline a central principle underpinning the emergence and maintenance of intra-tumoral heterogeneity. These findings are in stark contrast to the low plasticity and growth potential exhibited by the AT1-like state in our experiments. In line with our experimental results, our computational modeling suggests that the

HPCS and the AT1-like state represent the opposite ends of a spectrum of malignant states with variable plasticity^4^. As such, it is possible that other malignant states besides the HPCS harbor relatively high plasticity. Indeed, the Hybrid lung/gastric state was recently shown to harbor co-accessibility at both pulmonary and gastric genes^43,44^, suggesting potential for differentiation towards both lung- or gastric-like downstream states. Future studies utilizing similar lineage tracing and ablation approaches as we report here are needed to functionally interrogate these other cancer cell states *in situ* in lung tumors.

We demonstrate the HPCS is a steppingstone to expansion of intra-tumoral heterogeneity in early-stage tumorigenesis and critical to LUAD progression beyond small nodules. Thus, targeting the HPCS in early lung neoplasias may enable early interception of lung cancer and, possibly, other carcinomas. These findings are consistent with our previous work reporting the HPCS is acquired early in mouse LUAD progression in a subset of early neoplastic cells^4^. Moreover, processes that induce proinflammatory signals required for acquisition of plasticity in wild-type AT2 cells, such as lung injury and exposure to air pollution, promote tumorigenesis^55,63,64^. As such, targeting the HPCS may be particularly effective in preventing lung cancer in patients subjected to such exposures.

Importantly, the HPCS continues to play a central role in established tumors, as evidenced by tumor regression and arrested growth upon HPCS ablation. Our longitudinal lineage tracing results indicate the HPCS-derived cancer cells harbor significantly higher growth potential compared to LUAD cells on average and, thus, ablating the HPCS eliminates a central growth driver in the lung tumors. Furthermore, we found the HPCS enables transitions between cancer cell states, and it can be acquired by non-HPCS cells. As such, during continuous cytoablation, any cell that acquires the HPCS will be killed, contributing to the anti-tumor effect. We note that the rate of non-HPCS cell conversion to HPCS may be accelerated in response to HPCS cytoablation via vacation of HPCS niches in tumors that then become available to non-HPCS cells, as shown previously in the context of ablation of stem-like colon cancer cells^65^. A third possible mechanism involves a non-cell autonomous role for the HPCS in the cellular ecosystem of the tumors, such as immunosuppression or production of paracrine signals that drive cancer cell survival and growth. Indeed, a number of notable growth factors and cytokines with established roles in cancer progression are either specific to or strongly enriched in the HPCS (e.g. *Tnf*, *Il23*, *Vegf*, *Cxcl2*, *Wnt7a*, *Wnt7b*, *Wnt10a*, *Areg*, *Ereg*, and *Tgfa*)^4^. Understanding the contribution of the HPCS to the organization and maintenance of the tumor ecosystem is an important area of future inquiry.

Perhaps the most intriguing explanation for the anti-tumor effect is that eradication of the HPCS as a bridge between defined cell states limits the ability of cancer cells to transition and adapt to the dynamic changes associated with tumor progression, stalling tumor growth. In support of this model, ablation of the HPCS early in LUAD tumorigenesis precluded the emergence of cell states that may be critical to hyperplasia➔adenoma or adenoma➔adenocarcinoma transitions. This model of HPCS as a driver of adaptability is also consistent with the robust synergistic therapeutic effect of combined HPCS ablation and standard-of-care LUAD therapies. These findings suggest the eradication of the HPCS abrogates the ability of treatment-sensitive cancer cell states to transition into resistant states, such as the AT1-like state that is resistant to KRAS inhibition^9^, resulting in a dramatic increase in drug efficacy. Interestingly, both chemotherapy and targeted therapy reduce the fraction of traced cells that remained in the HPCS, suggesting that the HPCS is itself not resistant to therapy or that the therapies reduce the rate at which the HPCS is acquired by the cells that survive in the minimal residual disease.

Our findings elucidate the HPCS as a therapeutic entry point to target therapy resistance enabled by cancer plasticity, encouraging a shift in the current clinical paradigm whereby oncologic therapies invariably result in a drug-tolerant minimal residual disease. We find the HPCS is a significant source of residual disease, especially in the context of chemotherapy, casting the HPCS as a prime target for limiting or even eradicating residual disease (**Fig. 4l**). uPAR-directed CAR T cell therapy targeting the HPCS in LUAD and, potentially, other carcinomas is one possible avenue for the clinical translation of this therapeutic concept. The similarity of the *Stress*-associated cell state that recurs across human solid tumors to the HPCS and the marker genes *SLC4A11* and *PLAUR*, suggests plasticity is concentrated in the *Stress*-associated state in other cancers. Importantly, we found the HPCS program only emerges in epithelia during tissue regeneration and is not present in homeostatic conditions, suggesting cytoablation of the HPCS or targeting the molecular drivers of the HPCS may be safe in cancer patients. Taken together, targeting the HPCS is a promising strategy for suppressing progression and eradicating therapy resistance across carcinomas.

## Methods

### Mice

Previously published genetically engineered mouse strains were used in this study: *Kras^LSL-G12D/+^* (ref. ^19^), *Trp53^flox/flox^* (ref. ^66^), *Kras^FSF-G12D/+^* (ref. ^21^), *Trp53^Frt/Frt^* (ref. ^29^), *Rosa26^mTmG/+^* (ref. ^67^), and *Rosa26^FSF-CreERT2/+^* (ref. ^68^). *Hipp11^FSF-GGCB^*, *Slc4a11^FSF-MCD^*, and *Hopx^FSF-MACD^* reporters were generated in this study as described in detail below. All mice bearing autochthonous *KP* lung tumors were maintained in a C57BL/6 x Sv129 mixed background. NOD.Cg- *Prkdc^scid^ Il2rg^tm1Wjl^*/SzJ (aka NSG mice)^69^ (The Jackson Laboratory, #005557) were used as recipients in all allotransplant studies. All mice were monitored by the investigators and veterinary staff at the Research Animal Resource Center at MSKCC with food and water provided *ad libitum*.

### Autochthonous and transplantation models of lung cancer

Autochthonous LUAD tumors were induced in *Kras^LSL-G12D/+^*; *Trp53^flox/flox^* or *Kras^FSF-G12D/+^*; *Trp53^frt/frt^ (KPfrt*) mice with 1×10^8^-1×10^9^ plaque-forming units (PFU) of AdSPC-Cre, AdSPC-FlpO (Iowa Viral Vector Core), or lentiviral FlpO at 3×10^5^ or 6×10^5^ transforming units, as previously described^70^, in mice that were between 8-12 weeks of age. Immunocompromised NSG mice were used as recipients for either subcutaneous, orthotopic, or intravenous transplantation of *KP* LUAD cell line allografts. For subcutaneous transplantation, cells were resuspended in S-MEM (Gibco, #11380-037) and mixed with Matrigel (Fisher Scientific, #CB-40230C) at a 1:1 ratio. 250,000 cells were implanted subcutaneously into both flanks of NSG mice. For orthotopic transplantation, sorted cells were resuspended in PBS (Gibco, #10010-023) and intratracheally administrated to NSG mice. For intravenous transplantation, 200,000 cells were resuspended in S-MEM and injected into NSG mice via the tail vein. All cell lines were continuously monitored for mycoplasma contamination. Approximately equal numbers of male and female mice were included in all experimental groups in all mouse experiments. Mice were treated in accordance with all relevant institutional and national guidelines and regulations, and mice were euthanized by CO_2_ asphyxiation, followed by intracardiac perfusion with S-MEM to clear tissues of blood when appropriate. A complete list of mice along with age, sex, and age of tumor used in experiments is available (**Supplementary Table 4**). All animal studies were approved by the Memorial Sloan Kettering Cancer Center (MSKCC) Institutional Animal Care and Use Committee (protocol # 17-11-008).

### Generation of donor vectors for embryonic stem cell targeting

For the generation of *Slc4a11-FSF-MCD* donor vector, homology arms ∼1200 bp in length 5’ and 3’ to the end of *Slc4a11* exon 21 (**Extended Data Fig. 1b**) were amplified from genomic DNA of C57BL/6 mouse embryonic stem cell (mESC) using high-fidelity PCR (NEB, #M0494). A homology-directed repair template donor vector was constructed by flanking *frt-bGlobinpA-(PGK-Hygromycin-pA)i-frt-P2A-mScarlet-T2A-CreERT2-P2A-DTR-WPRE-bGHpA* cassette with the 5’ and 3’ homology arms and cloned into the pUC19 plasmid backbone (Takara Bio Inc, #638949) using Gibson Assembly (NEB, #E2611).

For the generation of *Hipp11-FSF-GGCB* donor vector, homology arms ∼5000 bp in length 5’ and 3’ to the safe harbor of *Hipp11* intergenic region (positioned between *Eif4enif1* and *Drg1* genes, **Extended Data Fig. 2a**) were amplified from genomic DNA of C57BL/6 mouse embryonic stem cell (mESC) using high-fidelity PCR. A homology-directed repair template donor vector was constructed by flanking *CAG-loxP-frt-Neomycin-PGKpA-SV40pA-frt-G-Luc-P2A-mEGFP-bGlobinpA-loxP-C-Luc-E2A-TagBFP-3xFlag-WPRE* cassette with the 5’ and 3’ homology arms and cloned into the pUC19 plasmid backbone using Gibson Assembly.

For the generation of *Hopx-FSF-MACD* donor vector, homology arms ∼1500 bp in length 5’ and 3’ to the end of *Hopx* exon 3 (**Extended Data Fig. 9a**) were amplified from genomic DNA of C57BL/6 mouse embryonic stem cell (mESC) using high-fidelity PCR. A homology-directed repair template donor vector was constructed by flanking *frt-bGlobinpA-(PGK-Hygromycin-pA)i-frt-P2A-mScarlet-AkaLuc-T2A-CreERT2-P2A-DTR-WPRE-bGHpA* cassette with the 5’ and 3’ homology arms and cloned into the pUC19 plasmid backbone using Gibson Assembly.

### Embryonic stem cell targeting, genotyping, and chimera generation

*A Kras^FSF-G12D/+^; Trp53^frt/frt^* (*KPfrt*) mESC cell line in the *C57BL/6J* background was generated by crossing a hormone-primed C57BL/6J *Trp53^frt/frt^* female with a *Kras^FSF-G12D/+^; Trp53^frt/frt^* male. At 3.5 days after coitum, blastocysts were flushed out from the pregnant uterus, isolated, and cultured on mouse embryonic fibroblast (MEF) feeder layer. Individual ES cell lines were genotyped by PCR detecting *Kras^FSF-G12D/^, Trp53^frt/frt^,* and *Zfy* (Y-chromosome specific) loci.

For the generation of *Slc4a11^FSF-MCD^*^/+^, *Hopx^FSF-MACD/+^*, and *Hipp11^FSF-GGCB^*^/+^ knock-in mESCs, donor vectors (*Slc4a11-FSF-MCD*, *Hopx-FSF-MACD*, or *Hipp11-FSF-GGCB*, respectively) and ribonucleoprotein (RNP) complex containing HiFi Cas9 nuclease (IDT, #1081061) and crRNA:tracrRNA duplex (IDT) were co-transfected into *KPfrt* mESC line by electroporation (Lonza, 4D Nucleofector). Sequences of crRNAs are listed in **Supplementary Table 4.**

*KPfrt* mESCs were thawed 2 days before targeting and media were changed 1 day and 2 hours before electroporation, respectively. Prior to electroporation, sequence specific crRNA and universal tracrRNA were resuspended in IDTE buffer (IDT) at a concentration of 200 µM and then the crRNA:tracrRNA duplex was formed (final concentration: 44 µM) by combining equimolar concentration of crRNA and tracrRNA and annealing at 95 °C for 5 mins (followed by cooling down to room temperature at ramp-rate of 0.1 C/sec). RNP complexes were formed by combining 22 pmol of crRNA:tracrRNA duplex and 22 pmol HiFi Cas9 nuclease and incubating at room temperature for 20 minutes. For each electroporation, 500,000 mESCs depleted of MEFs, 1 µl donor vector (3 µg/µl), 1 µl RNP complex, 2 µl electroporation enhancer (10 µM, IDT), 16.4 µl Nucleofector P3 primary cell solution and 3.6 µl Nucleofector Supplement 1 were combined and loaded into electroporation cuvette. The ESCs were then plated on top of feeder MEFs and 48 hours later the ESCs were selected with either Hygromycin (*Slc4a11-FSF-MCD* and *Hopx-FSF-MACD*, 150 µg/ml) or G418 (*Hipp11-FSF-GGCB*, 400 µg/ml) for 1 week. Resistant clones were manually picked, expanded, and validated by genotyping using the primers listed in **Supplementary Table 4**.

### Generation of genetically engineered reporter mouse strains

Chimeric F0 mice were obtained by injecting genotype-verified mESCs into host embryos at the 8-cell stage and genotyped at 2 weeks of age. F0 mice were crossed into the *Kras^FSF-G12D/+^; Trp53^frt/frt^*background to generate mice appropriate for the given experiments.

### Generation of LUAD reporter and lineage tracing cell lines

The *Hopx^MACD/+^ KP* LUAD reporter cell line was generated as previously described^9^. For the generation of *Slc4a11^MACD/+^ KP* LUAD reporter cell line, a *KP* LUAD cell line derived from a mouse bearing autochthonous *Kras^LSL-G12D/+^; Trp53^flox/flox^* tumors at 24 weeks post tumor initiation (PTI) was generated. The *KP* LUAD cells were then co-transfected with *Slc4a11-FSF-MACD* donor vector together with *U6-sgSlc4a11-EFS-Cas9* vector expressing guide RNA (ACATATGGGGAGGTATGAGC) targeting the last exon of *Slc4a11* at 1:1 ratio using Lipofectamine 3000 (Thermo Fisher Scientific, #L3000015). The transfected cells were selected with Hygromycin (Sigma-Aldrich, #400053) at a concentration of 150 µg/ml for 2 weeks and single cell-derived drug resistant clones were manually picked for expansion and genotyping with following primers (5’ KI_F1 and 5’ KI_R1; 3’ KI_F1 and 3’ KI_R1). The single cell-derived clones were treated with AdCMVFlpO (Iowa Viral Vector Core) at a multiplicity of infection (MOI) of 500 to remove (*frt-bGlobinpA-(PGK-Hygromycin-pA)i-frt*) *“STOP”* cassette. Excision of the *“STOP”* cassette was confirmed by genotyping spanning the left homology arm and mScarlet (recombined; 5’ KI_F1 and 5’ KI_R2) and by flow cytometry analysis detecting mScarlet fluorescence (**Extended Data Fig. 14d**). An additional *Slc4a11-MACD KP* LUAD reporter cell line was generated via *ex vivo* transformation of an AT2 organoid culture intermediate, as before^9^ (see below). The lentiviral lineage tracing vectors (Lenti-EFS-Flex-TagBFP-PGK-EGFP or *PGK-Gluc-miRFP670-EFS-lox-BFP-lox*^9^) (**Extended Data Fig. 14g-i**) were transduced into the *Slc4a11-MACD KP* LUAD reporter cell lines and FACS sorted based on EGFP or miRFP670 fluorescence.

### Dissociation of lung adenocarcinomas and lung tissue

For isolation of normal AT2 cells and autochthonous LUAD cells, mice were euthanized at the indicated time points post tumor induction and were perfused with sterile S-MEM (Gibco, #11380-037) through the right ventricle of the heart. Dissected lungs or microdissected tumors were dissociated with a mixture of Dispase II (Corning, #354235, 0.6 U/ml), Collagenase Type IV (Thermo Fisher Scientific, #17104019; 0.167 U/ml), and DNase I (Stemcell Technologies, #07469; 10 U/ml) in S-MEM solution at 37 °C as previously described^4^ for 1 hour. The dissociated cells were filtered using a 100 µm filter and spun at 1500 rpm for 10 minutes at 4 °C. The supernatant was removed by aspiration and red blood cell lysis was performed using BD Pharm Lyse (BD Biosciences, #555899) for 1 minute on ice. Cells were then washed with sterile media containing 2% heat inactivated (HI)-FBS (Hyclone, #SH30910.03), passed through a 40 µm filter, and pelleted at 300 g for 5 minutes at 4 °C. The supernatant was removed, and live cells were purified using the Akadeum Dead Cell Removal Microbubble kit as per manufacturer instructions (Akadeum Life Sciences, #11510-211). Cells were resuspended in FACS buffer (2% heat-inactivated FBS in PBS) and counted for use in FACS, as described below.

### Flow cytometry analysis and fluorescence-activated cell sorting (FACS)

Cells were prepared as described above, and Fc block (BD Biosciences, #553142) was added on ice for 10 minutes prior to being stained with the appropriate antibody panel (**Supplementary Table 4**). After 20 minutes of staining on ice, cells were washed twice with FACS buffer and pelleted by a 5 minute spin (300 g at 4 °C). Cell pellets were resuspended in PBS with 2% HI-FBS containing DAPI (Sigma Aldrich, #D9542, 1 μg/ml) or Helix NP NIR (Biolegend, #425301, 5 nM) to identify dead cells. Cell sorting was performed at the Flow Cytometry Core Facility at Sloan Kettering Institute/MSKCC, using a BD FACS Aria Sorter. Cells were sorted using the ‘4-way purity’ mode. Cancer cells were sorted as (CD45/CD31/CD11b/CD11c/F4/80/TER-119)^-^/Helix NP NIR^-^ or DAPI^-^ (live) with specific fluorescent positive cell populations indicated in each experiment.

### Alveolar organoid culture and *ex vivo* transformation protocol

FACS-purified AT2 cells [gated as MHCII^+^/EpCAM^+^/Sca1^-^/podoplanin^-^/lineage^-^ (CD45/CD31/CD11b/CD11c/F4/80/Ter-119)^-^/DAPI^-^] from *KPfrt; Hipp11^GGCB/+^* chimeras were transduced by lentivirus (Lenti-PGK-FlpO or Lenti-PGK-FlpO-P2A-CreERT2) at a MOI of 10 by spinfection (600 g, 37 °C, 30 minutes). 4,000 transduced AT2 cells were resuspended with 50,000 primary pulmonary endothelial cells isolated from 4-week-old *Rosa26^mTmG/+^*mice by FACS (CD31^+^/CD45^-^/DAPI^-^) in 50 µl alveolar organoid culture media [Ham’s F-12 (Thermo Fisher Scientific, #11765047), 10% FBS (Hyclone, #SH30910.03), 1% GlutaMax (Thermo Fisher Scientific, #35050061), 1% Pen/Strep (Thermo Fisher Scientific, #15070063), 1% ITS (Millipore Sigma, #I3126) and 1% HEPES (Thermo Fisher Scientific, #15630080)]. The resuspension was then mixed with 50 µl Matrigel (Fisher Scientific, #CB-40230C) and placed in cell culture inserts (Thermo Fisher Scientific, #08-770). Alveolar organoid culture media (500 µl) was added to the reservoir (Thermo Fisher Scientific, #353504) outside the insert and replaced every 3 days. Primary organoids were digested at day 7 for secondary organoid culture with 5 U/ml dispase (Corning, #354235) for 1 hour at 37 °C and re-plating without endothelial cells to select for transformed tumor spheres. 4-hydroxytamoxifen (4-OHT, Sigma, H6278) was added at a concentration of 1 µM at day 10. Supernatants were collected every 3 days for G-Luc and C-Luc measurement starting at day 1. Organoids were imaged with an EVOS M5000 microscope and dissociated for flow cytometry analysis at day 16 (6 days after 4-OHT).

### Tissue processing for immunofluorescence and immunohistochemistry

Mice were euthanized by CO_2_ asphyxiation followed by systemic perfusion with S-MEM (Gibco, #11380-037) or PBS (Gibco, #10010-023) to clear lungs of blood. Tissues were fixed in 10% neutral buffered formalin (Sigma Aldrich, #HT501128) for 24-48 hours at 4 °C and either embedded in paraffin or dehydrated using 30% sucrose for 16-24 hours before embedding in O.C.T. compound at −80 °C (Thermo Fisher Scientific, #23-730-571).

Immunofluorescence imaging was performed on 7 µm formalin-fixed paraffin-embedded (FFPE) sections or cryosections. FFPE sections de-paraffinized and heat-induced antigen retrieval was performed using EDTA antigen retrieval buffer (Sigma Aldrich, #E1161). For cryosections, slides were air-dried for 1 hour at room temperature and fixed by acetone at −20 °C for 10 minutes. Sections were blocked in donkey immunomix [0.2% BSA (Sigma, #810533), 5% donkey serum (Thermo Fisher Scientific, #31874), 0.3% Triton-X (Fisher Scientific, #BP151-100) in PBS (Gibco, #10010-023)] at room temperature for 30 minutes. Incubation of primary antibodies diluted in donkey immunomix was performed at 4 °C overnight. AlexaFluor-conjugated secondary antibodies raised in donkey were used for signal detection (Invitrogen, #A31573, #A78948, #A10037, #A78947). Sections were counterstained with 1 µg/ml DAPI (Sigma Aldrich, #D9542) for 10 minutes and mounted with coverslips using Mowiol mounting reagent (EMD Millipore, #475904). Mounted slides were imaged using the Zeiss Axio Imager Z2 and ZEN 2.3 software or digitally scanned using Mirax Midi-Scanner (Carl Zeiss AG). Image analysis was performed using Fiji software.

Hematoxylin and eosin (H&E) staining was performed using a standard protocol and tumor grades were assigned using an AI-based Aiforia software (NSCLC_v25 algorithm, Aiforia Technologies Plc, Helsinki, Finland), as before^71^. For immunohistochemistry, tissue sections were incubated at 60 °C and deparaffinized in xylene and rehydrated in alcohol. Antigen retrieval was performed in citrate buffer at 120 °C for 10 minutes in a decloaking chamber. Endogenous peroxidase was blocked by 3% hydrogen peroxide and sections were incubated overnight at 4 °C with antibodies to: Cre Recombinase (Cell Signaling Technology, #15036), uPAR (R&D Systems, #AF543), NKX2-1 (Abcam, # ab76013), or HMGA2 (Cell Signaling Technology, #8179). After primary antibody, biotinylated horse secondary antibodies (R.T.U. Vectastain ABC kit; Vector Laboratories) were added. Signal detection was carried out using the 3, 3′ diaminobenzidine (DAB, Dako) chromogen, followed by counter-staining with hematoxylin. Image analysis was performed using Fiji software. Catalog numbers and dilutions of all antibodies are available in **Supplementary Table 4**.

### *In vivo* EdU/BrdU dual labeling and imaging

Following sequential *in vivo* incorporation of EdU (20 mg/kg, IP, 16 hours prior to euthanasia) and BrdU (100 mg/kg, IP, 4 hours prior to euthanasia), lung tissues were harvested, fixed in formalin (Sigma Aldrich, #HT501128), embedded in paraffin, sectioned, and mounted on slides using standard FFPE procedures. Tissue sections were deparaffinized and rehydrated, followed by EdU staining using the Click-iT™ Plus EdU Cell Proliferation Kit (Invitrogen, #C10637). Next, automated multiplex IF was conducted with the Leica Bond BX staining system. Sections were treated with EDTA-based epitope retrieval ER2 solution (Leica, #AR9640) for 20 minutes at 100°C. The primary antibodies against BrdU (Roche, #1170376), GFP (Abcam, #ab13970) and Cre (Biolegend, #908001) were used. Secondary antibodies were incubated followed by nuclear counterstaining with DAPI (Sigma Aldrich, 5 μg/ml). Slides were mounted using Mowiol 4–88 (Calbiochem) prior to imaging. Image analysis was performed using Fiji software.

### In situ hybridization

mRNA *in situ* hybridization was performed on formalin-fixed paraffin-embedded tissues using the manual Advanced Cell Diagnostics RNAscope 2.5 HD Reagent Kit (# 322350) or the RNAscope Multiplex Fluorescent Reagent Kit v2 (#323100) per the manufacturer’s instructions. Antigen retrieval times and protease digestion times were 15 and 20 minutes for mouse LUAD tissues, respectively. Probes for *Slc4a11* and *Sftpc,* as well as Opal dyes from Akoya Biosciences and their dilutions for use with multiplex fluorescence *in situ* hybridization, are listed in **Supplementary Table 4**.

### AkaLuc in vivo bioluminescence imaging

NSG mice bearing subcutaneous transplants of *KP Slc4a11-MACD* reporter cells and mice bearing autochthonous *KPfrt; Hopx^MACD/+^; Hipp11^GGCB/+^* LUAD tumors were intraperitoneally injected with 100 µl of 30 mmol/l AkaLumine-HCl substrate (Sigma Aldrich, #808350) resuspended in PBS and imaged on an IVIS Lumina II (PerkinElmer).

### Plasma sampling and G-Luc/C-Luc measurements

Whole venous blood was harvested by puncturing the submandibular vein, followed by the collection of 100 μL of blood into capillary blood collector vials (Fisher Scientific, #02-675-185). Plasma was separated by centrifugation at >8,000g for 10 minutes at 4 °C. For G-Luc measurement, plasma or cell culture supernatant was diluted 1:10 in PBS and 200 μmol/L *Gaussia* luciferase substrate coelenterazine-h (NanoLight, #3011) was added. For C-Luc measurement, plasma or cell culture supernatant was diluted 1:100 in PBS and 0.617 μmol/L *Cypridina* luciferase substrate vargulin (NanoLight, #305) was added. Luminescence was immediately measured on a BioTek Cytation 1 (Agilent) at room temperature.

### Administration of DT, MRTX1133, and cisplatin

DT (Sigma, #D0564) was first dissolved in sterile water and diluted in sterile saline for intraperitoneal (i.p.) treatment (at 50 µg/kg daily or 25 µg/kg every other day for long-term treatment and combination treatment studies). Cisplatin (West-Ward, #NDC: 0143-9504-01) was dosed at 1.5 mg/kg by IP every 3 days. MRTX1133 (a kind gift from Mirati Pharmaceuticals) in captisol (#HY-17031, MedChem Express) at 30 mg/kg, b.i.d. was given i.p. as previously described^9^.

### Lineage tracing of Slc4a11^+^, Hopx^+^, and random Rosa26^CreERT2/+^ cell state cells

Mice bearing autochthonous lung tumors were administered one or two doses of tamoxifen (200 mg/kg by oral gavage) at indicated time points. For scRNA-seq analysis of *Hopx^+^* cells and for randomly labeling *Rosa26^CreERT2/+^* cancer cells, one dose of tamoxifen (20 mg/kg by oral gavage) were provided. The baseline measurement at 3 days was chosen to account for the conversion of tamoxifen to its active metabolite 4-hydroxytamoxifen (4-OHT), recombination of the lineage-traced cells, and elimination of residual 4-OHT. Tamoxifen was dissolved in corn oil at 20 mg/mL or 2 mg/mL at 60°C for 1 hour, as described previously^9^.

### Generation of lentivirus

HEK293FreeStyle (HEK293FS) cells were transfected with lentiviral transfer plasmids and the second-generation lentiviral packaging plasmid psPAX2 (Addgene, #12260) and the envelope plasmid pMD2.G (Addgene, #12259) via either the TransIT-LT1 kit (Mirus Bio, #MIR 6000) or Lipofectamine 2000 Transfection Reagent (Thermo Fisher Scientific, #11668500). At 24 hours after transfection, media was discarded and replaced with fresh complete media. Viral media was harvested and filtered through 0.45 µm PES filters (Cytiva, #6780-2504) at 48 hours and 72 hours post transfection. All viral media collected was concentrated using an ultracentrifuge with rotor speed set at >130,000 *g* for 2 hours at 4 °C. The supernatant was discarded into bleach and viral pellets were allowed to solubilize overnight at 4 °C. Concentrated virus was gently mixed and aliquoted. Aliquots were immediately placed on dry ice and stored at −80 °C. A fibroblast reporter cell line expressing GFP upon Flp-mediated recombination was used to titer lentivirus^72^.

### Generation of uPAR CAR T cells

Both mouse SFG γ-retroviral m.uPAR-m28z and human SFG γ-retroviral m.uPAR-h28z plasmids were previously described^49^. In the human m.uPAR-h28z CAR, the anti-mouse uPAR scFV is preceded by a human CD8a signal peptide and followed by a CD28 hinge-transmembrane-intracellular domain, a CD3z intracellular signaling domain and is linked to a P2A sequence to simultaneously express truncated LNGFR. In the mouse m.uPAR-m28z CAR, the anti-mouse uPAR scFV is preceded by a mouse CD8a signal peptide and followed by the Myc-tag sequence, mouse CD28 transmembrane and mouse CD3z intracellular domain^73^. Plasmids encoding the SFG γ-retroviral vectors were used to transfect gpg29 fibroblasts (H29) to generate VSV-G pseudotyped retroviral supernatants, which were used to construct stable retrovirus-producing cell lines as previously described^73,74^. To isolate human T cells from peripheral blood, buffy coats from anonymous healthy donors were purchased from the New York Blood Center. Peripheral blood mononuclear cells were isolated by Ficoll-based density gradient centrifugation. T cells were purified using the human Pan T cell isolation kit (Miltenyi Biotec, #130-096-535), stimulated with CD3/CD28 T cell activator Dynabeads (Invitrogen, #11131D) as described^75^, and cultured in X-VIVO 15 (Lonza, #BEBP04-744Q) supplemented with 5% human serum (Gemini Bio-Products, #100-110-100), 5 ng/ml interleukin-7 and 5 ng/ml interleukin-15 (PeproTech, #200-07-10UG and 200-15-10UG, respectively). T cells were counted using an automated cell counter Vi-CELL BLU (Beckman). Forty-eight hours after initiating T cell activation, T cells were transduced with retroviral supernatants by centrifugation on RetroNectin-coated plates (Takara, #T110B). Transduction efficiencies were determined four days later by flow cytometry and CAR T cells were adoptively transferred into mice or used for in vitro experiments. All blood samples were handled following the required ethical and safety procedures. To isolate mouse T cells from peripheral blood, SV-129 and C57BL/6 mixed background mice were euthanized, and spleens collected. After tissue dissection and red blood cell lysis, primary mouse T cells were purified using the mouse Pan T cell Isolation Kit (Miltenyi Biotec, #130-095-130). Purified T cells were cultured in RPMI-1640 (Fisher Scientific, #11-875-119) supplemented with 10% FBS (GeminiBio, #900-108), 10 mM HEPES (Fisher Scientific, #15-630-080), 2 mM l-glutamine (Thermo Scientific, #A2916801), MEM non-essential amino acids 1x (Thermo Fisher Scientific, #11140050), 55 µM β-mercaptoethanol (Thermo Scientific, #A2916801), 1 mM sodium pyruvate (Thermo Scientific, #11360070), 100 IU ml−1 recombinant human IL-2 (Proleukin; Novartis) and mouse anti-CD3/28 Dynabeads (Gibco, #11453D) at a bead:cell ratio of 1:2. T cells were spinoculated with retroviral supernatant collected from Phoenix-ECO cells 24 hours after initial T cell activation as described^73,76^ and used for functional analysis 3-4 days later.

### Administration of uPAR CAR T cells

For uPAR CAR T transfusion into autochthonous *KP* tumor bearing mice, intraperitoneal cyclophosphamide (200 mg/kg, Long Grove Pharmaceutical, #NDC: 81298-8114-1) was administered to control and uPAR CAR T treatment groups. 16 hours later, a total of 2×10^6^ mouse-derived uPAR CAR T cells per mouse were administered via intraperitoneal injection in uPAR CAR T treated mice. Mice were monitored daily and harvested 7 days post uPAR CAR T transfusion as indicated. For human uPAR CAR T transfusion into NSG allografts, between 5 and 7.5×10^6^ human-derived uPAR CAR T cells were administered intratumorally per mouse.

### Hyperoxia lung injury

Alveolar injury was induced in young C57BL/6 mice by a 32 hour exposure to >85% O_2_ in a hyperoxia chamber (BioSpherix), with FiO_2_ concentration maintained at a constant flow of ∼3 L O_2_/minute and monitored by an in-line oxygen analyzer. Mice were euthanized on day 7 after the 32-hour exposure, followed by collection of lungs for histological analysis.

### Processing of cells for droplet based scRNA-seq

Single cell suspensions from LUAD tumors were prepared and stained as above. Samples were multiplexed using the TotalSeq B cell hashing protocol^77^ (Biolegend, **Supplementary Table 4**). Live sorted cells were collected by flow cytometry, washed once with PBS containing 1% bovine serum albumin (BSA) and resuspended to a final concentration of 700–1,300 cells per μl of PBS + 1% BSA and processed by droplet based scRNA-seq as below.

### Single-cell mRNA sequencing

Single cell suspensions were stained with Trypan blue and Countess II Automated Cell Counter (ThermoFisher) was used to assess both cell number and viability. Following QC, the samples were loaded onto Next GEM Chip G (14143, 15123, 15342, 15488, 15600, 15601, 15771) or GEM-X Single Cell Chip (16235, 16318, 16562, 16686, 17402, 17483, 17543) (10X Genomics PN 1000690 & 2000060) and GEM generation, cDNA synthesis, cDNA amplification, and library preparation of ∼4-50,000 cells proceeded using the Chromium Next GEM Single Cell 3’ Kit v3.1 or GEM-X Single Cell 3’ Kit v4 (10X Genomics PN 1000268 & 1000691) according to the manufacturer’s protocol. cDNA amplification included 11-12 cycles, and 78-863 ng of the material was used to prepare sequencing libraries with 8-14 cycles of PCR. Indexed libraries were pooled and sequenced on a NovaSeq 6000 (14143, 16235, 16562) or X (15123, 15342, 15488, 15600, 15601, 15771, 16235, 16318, 16562, 16686, 17402, 17483, 17543) in a PE28/88 run using the NovaSeq 6000 S4 (200 Cycles) or X 10B (100 Cycles) or 25B (300 cycles) Reagent Kit (Illumina). An average of 38 thousand paired reads was generated per cell.

### Cell surface protein feature barcode analysis

Amplification products generated using the methods described above included both cDNA and feature barcodes tagged with cell barcodes and unique molecular identifiers. Smaller feature barcode fragments were separated from longer amplified cDNA using a 0.6X cleanup using aMPure XP beads (Beckman Coulter, #A63882). Libraries were constructed using the 3’ Feature Barcode Kit (10X Genomics PN 1000276) according to the manufacturer’s protocol with 10-12 cycles of PCR. Indexed libraries were pooled and sequenced on a NovaSeq 6000 (14143_B,16562_B) or X (15123_B, 15342_B, 15488_B, 15600_B, 15601_B, 15771_B, 16235_B, 16318_B, 16562_B, 16686_B, 17402_B, 17543_B) in a PE28/88 run using the NovaSeq 6000 S4 (200 Cycles) or X 10B (100 cycles) or 25B (300 cycles) Reagent Kit (Illumina). An average of 359 million paired reads was generated per sample.

#### Computational Analyses

Jupyter notebooks executing the analysis workflow and figure generation are available on GitHub at https://github.com/dbetel/HPCS_LUAD. All generated sequencing data and count matrices are available at the NCBI Gene-Expression Omnibus under accession record <u>GSE277777</u>.

### Processing of scRNA-seq data

FASTQ files of scRNA-seq data generated on the 10X Chromium or ChromiumX platform were processed using the standard CellRanger pipeline (version >=6.2)^78^. Reads were aligned to a custom GRCm38 / mm10 references including the *EGFP*, *TagBFP, tdTomato, Gluc, Cluc, Akaluc, mScarlet, CreERT2,* and *DTR* transgenes. Cell-gene count matrices were analyzed using a combination of published packages and custom scripts centered around the SCANPY / AnnData ecosystem^79^. Single-cell RNA-sequencing datasets from different mouse models and primary patient samples were analyzed separately using similar workflows.

Single-cell RNA-sequencing data were compiled into a combined count matrix. In general, cells with less than 300 UMIs, more than 10-20% mitochondrial UMIs, and low complexity based on the number of detected genes vs. number of UMIs were removed. Where applicable, doublets were filtered by modeling the TotalSeq B hash count distribution as a Bayesian Gaussian mixture model with variational inference. The same method was used to demultiplex the sample into individual hashes. UMI counts were normalized using the default CPM normalization. In the case of non-hashed transplant samples, the R package *scDblFinder*^80^ was used to detect doublets, which were then removed prior to further analysis.

To identify highly variable features, variance stabilizing transformation and dimensionality reduction was performed on normalized, log_2_-transformed count data using principal component analysis. The resulting dimensionality-reduced count matrices were used as input for UMAP-embedding and unsupervised clustering with the Leiden algorithm^81^.

### Cell state classifications

To compare data generated from the droplet based 10x Chromium platforms with our prior work^4^, we first identified the common genes between each new 10x dataset and our previously published single cell dataset, which was generated using the SmartSeq2 method^82^. For each 10x dataset, we trained a multiclass logistic regression model using the scikit-learn *LogisticRegression* class with options *multi_class=’multinomial’* and *solver=’lbfgs’* using our original cluster labels (hereafter referred to as ‘cell state identities’) and gene counts from our prior work^4^, using only the genes in common between both datasets. We then used this model to classify the cells generated in our 10x dataset. To assign cell state identities to each Leiden cluster generated from our SCANPY pipeline above, we took a pluralistic voting approach where the cell states that were the most represented in a Leiden cluster were used as that cluster’s cell state identity. A rare exception to this were those Leiden clusters where a significant proportion of cells were identified as ‘Highly proliferative’ and were subsequently shown to have high *Mki67* expression. These Leiden clusters were assigned a ‘Highly proliferative’ cell state identity. Cell state assignments can be found in the source code available at the GitHub repository above. Of note, based on recent work identifying a Hybrid lung/gastric-like cell state expressing *Nkx2-1* and *Hnf4α*^43,44^, we reannotated cells classified as clusters 8 and 10 in our original work^4^ to comprise the Hybrid lung/gastric-like cell state.

### Marker evaluation for classification of cell states

Presence of marker transcripts were determined on processed cell counts as transcript levels greater than the minimum value detected by scRNA-seq (i.e. non-zero counts). Cell state classifications were determined as above, with the modification that putative transcript markers were removed as input factors from all processing that could modify cell state assignments. We calculated True Positive (TP), False Positive (FP), True Negative (TN), and False Negative (FN) metrics and calculated marker sensitivity, specificity, positive predictive values, and negative predictive value as below:

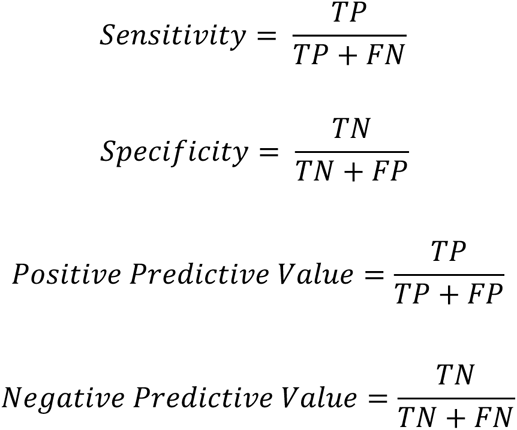

For similar calculations using the SmartSeq2 LUAD data^4^ from which the signatures were originally derived, we reprocessed raw count data using the pipeline described above up to the Leiden clustering step. We then calculated gene scores using the built-in SCANPY *score_genes* function using the top 100 genes from the previously determined HPCS gene signature^4^ (**Supplementary Table 5**), again with the marker genes removed to prevent bias. We labeled the clusters enriched for the HPCS gene signature as the HPCS cluster and calculated the sensitivity, specificity, positive predictive value, and negative predictive values as above for the marker gene of interest. Results for all sensitivity, specificity, positive predictive value, and negative predictive values for marker genes are available in **Supplementary Tables 1-3**. We estimate the sensitivity of the HPCS to be 9.26% and 65.4% based on the 10x droplet-based and SmartSeq2 scRNA-Seq data, respectively, while the specificity is 99.7% and 84.3%.

### Phenotypic volume calculations

Phenotypic volumes, quantitative measures capturing the diversity of cellular phenotypes in cell populations, were performed on highly variable genes, as previously described^40^. Distributions of phenotypic volumes were calculated by sampling 100 cells randomly without replacement from the cells of interest and calculating the phenotypic volume for 1000 replicates per cell population. Statistical significance was determined using either t-tests or ANOVAs to compare distributions of phenotypic volumes.

### Gene signature score calculation and correlation

Gene signatures were compiled from a variety of sources listed in **Supplementary Table 5** and were used to calculate scores via the SCANPY *score_genes* function. Scatter plots and Pearson correlations were generated by independently calculating each cell’s gene signature score and HPCS score and then comparing the distribution of scores. Lines of best fit and r-squared values were calculated using *scipy.stats.linregress* and statistical significance for Pearson correlations was determined via an exact distribution using the built-in *scipy.stats.pearsonr* function. Gene signature scores were used to compare cells with or without *Slc4a11* expression as indicated.

### External data analysis

All gene signatures and their corresponding publications used in this publication are listed in **Supplementary Table S5.**

### Time series analysis

SmartSeq2 scRNA-seq LUAD data (GSE152607)^4^ was downloaded from the NCBI Gene-Expression Omnibus. scRNA-seq data from wild-type AT2 cells, KP adenoma, and KP adenocarcinoma tumor data from 2, 12, 20, and 30 week post tumor induction timepoints were used with Moscot’s^83^ *TemporalProblem* to determine inter-timepoint couplings via optimal transport, and the *RealTimeKernel.from_moscot()* function was used to convert the inter-timepoint couplings into a CellRank^84^ transition matrix. This matrix was used with a Generalized Perron Cluster Cluster Analysis (*GPCCA*) estimator to identify terminal macrostates. Gene expression trajectories to the terminal macrostates were plotted with the Cellrank *gene_trends* function using a Generalized Additive Model (*GAMR*) and with the time component determined by Palantir^85^ pseudotime. The relative estimated start and end of the HPCS was modeled in Palantir pseudotime by plotting the trend of a calculated HPCS score and comparing it to known marker genes using the above process. The calculated HPCS score was defined using the top 100 genes of the HPCS gene signature (**Supplementary Table 5**) with the SCANPY *score_genes* function.

### Software Versions

SCANPY (v >=1.9), pingouin (v 0.5.4), gseapy (v1.1.1), numpy (v >=1.26), scipy (v >=1.12), scikit-learn (v >=1.13), leidenalg (v 0.10.2), matplotlib (v 3.8.4), Cellrank (v2.0.7), Palantir (v1.4.1), R (v4.3.3), Fiji (v2.14.0)

### Statistics

Statistical analyses were performed using Student’s t-test, Welch’s test, Mann–Whitney U test, Kruskal Wallis test, one-way ANOVA, or two-way ANOVA, as appropriate. Statistical significance for the figures was indicated as raw p*-*values or was defined as *p<0.05, **p<0.01, ***p<0.001, ****p<0.0001; ns, not significant.

## Acknowledgments

We thank members of the Tammela laboratory for helpful discussions; L. Bombardelli for advice and suggestions regarding the G-Luc and C-Luc secreted luciferases; K. Manova, W. Kang, M. Tipping, and the Molecular Cytology Core for histology support; E. Chan and E. Rosiek for help with image analysis and quantification; E. de Stanchina and the Antitumor Assessment Core Facility for support with drug administration and tumor transplant experiments; R. Gardner, M. Kweens, and A. Longhini for FACS support; N. Mohibullah and the Integrated Genomics Operation for next-generation sequencing support; H. Alcorn and O. Grbovic-Huezo for laboratory management; and M. Blum, S. Ding, A. Hudson, E. Rivas-Hernandez, and B. Christensen for help with experiments. We are grateful to J. Christensen and J. Hallin at Mirati Therapeutics for providing MRTX1133. This work was supported by NIH K08-CA267072, the NIH Loan Repayment Program, and the Linn Fund for Sarcoma Research (to J.E.C.); American Cancer Society award, PF-25-1422234-01-PFCBI, Grant DOI #: https://doi.org/10.53354/ACS.PF-25-1422234-01-PFCBI.pc.gr.230344 (to C-H.P.); a PhD fellowship from Boehringer Ingelheim Fond (to K.K.); a Damon Runyon postdoctoral fellowship (2467-22), a postdoctoral fellowship from the American Federation for Aging Research, and the St. Louis Ovarian Cancer Awareness Research Grant for Ovarian Cancer from the Foundation for Women’s Cancer (to Z.Z.); New York Stem Cell Science NYSTEM training award (C32559GG), the Center for Stem Cell Biology at MSKCC, and support from the Druckenmiller Center for Lung Cancer Research at MSKCC (to X.Z); NIH R01-AG054720 (to D.B.); NIA R01-AG065396, support from the MSK Technology Development Fund (TDF) (FP00009954), the Mark Foundation Grant for Cancer Research, HHMI and the Geoffrey Beene Cancer Research Center (to S.W.L); the Sigrid Juselius Foundation, National Natural Science Foundation 82373443, Fundamental Research Funds for Central University 2662025SYPY005, Huazhong Agricultural University Pilot Project Fund (to Y.Y); NIH R01-CA270116, NIH R01-CA293718, an AACR Next Generation Transformative Award, and Josie Robertson and Rita Allen Scholarships (to T.T.); and by the NIH/NCI Cancer Center Support Grant P30-CA08748 (to MSKCC). We acknowledge the use of the Antitumor Assessment, Integrated Genomics Operation, Flow Cytometry, Molecular Cytology Core, and Single Cell Analytics and Innovation Lab Facilities at the Sloan Kettering Institute, funded by CCSG P30-CA08748, Cycle for Survival, and the Marie-Josée and Henry R. Kravis Center for Molecular Oncology. The schematic illustrations were created using BioRender (https://biorender.com).

## Author contributions

J.E.C, C-H.P., Y.Y., and T.T. conceived and designed the study; J.E.C. and T.T. wrote the manuscript; C-H.P., Y.Y., and D.B. contributed to the writing of the manuscript; J.E.C, C-H.P., Y.Y., D.B., and T.T. interpreted the data; J.E.C, C-H.P., Y.Y., K.K., E.B., Z.Z., H.S., G.G., G.H., Z.L., and X.Z. performed experiments and data analysis; Z.Z. and S.W.L. contributed CAR T cells and associated methodology; J.E.C., J.R., and D.B. performed computational analysis; J.E.C. and D.B. supervised computational analysis and interpreted data; S.W.L. and T.T. supervised experimental work; S.W.L. and T.T. obtained funding. All of the authors approved the final manuscript.

