## Extended Data Figures for "Functional interrogation uncovers a critical role for a high-plasticity cell state in lung adenocarcinoma"

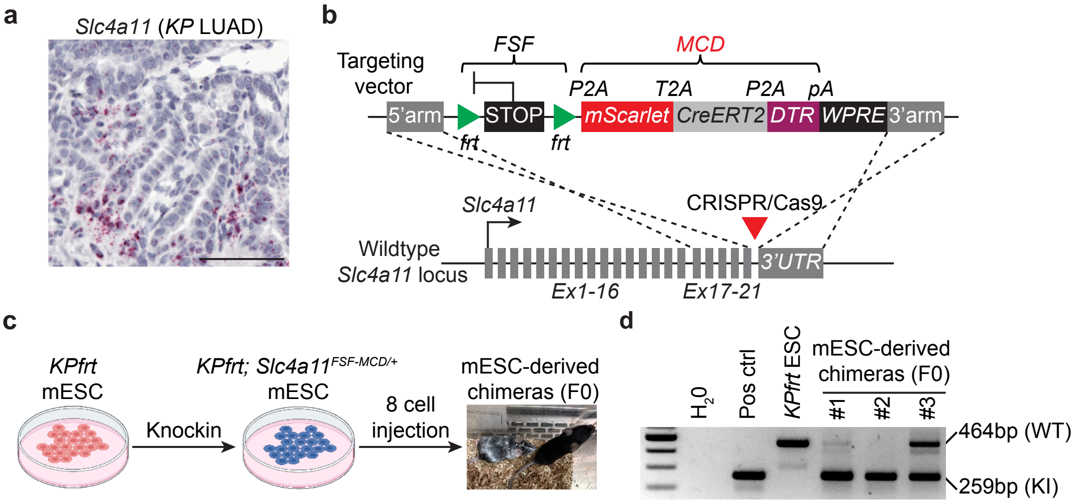




Chan, Pan et al.,
Extended Data Fig. 1

**Extended Data Figure 1. Construction of the *Slc4a11*$^{FSF\text{-}MCD/+}$ reporter allele. a,** Single-molecule mRNA *in situ* hybridization detecting *Slc4a11* expression in *KP* LUAD 20 weeks post tumor initiation (PTI). Scale bar: 50 μm. **b,** Components of the *Slc4a11*$^{FSF\text{-}MCD/+}$ reporter and targeting strategy by CRISPR/Cas9 mediated homology dependent repair (HDR). Strategy for generating genetically engineered *Slc4a11*$^{MCD/+}$ knock-in (KI) reporter enabling lineage tracing and ablation of *Slc4a11*$^{+}$ HPCS in *KP* LUAD: *frt-stop-frt-P2A-mScarlet-T2A-CreERT2-P2A-DTR-WPRE* (*FSF-MCD*) reporter construct was knocked in frame into the stop codon of exon 21 of *Slc4a11* gene in the presence of CRISPR/Cas9 mediated gene editing. **c,** Experimental outline to generate *Kras*$^{FSF\text{-}G12D/+}$*; Trp53*$^{frt/frt}$*; Slc4a11*$^{FSF\text{-}MCD/+}$ reporter mouse. The *FSF-MCD* reporter construct was knocked into a *Kras*$^{FSF\text{-}G12D/+}$*; Trp5*$^{frt/frt}$ (*KPfrt*) mouse embryonic stem cell (mESC) and the correctly targeted mESC clones were microinjected into 8-cell stage embryos to obtain mESC-derived chimeras. **d,** Gel electrophoresis of the PCR products from a correctly targeted mESC clone (pos ctrl), parental *KPfrt* mESC, or mESC-derived chimeras using primer pairs detecting either wild-type (WT, 464 bp, *top*) or knock-in (KI, 259 bp, *bottom*) alleles, respectively.

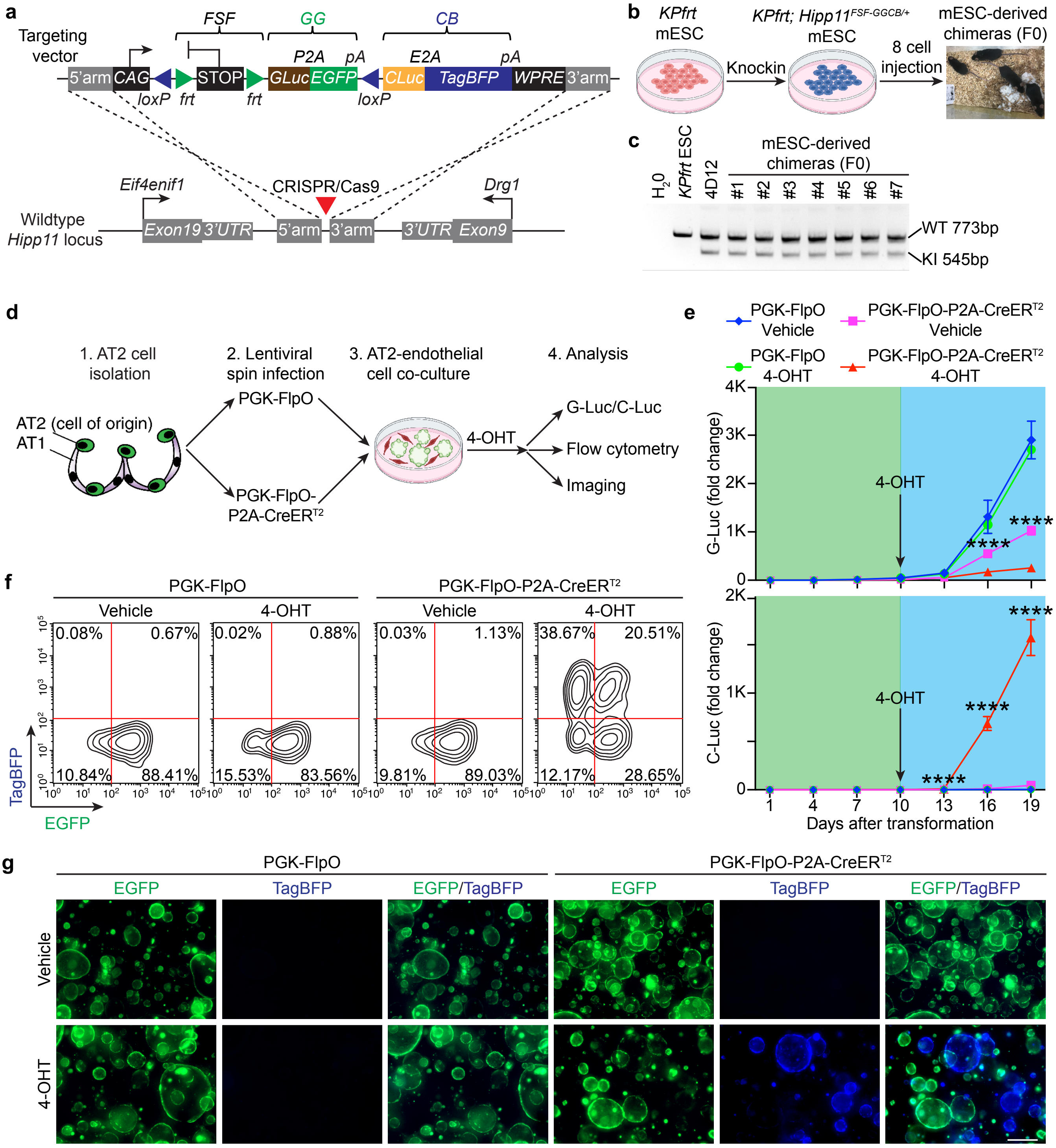


Chan, Pan et al.,
Extended Data Fig. 2

**Extended Data Figure 2. Construction and validation of the *Hipp11*$^{FSF\text{-}GGCB/+}$ reporter *in vitro*. a,** Vector map of *Hipp11*$^{FSF\text{-}GGCB/+}$ targeting vector and strategy for the generation of *Hipp11*$^{FSF\text{-}GGCB/+}$ reporter by CRISPR/Cas9 mediated dependent HDR. *CAG-LoxP-frt-stop-frt-G-Luc-P2A-EGFP-pA-LoxP-C-Luc-E2A-TagBFP-pA-WPRE* (*FSF-GGCB*) reporter construct was knocked into the safe harbor of the *Hipp11* intergenic region (positioned between *Eif4enif1* and *Drg1* genes) in the presence of CRISPR/Cas9 mediated gene editing. FlpO-mediated recombination removes the *FSF* cassette and activates the *G-Luc-P2A-EGFP (GG)* element, *while* Cre-mediated recombination removes the *loxP-stop-loxP* (*LSL*) cassette and activates the *C-Luc-E2A-TagBFP* (*CB*) element. G-Luc: *Gaussia* luciferase; C-Luc: *Cypridia* luciferase. **b,** Experimental outline to generate the *Kras*$^{FSF\text{-}G12D/+}$*; Trp53*$^{frt/frt}$*; Hipp11*$^{FSF\text{-}GGCB/+}$ reporter mouse. *CAG-FSF-GGCB* reporter construct described in panel (**a**) was knocked into *KPfrt* mESCs, the correctly targeted mESC clones were microinjected into 8-cell stage embryos to obtain mESC-derived chimeras. **c,** Gel electrophoresis of the PCR products from either a parental mESC (*KPfrt* mESC), a correctly targeted mESC clone (4D12), or mESC-derived chimeras using primer pairs detecting either wild-type (WT, 773 bp, *top*) or knock-in (KI, 545 bp, *bottom*) alleles. **d,** Experimental outline to validate the *Hipp11*$^{FSF\text{-}GGCB}$ reporter in alveolar type 2 (AT2) cells derived organoids using *ex vivo* transformation assay. AT2 cells isolated from the chimeras were transduced with lentivirus expressing either FlpO or FlpO-P2A-CreERT2 to remove the *FSF* cassette and activate oncogenic *Kras*$^{G12D/+}$ and delete *Trp53*, resulting in the generation of *Kras*$^{G12D/+}$ mutant and *Trp53* deficient LUAD organoids. The transformed organoids were exposed to 4-OHT and analyzed with the indicated approaches. **e**, Longitudinal monitoring of G-Luc (*top*) and C-Luc (*bottom*) activities depicted as fold change over day 0 from *ex vivo* transformed LUAD organoids as in (**d**). Organoids were exposed with 4-OHT (1 μM) at day 10. Media was refreshed every 3 days before supernatant was collected for luciferase activity measurement. N = 6 for each group. Error bars are SEM. Two-way ANOVA. **f**, Flow cytometry analysis of EGFP vs. TagBFP expression from PGK-FlpO (*left*) or PGK-FlpO-P2A-CreERT2 (*right*) transformed organoids 6 days after treatment with either vehicle control or 4-OHT. TagBFP is only induced in organoids harboring $CreER^{T2}$ and exposed to 4-OHT. **g**, Fluorescent images of EGFP and TagBFP from LUAD organoids transformed with indicated lentivirus 6 days after treatment with either vehicle control or 4-OHT. Consistent with (**f**), TagBFP is only present in organoids harboring $CreER^{T2}$ and exposed to 4-OHT. Scale bar: 200 μm.

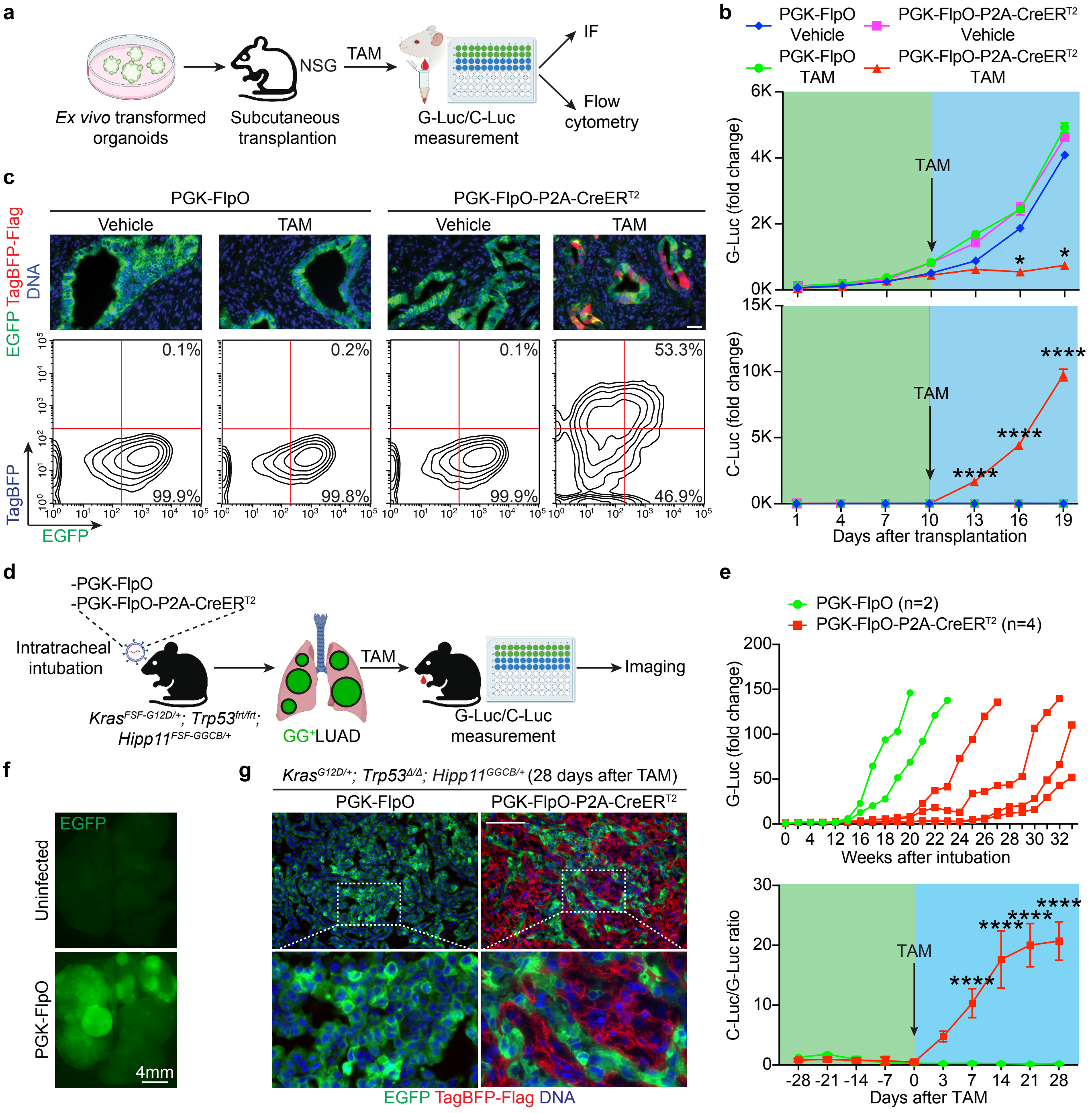




Chan, Pan et al.,
Extended Data Fig. 3

**Extended Data Figure 3. Validation of the *Hipp11*$^{GGCB/+}$ reporter *in vivo*. a,** Experimental outline to validate the *Hipp11*$^{FSF\text{-}GGCB/+}$ reporter in subcutaneous transplanted LUAD tumors derived from *ex vivo* transformed organoids. *Ex vivo* transformed tumor organoids were transplanted into the subcutaneous flank of NSG mice and administered with tamoxifen (TAM, 200 mg/kg) at day 10 post transplantation. Mice were cheek bled every 3 days for G-Luc and C-Luc measurements. Tumors were harvested at day 19 for flow cytometry analysis and IF staining. **b**, Longitudinal monitoring of G-Luc (*top*) and C-Luc (*bottom*) activity (shown as fold changes relative to day 1) at the indicated time points from the indicated groups of tumor-bearing mice. N = 2. Two-way ANOVA. **c,** IF images (*top*) and flow cytometry plots (*bottom*) for the analysis of EGFP and TagBFP expression from transplanted tumors at indicated conditions. TagBFP IF was performed using anti-Flag antibody detecting the TagBFP-3xFlag fusion protein. Scale bar: 20 μm. **d,** Experimental outline to validate the *Hipp11*$^{FSF\text{-}GGCB/+}$ reporter in autochthonous LUAD tumors. *Kras*$^{FSF\text{-}G12D/+}$; *Trp53*$^{frt/frt}$; *Hipp11*$^{FSF\text{-}GGCB/+}$ mice were intubated with lentivirus expressing either PGK-FlpO or PGK-FlpO-P2A-CreER$^{T2}$ to initiate LUAD and were cheek bled every other week to monitor for tumor formation. Mice from both groups were treated with a single dose of TAM (200 mg/kg) when the G-Luc level reached a 10-fold increase compared to non-infected control mice. Mice were cheek bled every 3 days after TAM treatment to assess the C-Luc/G-Luc ratio. **e,** Longitudinal monitoring of G-Luc activity (*top*) or C-Luc/G-Luc ratios (*bottom*) from mice intubated with lentivirus expressing either PGK-FlpO or PGK-FlpO-P2A-CreER$^{T2}$. G-Luc and C-Luc activities were measured and shown as fold change over non-infected control mice. N = 2 for PGK-FlpO; n = 4 for PGK-FlpO-P2A-CreER$^{T2}$. Two-way ANOVA. **f,** Dissection microscope images depicting EGFP signal in uninfected (*top*) vs PGK-FlpO (*bottom*) infected *KP*; *Hipp11*$^{GGCB/+}$ mice. **g,** IF staining of tumors from (**d**) stained with antibodies raised against EGFP (green) and Flag (TagBFP-Flag, red). Scale bar: 200 µm. Error bars are SEM.

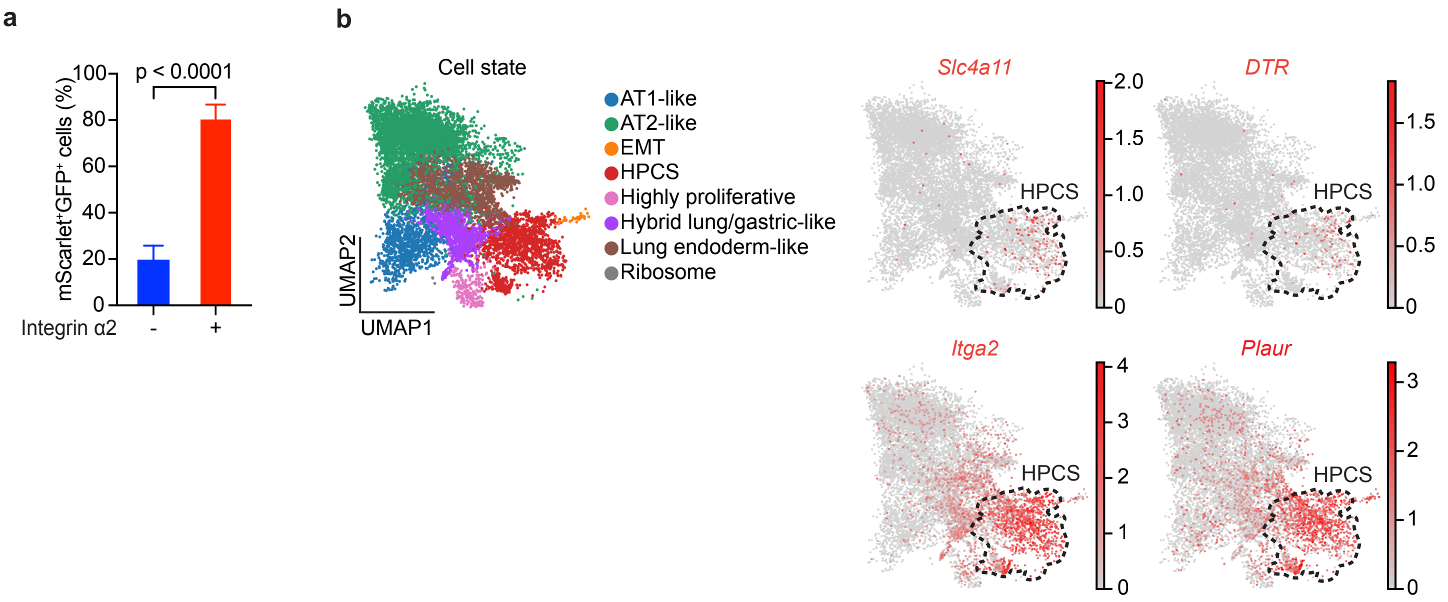




Chan, Pan et al.,
Extended Data Fig. 4

**Extended Data Figure 4. Validation of the *$Slc4a11^{MCD/+}$* allele *in vivo.* a,** Percentage of mScarlet$^+$/GFP$^+$ cells staining for integrin α2 measured by IF imaging. N = 19 tumors from 3 mice. Error bars are SEM. Welch's t-test. **b,** Plots of eight annotated LUAD cell states that molecularly define LUAD tumors collected from six *KP* mice 15-16 weeks PTI. Distribution of gene expressions for *Slc4a11* (*upper left*)*, DTR* (*upper right*), *Itga2* (*lower right*), and *Plaur* (*lower right*).

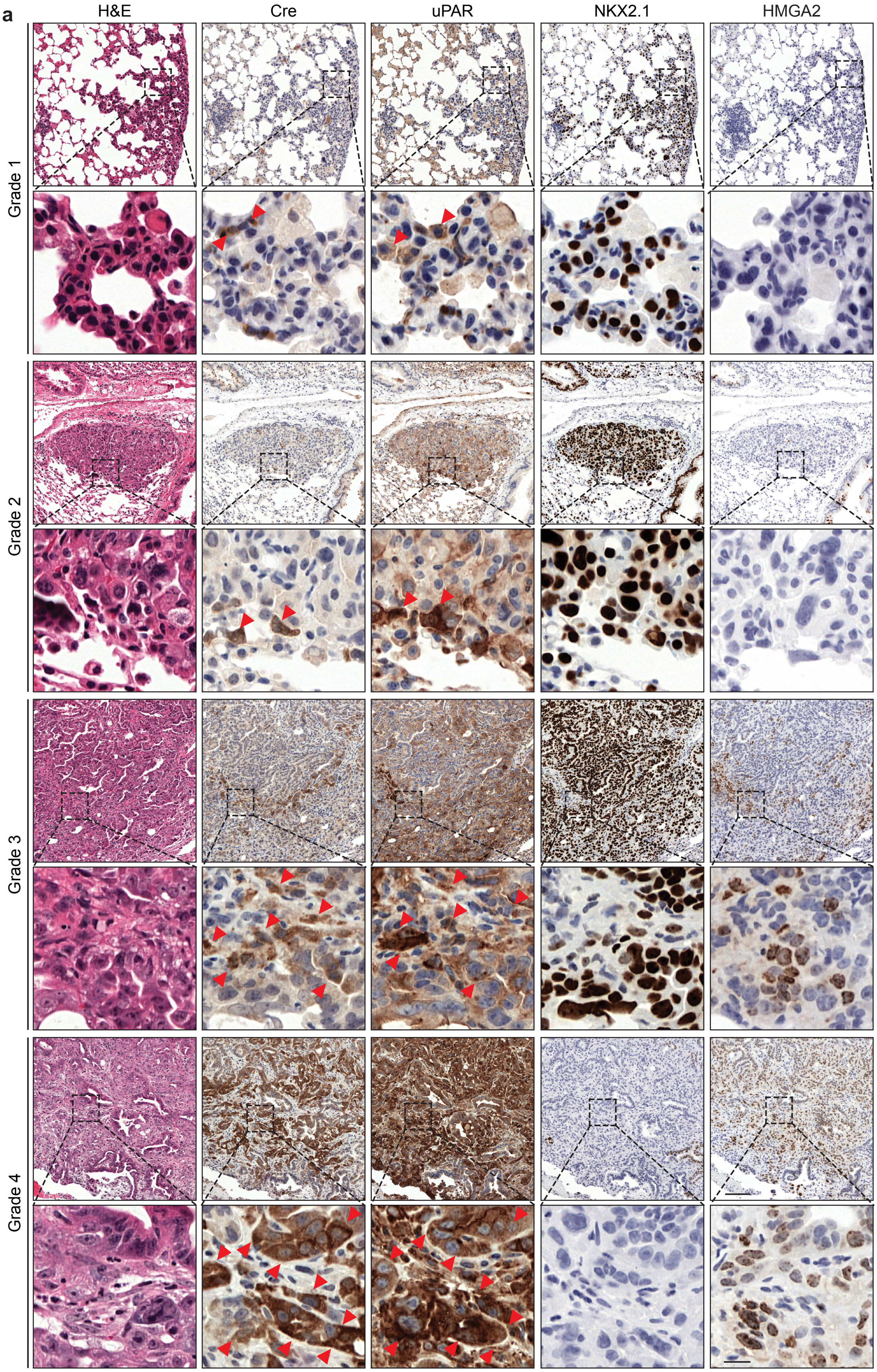


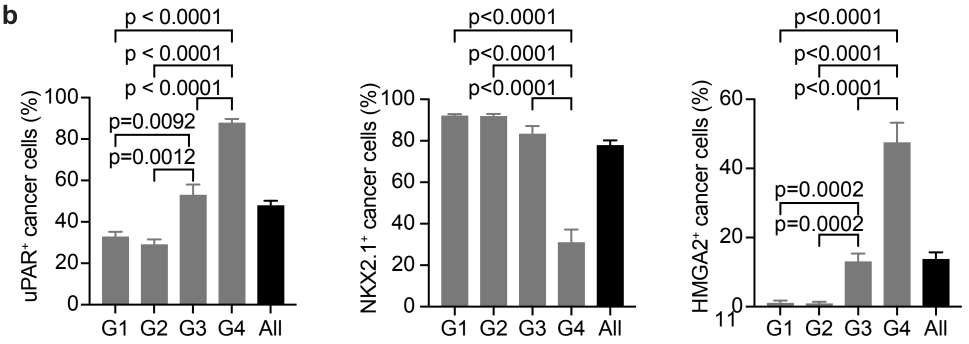


Chan, Pan et al.,
Extended Data Fig. 5

**Extended Data Figure 5. Immunohistochemistry staining of Cre (marking the HPCS), uPAR, NKX2.1, and HMGA2 across tumors of distinct histopathological grades. a,** Representative H&E and immunohistochemistry (IHC) staining of Cre, uPAR, NKX2.1 and HMGA2 on serial sections from autochthonous *KPfrt; Hipp11*$^{GGCB/+}$*; Slc4a11*$^{FSF\text{-}MCD/+}$ LUAD tumors at 7 and 16 weeks PTI. Histopathological grades were assigned using artificial intelligence (AI)-based image analysis software provided by Aiforia. Scale bar: 100 µm (low magnification) and 10 µm (high magnification). **b,** uPAR$^{+}$ (*left*), NKX2.1$^{+}$ (*middle*), HMGA2$^{+}$ (*right*) tumor cell percentages across histopathological grades. N = 28-34 tumors from 4 mice per grade. One-way ANOVA. Error bars are SEM.

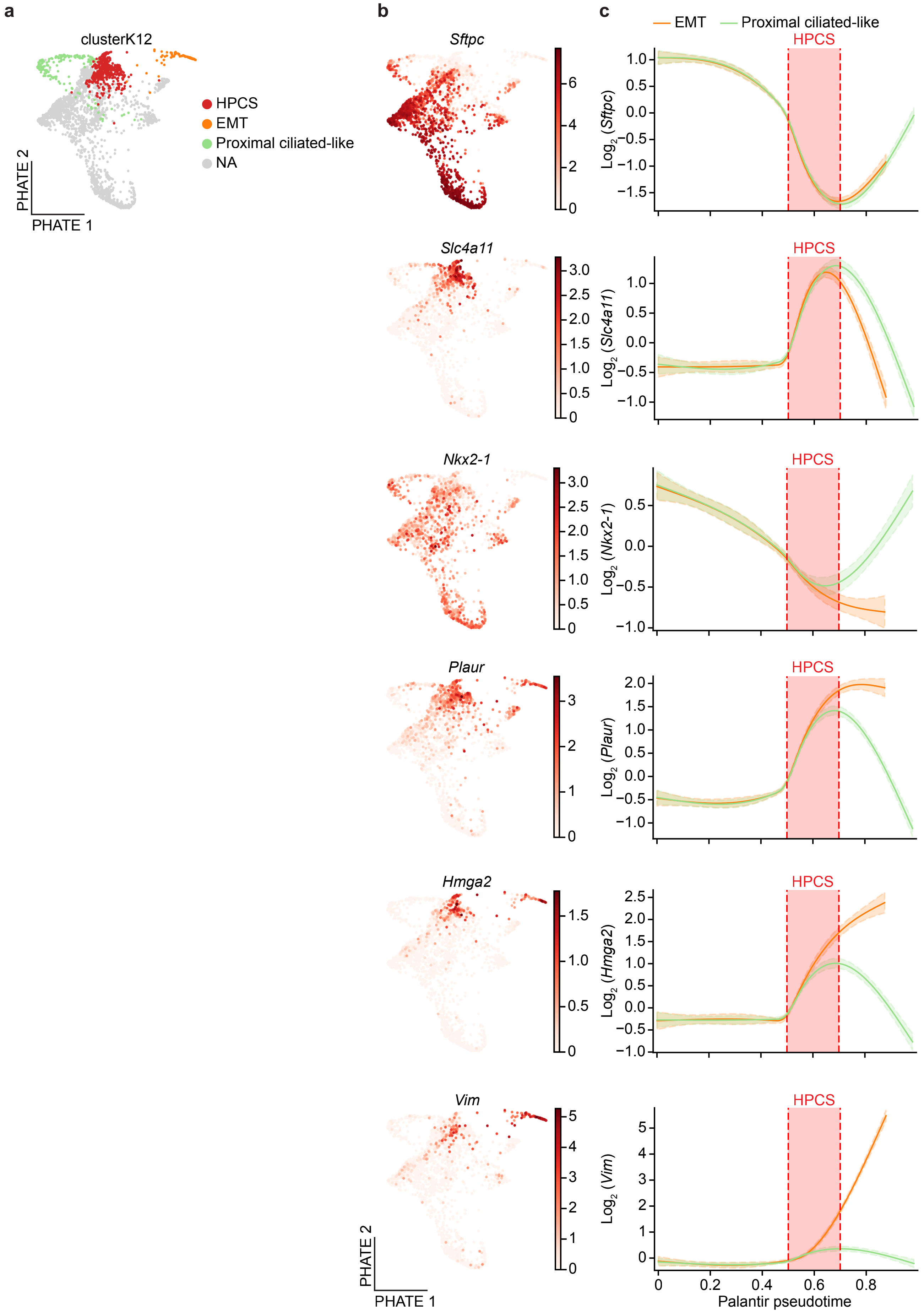


Chan, Pan et al.,
Extended Data Fig. 6

**Extended Data Figure 6. Gene expression trends in longitudinal scRNA-seq LUAD data. a,** Predicted end cell states (EMT, Proximal ciliated-like) and the HPCS projected on the original PHATE[100] map from Marjanovic*, Hofree*, Chan* et. al.[4]. **b,** Gene expression of the indicated genes plotted as PHATE maps. **c**, Gene expression trajectories for the indicated marker genes plotted over Palantir[85] *pseudotime*, with cell state trajectories predicted by CellRank[84], and estimated time spent in HPCS cell state shaded in red (see '*Time Series Analyses'* in **Methods**).

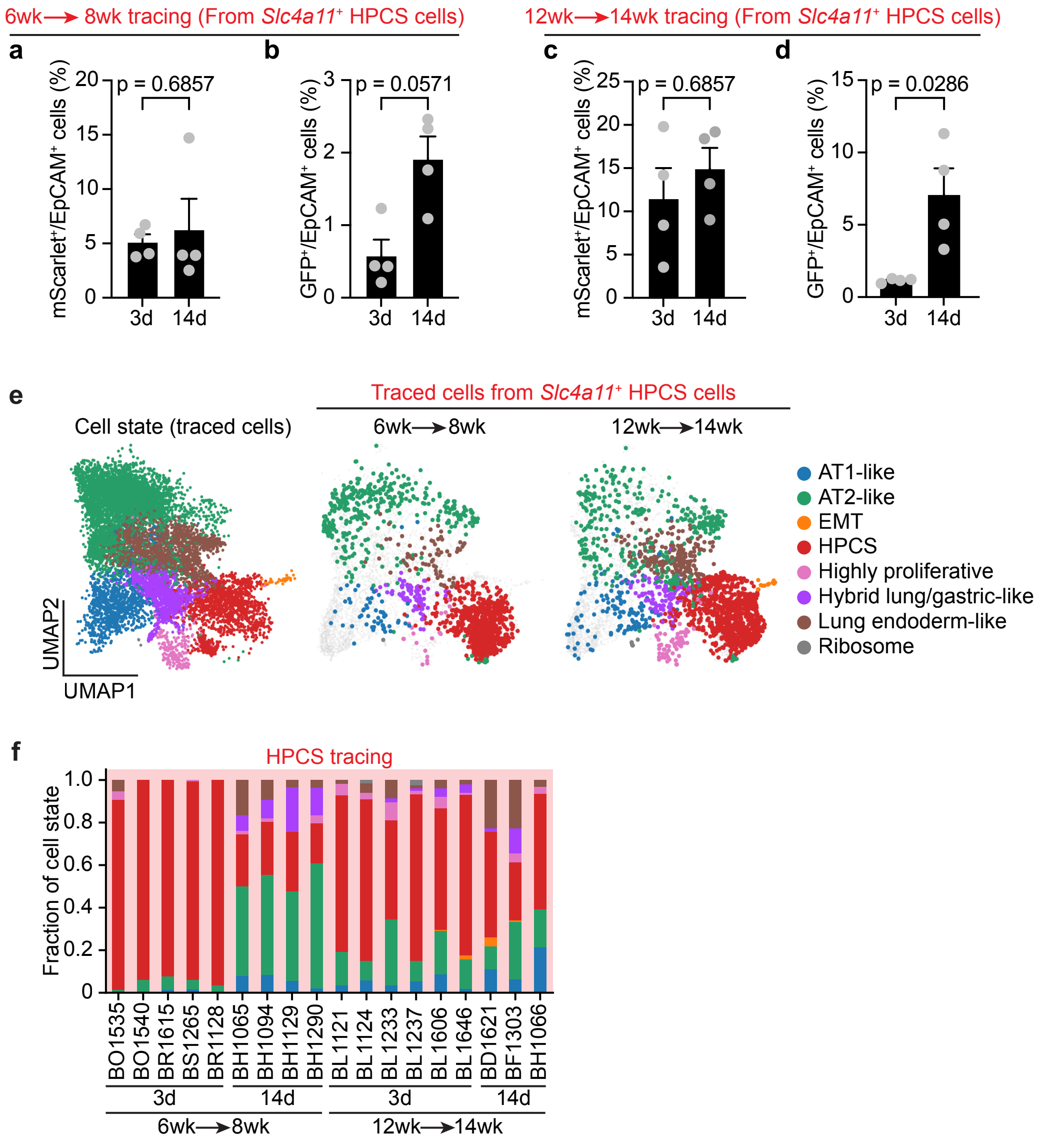




Chan, Pan et al.,
Extended Data Fig. 7

**Extended Data Figure 7. Flow cytometry and scRNA-seq data analyses of HPCS lineage tracing. a-b,** mScarlet$^+$/EpCAM$^+$ (HPCS/all cancer cells, **a**) and GFP$^+$/EpCAM$^+$ (Traced cells/all cancer cells, **b**) at 3 days (3d) or 14 days (14d) of lineage tracing (6 to 8 week). N = 4 mice. Mann Whitney U test. **c-d,** mScarlet$^+$/EpCAM$^+$ (HPCS/all cancer cells, **c**) and GFP$^+$/EpCAM$^+$ (Traced cells/all cancer cells, **d**) at 3 days (3d) or 14 days (14d) of lineage tracing (12 to 14 week). N = 4 mice. Mann Whitney U test. **e,** Cell state overview from combined tracing experiments from **Fig. 2a**. scRNA-seq data from HPCS lineage-traced cells are plotted at the indicated timepoints. **f,** Stacked bar graphs showing the distribution of cell states across individual mice from HPCS tracing experiments shown as stacked bar graphs as in **Fig. 2d, g**. Error bars are SEM.

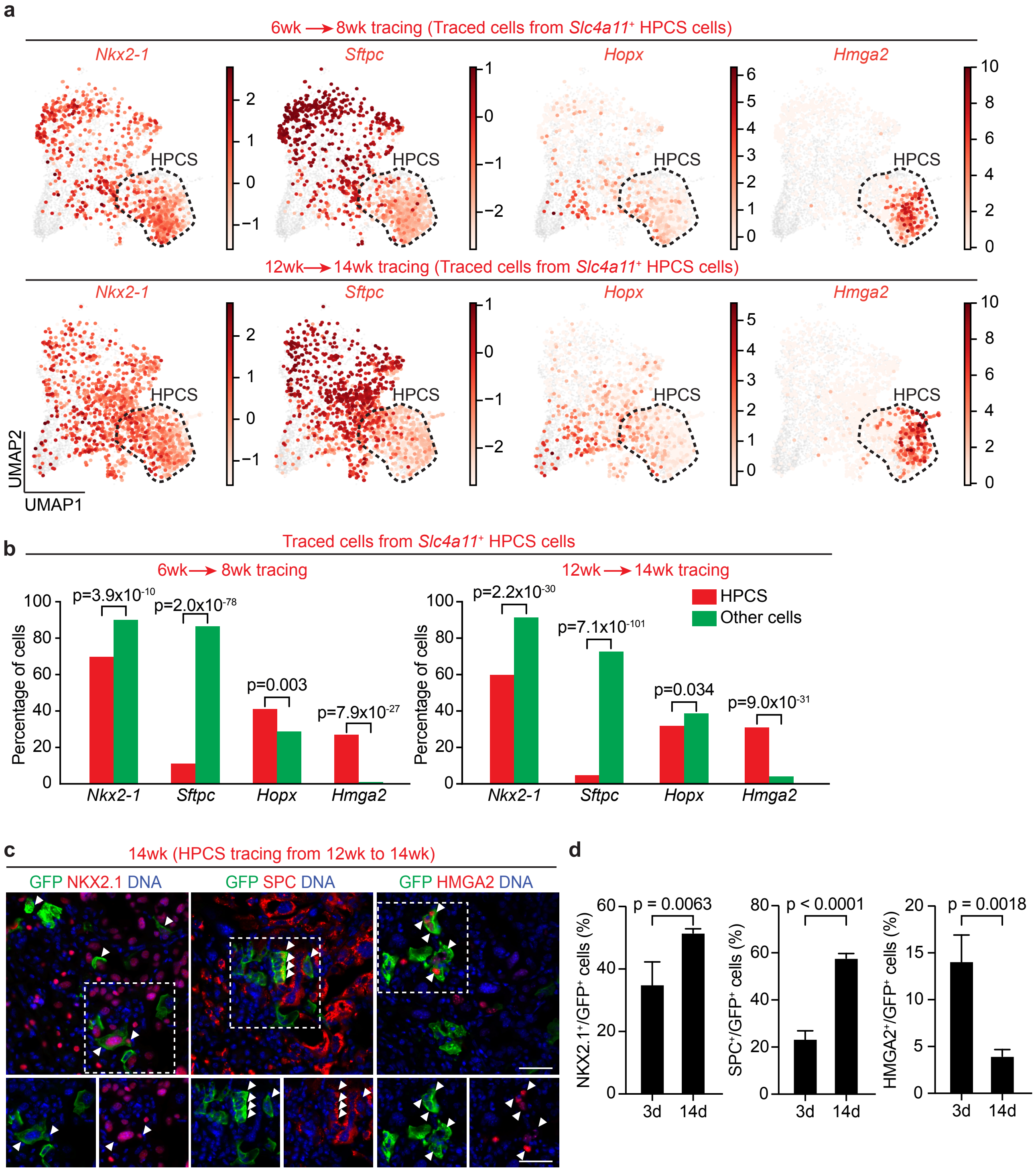




Chan, Pan et al.,
Extended Data Fig. 8

**Extended Data Figure 8. scRNA-seq data analyses and immunofluorescence imaging examining traced HPCS cells together with marker genes. a,** Distribution of *Nkx2-1*, *Sftpc*, *Hopx,* and *Hmga2* expression from traced LUAD cells at 8 weeks (6 to 8 week tracing, *top*) or 14 weeks (12 to 14 week tracing, *bottom*) PTI. **b,** Percentage of HPCS (red) or other cell states (green) expressing the indicated genes in traced LUAD cells at 8 weeks (6 to 8 week tracing, *left*) or 14 weeks (12 to 14 week tracing, *right*) PTI. Fisher's exact test. **c,** IF images showing co-staining of GFP (green) with either NKX2.1 (*left*, red), SPC (*middle*, red), or HMGA2 (*right*, red). Outlined areas shown as individual insets below their respective images. Scale bar: 20 µm. **d,** Percentage of NKX2.1$^{+}$ (*left*), SPC$^{+}$ (*middle*) or HMGA2$^{+}$ (*right*) cells in GFP$^{+}$ LUAD cells quantified from IF co-staining of LUAD tissues harvested at 14 weeks PTI after 3 (3d) and 14 (14d) days post tracing. NKX2.1: n = 21 tumors from 3 mice (3d) and n = 111 tumors from 3 mice (14d). SPC: n = 59 tumors from 3 mice (3d) and n = 89 tumors from 3 mice (14d). HMGA2: n = 84 tumors from 3 mice (3d) and n = 35 tumors from 3 mice (14d). Mann-Whitney U test. Error bars are SEM.

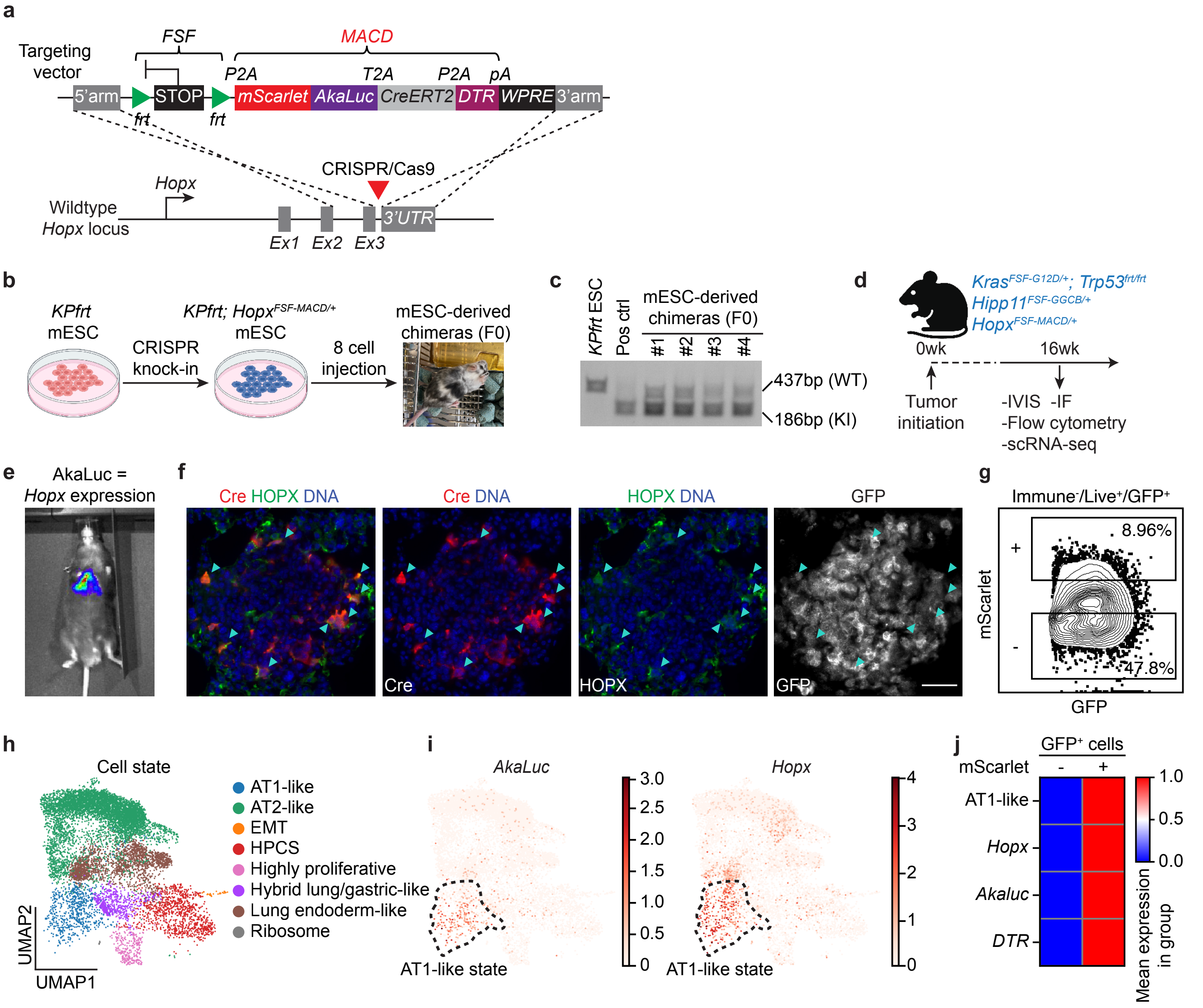


Chan, Pan et al., Extended Data Fig. 9

**Extended Data Figure 9. Construction of the *Hopx*$^{MACD/+}$ reporter allele. a,** Map of *Hopx*$^{FSF\text{-}MACD/+}$ reporter and targeting strategy by CRISPR/Cas9 mediated HDR. Strategy for generating the genetically engineered *Hopx*$^{MACD/+}$ knock-in (KI) reporter enabling lineage tracing and ablation of *Hopx*$^{+}$ AT1-like cells in *KP* LUAD: the *frt-stop-frt-P2A-mScarlet-T2A-Akaluc-P2A-CreERT2-P2A-DTR-WPRE* (*FSF-MACD*) reporter construct was knocked in frame into the stop codon of exon 3 of the *Hopx* gene in the presence of CRISPR/Cas9 mediated gene editing. **b,** Experimental outline to generate *Kras*$^{FSF\text{-}G12D/+}$*; Trp53*$^{frt/frt}$*; Hopx*$^{FSF\text{-}MACD/+}$ reporter mouse. *FSF-MACD* reporter construct was knocked into *KPfrt* mESCs, and the correctly targeted mESC clones were microinjected into 8-cell stage embryos to obtain mESC-derived chimeras. **c,** Gel electrophoresis of the PCR products from the parental *KPfrt* mESCs, a correctly targeted mESC clone (positive control), or mESC-derived chimeras using primer pairs detecting either wild-type (WT, 437 bp, *top*) or knock-in (KI, 186 bp, *bottom*) alleles, respectively. **d,** Experimental design to validate the *Hopx*$^{FSF\text{-}MACD/+}$ reporter. Tumors were initiated by intubation of PGK-FlpO lentivirus in *KPfrt; Hipp11*$^{GGCB/+}$*; Hopx*$^{MACD/+}$ mice and LUAD tumors were harvested 16 weeks PTI for flow cytometry, scRNA-seq, and IF staining. **e,** Bioluminescent imaging of mice with autochthonous *KPfrt; Hipp11*$^{GGCB/+}$*; Hopx*$^{MACD/+}$ LUAD tumors. AkaLuc bioluminescence equals *Hopx* expression as indicated. **f,** Representative IF images of mScarlet, HOPX, and GFP showing co-localization of mScarlet with HOPX (arrowheads) in GFP$^{+}$ tumor cells. Scale bar: 50 µm. **g,** Representative flow cytometry plot showing the GFP$^{+}$ tumor cells with mScarlet$^{+}$ or mScarlet$^{-}$ populations (highlighted regions) sorted from *KPfrt; Hipp11*$^{GGCB/+}$*; Hopx*$^{MACD/+}$ LUAD tumors 16 weeks PTI for scRNA-seq. N = 2 mice. **h,** LUAD cell states from the sorted cells shown in panel (**g**). **i,** Gene expression of *AkaLuc (left)* or *Hopx* (*right*) from GFP$^{+}$ tumor cells sorted from *KPfrt; Hipp11*$^{GGCB/+}$*; Hopx*$^{MACD/+}$ mice. **j,** Heatmap of scaled expression values for the AT1-like gene expression program, *Hopx*, or reporter cassette (as measured by *AkaLuc* and *DTR* expression) from sorted mScarlet$^{+}$/GFP$^{+}$ or mScarlet$^{-}$/GFP$^{+}$ LUAD cancer cells.

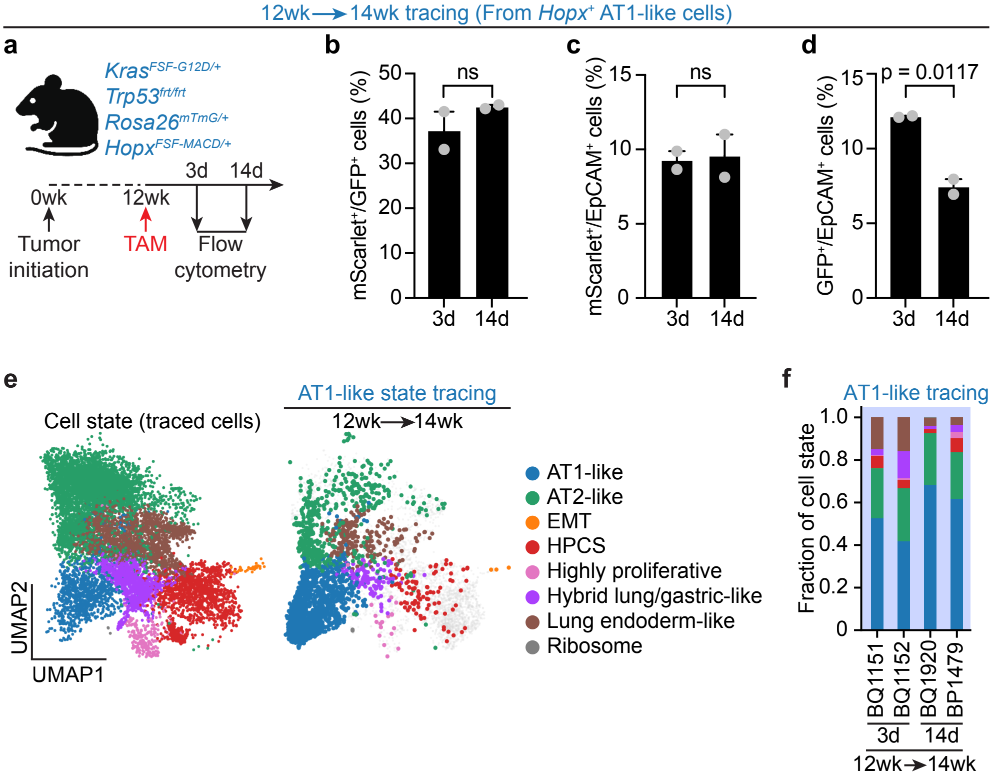


Chan, Pan et al.,
Extended Data Fig. 10

**Extended Data Figure 10. Flow cytometry and scRNA-seq data analyses of AT1-like cell state lineage tracing. a,** Experimental design to lineage-trace *Hopx*$^{+}$ AT1-like LUAD cells in *Kras*$^{FSF\text{-}G12D/+}$; *Trp53*$^{frt/frt}$; *Rosa26*$^{mTmG/+}$; *Hopx*$^{FSF\text{-}MACD/+}$ mice. At 12 weeks PTI, tumor-bearing mice were administered a single dose of tamoxifen (TAM; 20 mg/kg), which caused cells with high *Hopx*$^{MACD/+}$ expression to switch from tdTomato to GFP expression. Tumors were harvested at 3 days (3d) and 14 days (14d) for flow cytometry analysis. **b-d,** mScarlet$^{+}$/GFP$^{+}$ (AT1-like/all traced cells, **b**), mScarlet$^{+}$/EpCAM$^{+}$ (AT1-like/all cancer cells, **c**), and GFP$^{+}$/EpCAM$^{+}$ (Traced cells/all cancer cells, **d**) cancer cells at 3 or 14 days of lineage tracing. N = 2 mice per time point. Two-sample t-test. Error bars are SEM. **e,** Analysis of the combined data from tracing experiments from **Fig. 2a**. (*left*) and scRNA-seq data from AT1-like lineage-traced cells (*right*). **f,** Stacked bar graphs showing the distribution of cell states across individual mice from the AT1-like tracing experiments shown in in (**e**) and stacked bar graphs in **Fig 2j**.

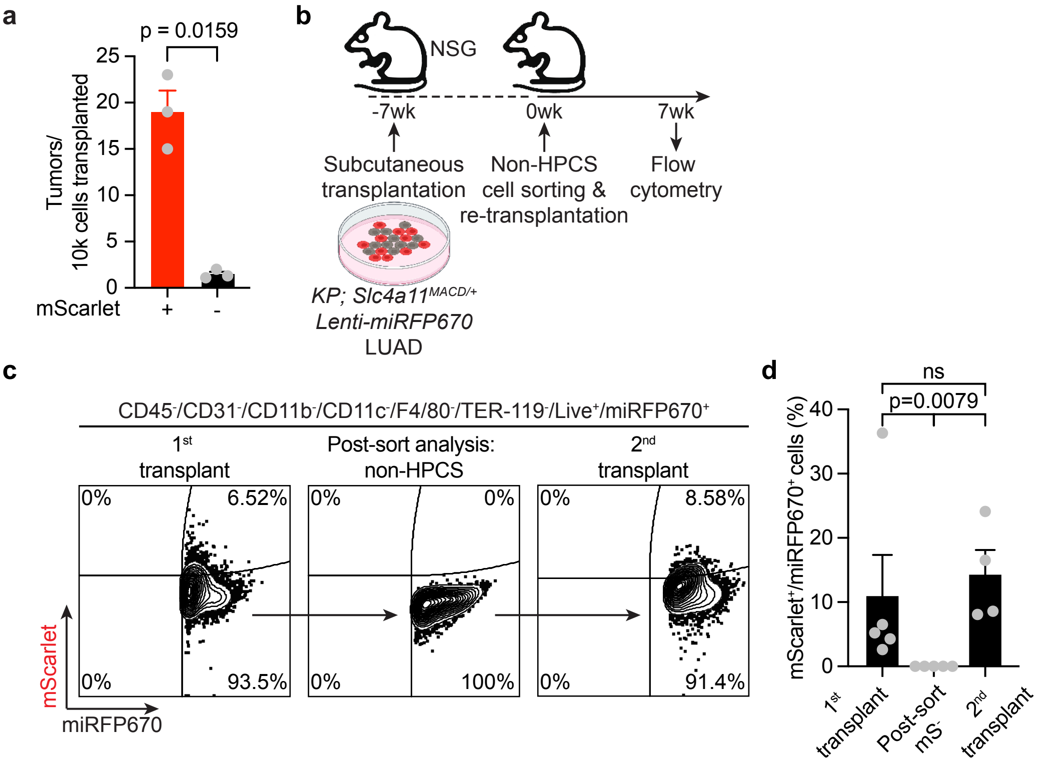




Chan, Pan et al.,
Extended Data Fig. 11

**Extended Data Figure 11. Serial transplantation studies demonstrate that non-HPCS cells can transition into HPCS. a,** Tumor number per 10,000 (10k) cells transplanted. Transplanted cells were either mScarlet$^-$ or mScarlet$^+$ (**Fig. 2m**). N = 3 mice. Welch's t test. **b,** Experimental schema to examine the emergence of HPCS from non-HPCS derived transplantation. A *KPfrt; Slc4a11*$^{MACD/+}$ LUAD cell line expressing the miRFP670 reporter was subcutaneously transplanted into NSG mice (-7wk). mScarlet$^-$/miRFP670$^+$ cells (non-HPCS) were sorted and subcutaneously transplanted into NSG mice (0wk). Tumors were harvested for flow cytometry analysis (7wk). **c,** Representative flow cytometry plot of mScarlet and miRFP670 expression in tumors of the first transplantation (1$^{st}$ transplant), sorted non-HPCS cells (post-sort analysis), and second transplantation (2$^{nd}$ transplant). Cells were gated as CD45$^-$/CD31$^-$/CD11b$^-$/CD11c$^-$/F4/80$^-$/TER-119$^-$/DAPI$^-$(Live$^+$)/miRFP670$^+$. **d,** Average percentage of mScarlet$^+$ cells in miRFP670$^+$ populations from the indicated groups. Post-sort mS$^-$: Post sort analysis on non-HPCS cells as in (**c**). N = 4-5 mice. Mann-Whitney U test. Error bars are SEM.

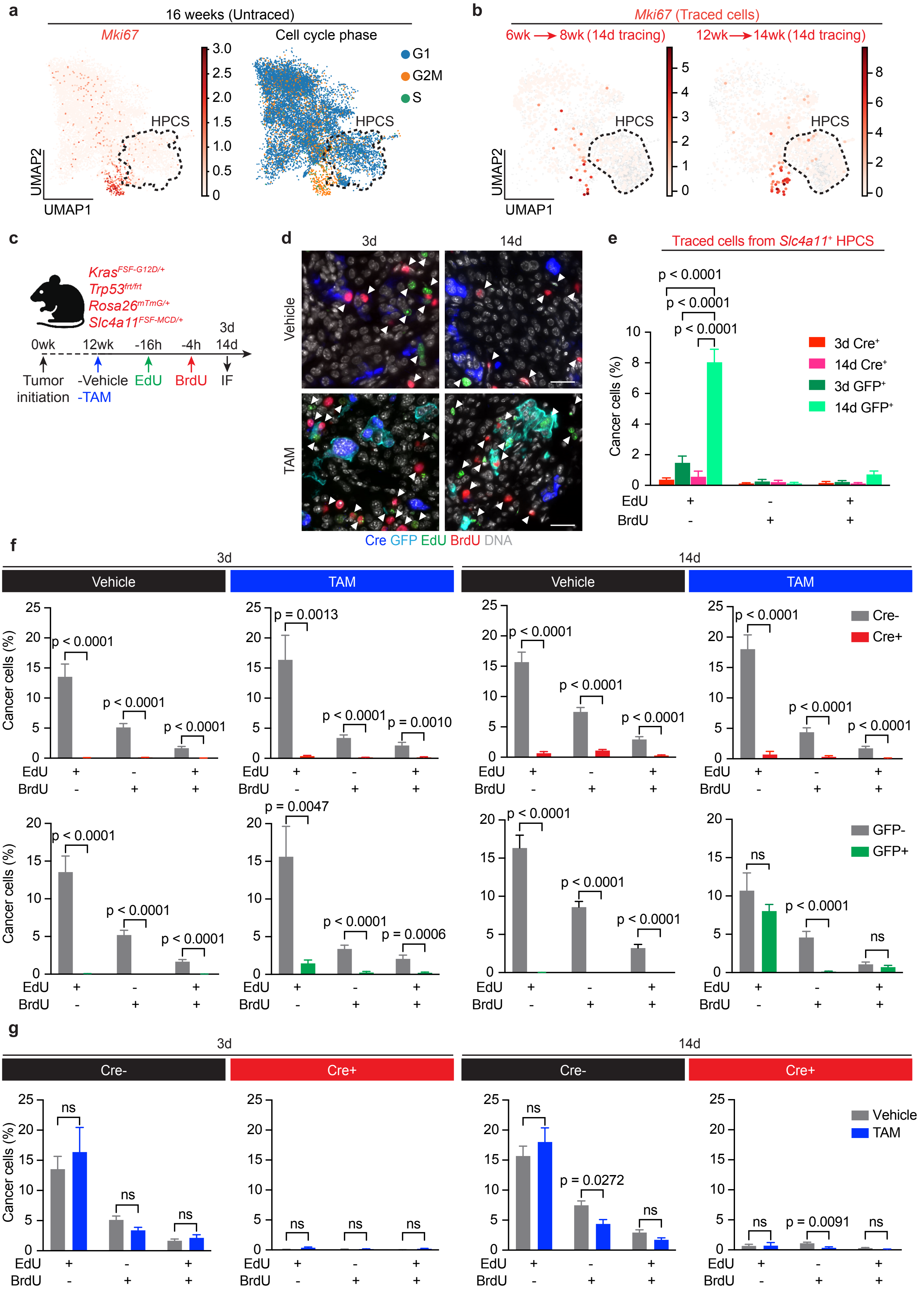


Chan, Pan et al.,
Extended Data Fig. 12

**Extended Data Figure 12. Single cell proliferation data and *in vivo* EdU/BrdU incorporation assays in lineage-traced HPCS cells. a,** *Mki67* expression (*left*) and the predicted cell cycle phase (*right*) in *KPfrt; Hipp11*$^{GGCB/+}$*; Slc4a11*$^{MCD/+}$ mice at 15-16 weeks PTI. **b,** *Mki67* expression in early-grade neoplasias and established adenocarcinomas, 14-days post-tamoxifen (TAM) labeling, in cells harvested at 8 weeks (*left*) and 14 weeks (*right*) PTI. **c,** Experimental design for EdU/BrdU dual labeling in *KPfrt; Rosa26*$^{mTmG/+}$*; Slc4a11*$^{MCD/+}$ mice. At 12 weeks PTI, tumor-bearing mice received a single dose of TAM (200 mg/kg), enabling lineage tracing of HPCS cells through a switch from tdTomato to GFP expression. EdU and BrdU were administered at 16 hours and 4 hours, respectively, before tumor harvest at 3 days (3d) and 14 days (14d) post lineage tracing for IF analysis. **d,** Images showing IF staining of Cre (HPCS, blue), GFP (traced-HPCS, turquoise), EdU (green), BrdU (red), and DNA (DAPI, white) with (*bottom*) or without (*top*) TAM treatment at 3d (*left*) or 14d (*right*) post lineage tracing. EdU$^{+}$ and BrdU$^{+}$ cells are pointed with arrowheads. Scale bar: 30 μm. **e,** Percentage of EdU$^{+}$/BrdU$^{-}$, EdU$^{-}$/BrdU$^{+}$, and EdU$^{+}$/BrdU$^{+}$ in Cre$^{+}$ (HPCS) or GFP$^{+}$ (lineage-traced) cells at 3d or 14d following lineage tracing. N = 30 tumors from 4 mice. Two-way ANOVA. **f,** Percentage of EdU$^{+}$/BrdU$^{-}$, EdU$^{-}$/BrdU$^{+}$, and EdU$^{+}$/BrdU$^{+}$ comparing Cre$^{+}$ (HPCS) and Cre$^{-}$ (non-HPCS) cells (*top*) and GFP$^{+}$ (traced) and GFP$^{-}$ (non-traced) cells (*bottom*) at 3d or 14d following TAM or vehicle treatment. N = 30 tumors from 4 mice. Two-way ANOVA. **g,** Percentage of EdU$^{+}$/BrdU$^{-}$, EdU$^{-}$/BrdU$^{+}$, and EdU$^{+}$/BrdU$^{+}$ comparing vehicle and TAM groups between Cre$^{-}$ (non-HPCS) and Cre$^{+}$ (HPCS) cells at 3d or 14d following treatment. N = 30 tumors from 4 mice. Two-way ANOVA. Error bars are SEM.

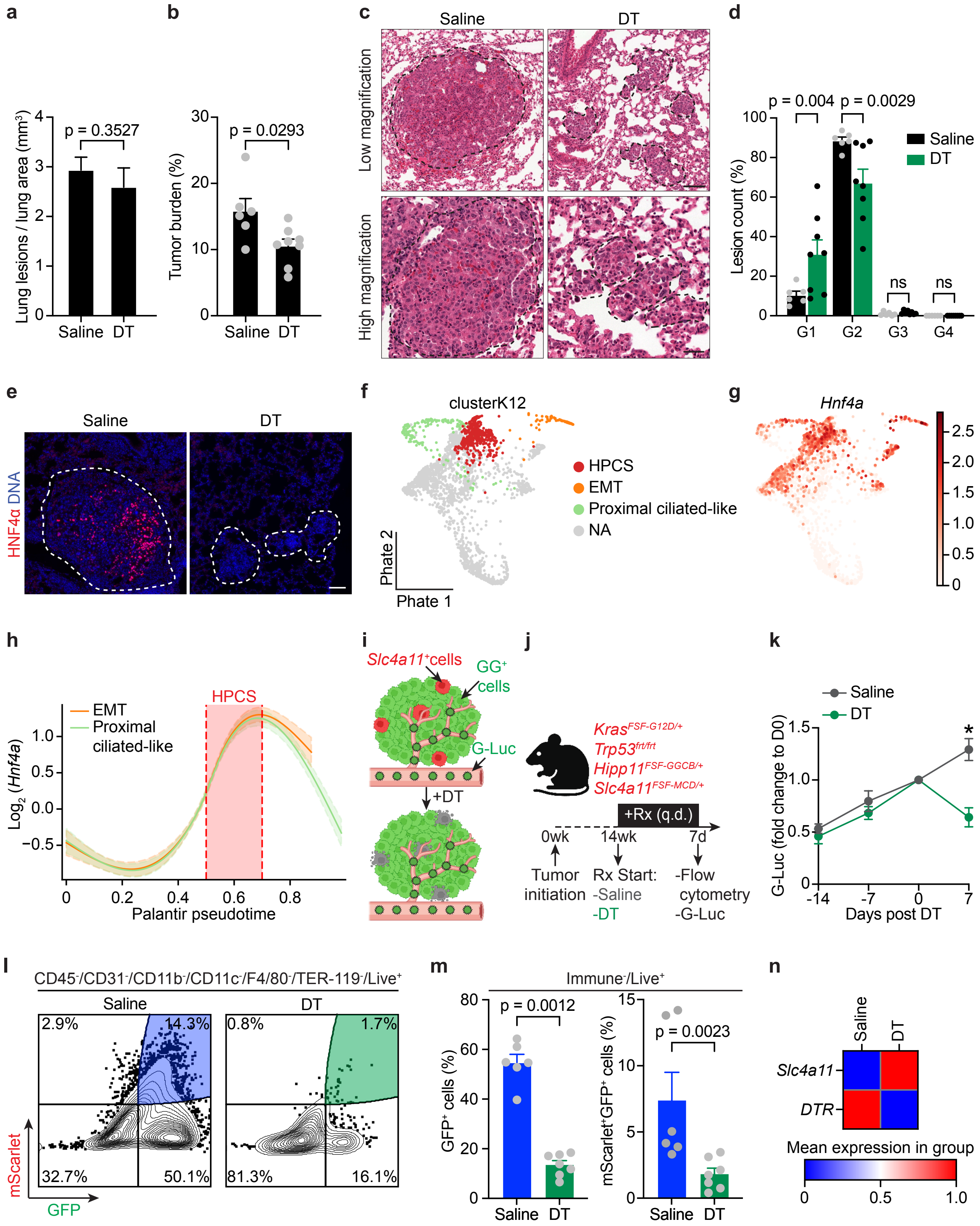




Chan, Pan et al.,
Extended Data Fig. 13

**Extended Data Figure 13. Histopathological and flow cytometry analyses of HPCS ablation in early neoplasias and established LUAD. a,** Number of tumors normalized by lung area in saline vs. DT treated groups. N = 10 lung areas from 3 mice. Mann-Whitney U test. **b,** Tumor burden (total tumor area / total lung area) in saline vs. DT treated groups. N = 6-8 mice. Mann-Whitney U test. **c,** Representative H&E stained lung sections showing tumor boundaries (dashed lines) in saline control vs. DT-treated mice. Scale bar: 100 µm (low magnification) and 40 µm (high magnification). **d,** Distribution of tumor numbers across tumor grades in saline vs. DT treated groups. N = 6-8 mice. Two-way ANOVA. **e,** Images showing IF staining of HNF4α (red) and DNA (blue) from lung sections in saline control vs. DT-treated mice. Scale bar: 100 µm. **f,** Predicted end cell states (EMT, Proximal ciliated-like) and the HPCS projected on the original PHATE[100] map from Marjanovic et. al.[4]. **g,** *Hnf4a* expression. **h,** Gene expression trajectories for *Hnf4a* plotted over Palantir[85] *pseudotime,* with estimated time spent in the HPCS shaded in red. **i-j,** Experimental design to test the effect of ablating *Slc4a11*$^{+}$ HPCS on tumor growth in autochthonous *KPfrt; Hipp11*$^{GGCB/+}$*; Slc4a11*$^{MCD/+}$ LUAD bearing mice as determined by longitudinal G-Luc activity monitoring. Experimental schema of LUAD bearing mice 14 weeks PTI were administered saline or DT (50 µg/kg, daily) and analyzed at 7 days as indicated. **k,** Measurement of G-Luc activity (secreted from GG$^{+}$ tumor cells) shown as fold change at indicated time points normalized to day 0 post DT treatment. N = 3 mice per group. Welch's t test. **l,** Representative flow cytometry plot of saline- and DT-treated (7 day) tumors. Shaded regions indicate gated populations analyzed. **m,** Quantification of total tumor burden (GFP$^{+}$ cells, *left*) or HPCS (mScarlet$^{+}$/GFP$^{+}$ cells, *right*) as a percentage of live, immune-negative cells [CD45$^{-}$/CD31$^{-}$/CD11b$^{-}$/CD11c$^{-}$/F480$^{-}$/TER-119$^{-}$/Helix NP NIR$^{-}$(Live)$^{+}$]. N = 6 (Saline) and 7 (DT) mice. Mann-

Whitney U test. **n,** Heatmap of scaled mean gene expression for *DTR* ($Slc4a11^{MCD/+}$ allele) and native *Slc4a11* in saline- vs. DT-treated $GFP^{+}$ cancer cells. Error bars are SEM.

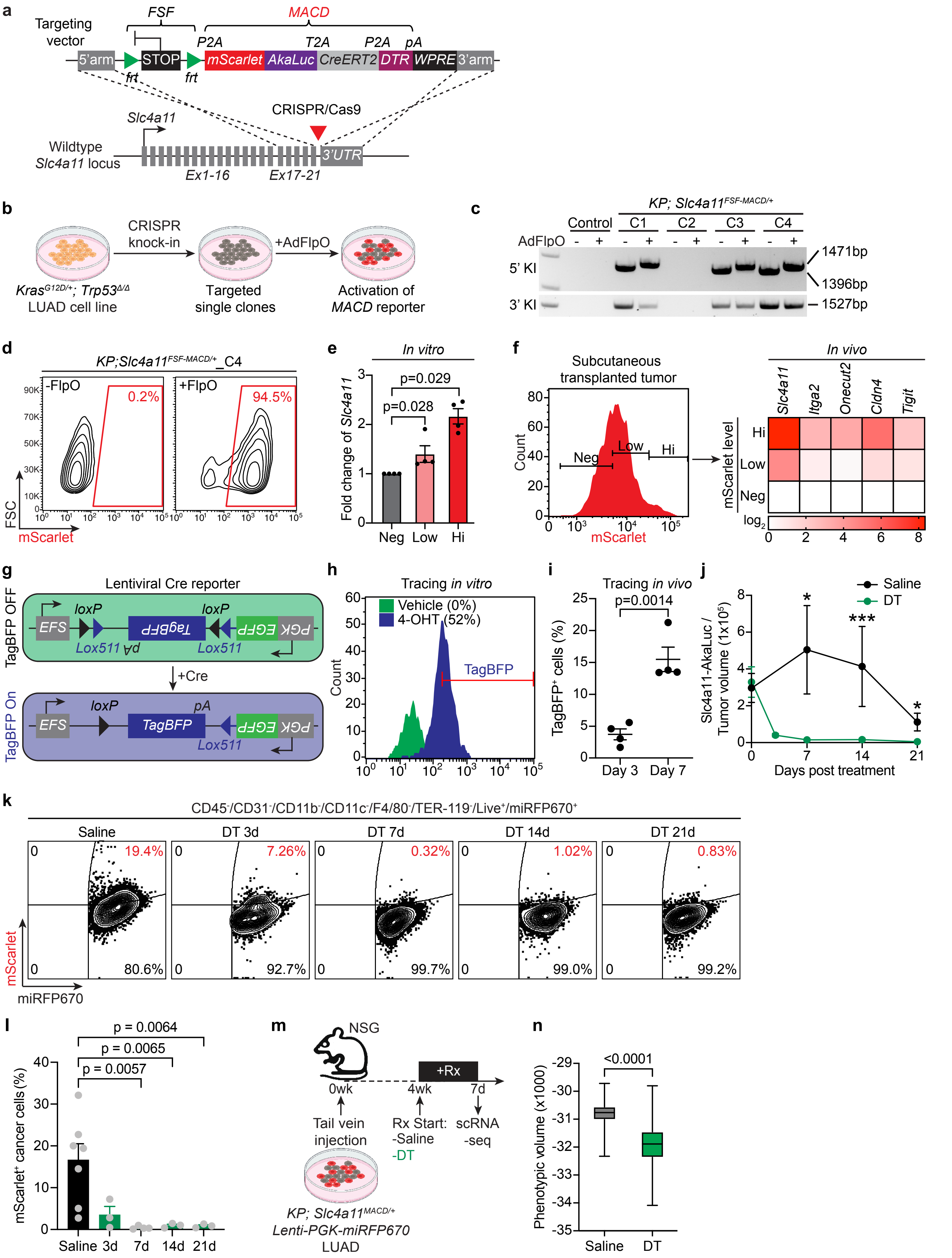


Chan, Pan et al., Extended Data Fig. 14

**Extended Data Figure 14. Construction, validation, and cytoablation of a *Slc4a11*$^{MACD/+}$ reporter cell line and allotransplants. a,** Vector map of *Slc4a11*$^{FSF\text{-}MACD/+}$ and targeting strategy of the reporter by CRISPR/Cas9 mediated homology directed repair (HDR). *frt-stop-frt-P2A-mScarlet-T2A-Akaluc-P2A-CreERT2-P2A-DTR-WPRE* (*FSF-MACD*) reporter construct was knocked in frame into the *Slc4a11* locus in the presence of CRISPR/Cas9 mediated gene editing. Akaluc: Akaluciferase; P2A and T2A: short polypeptide cleavage site. **b,** Experimental design to generate *Slc4a11*$^{MACD/+}$ reporter cell line. A *Kras*$^{G12D/+}$*; Trp53*$^{-/-}$ (*KP*) LUAD cell line was transfected with the *Slc4a11-MACD* targeting vector in the presence of U6-sg*Slc4a11*-EFS-Cas9 and targeted single clones were transduced with Adenoviral-CMV-FlpO (AdFlpO) to remove the *FSF* cassette and activate the *MACD* reporter. **c,** Gel electrophoresis of the PCR products from targeted clones before and after AdFlpO using primers detecting knock-in of the 5' (5' KI, *top*) and 3' (3' KI, *bottom*) arms, respectively. Successful removal of the *FSF* cassette resulted in an increase of band size in the PCR detection of the 5' arm knock-in. **d,** Flow cytometry analysis of mScarlet expression in targeted single clone 4 (*Slc4a11-FSF-MACD*_C4) of *KP* LUAD before and after AdFlpO transduction. **e,** Relative expression of *Slc4a11* transcripts (shown as fold change) measured by qRT-PCR in mScarlet negative (Neg), low (Low) or high (Hi) sorted cell populations from *Slc4a11-FSF-MACD*_C4 *KP* LUAD cells transduced with AdFlpO. N = 4. Mann-Whitney U test. **f,** *Left*: Representative flow cytometry plot of mScarlet expression from dissociated subcutaneously transplanted *Slc4a11-FSF-MACD*_C4 *KP* LUAD tumor. Cells were sorted for negative, low, or high expression of mScarlet as indicated. *Right*: Heatmap showing the relative expression of indicated genes (columns) measured by qRT-PCR from the sorted cell populations (rows). N = 2 biological replicates, scale is log2 as indicated. **g,** Schematic of lentiviral Cre recombinase reporter construct for the testing of CreER$^{T2}$ functionality from the *Slc4a11*$^{FSF\text{-}MACD/+}$

*KP* LUAD reporter. *Slc4a11*$^{FSF\text{-}MACD/+}$ *KP* LUAD cells infected with the lentiviral Cre reporter expresses GFP alone and upon exposure to 4-hydroxytamoxifen (4-OHT) expresses TagBFP in *Slc4a11*$^{+}$ cells. **h,** Flow cytometry analysis of TagBFP expression from *Slc4a11-FSF-MACD KP* LUAD cells infected with the lentiviral Cre reporter treated with either vehicle or with 4-OHT (1 µM) for 72 hours. **i,** Percentage of TagBFP$^{+}$ tumor cells from dissociated subcutaneous transplanted *Slc4a11*$^{FSF\text{-}MACD/+}$ *KP* LUAD (with lentiviral Cre reporter) at day 3 or day 7 post tamoxifen (TAM, 200 mg/kg) treatment. N = 4 mice. Unpaired t-test. **j,** *Slc4a11*-AkaLuc bioluminescence signal intensity normalized to tumor volume in saline- vs. DT-treated allograft tumors (n = 8 tumors each from 4 mice per group). Measurements were taken at the indicated time points. Welch's t test. **k,** A *KPfrt*; *Slc4a11*$^{MACD/+}$ LUAD cell line expressing the miRFP670 reporter was subcutaneously transplanted into NSG mice. Flow cytometry analysis of mScarlet versus miRFP670 expression in transplanted LUAD following saline or DT treatment for the indicated time. Cells were gated as CD45$^{-}$/CD31$^{-}$/CD11b$^{-}$/CD11c$^{-}$/F4/80$^{-}$/TER-119$^{-}$/DAPI$^{-}$ (live$^{+}$)/miRFP670$^{+}$. **l,** Percentage of mScarlet$^{+}$ cells within the miRFP670$^{+}$ tumor cell population as gated in (**k**) from subcutaneous allotransplanted LUAD cells treated with either saline or DT over the indicated time course. N = 7 (saline), 3 (3d), 4 (7d), 3 (14d), and 3 (21d) mice. One-way ANOVA. **m,** Experimental design to test the effects of ablating *Slc4a11*$^{+}$ HPCS cells on tumor heterogeneity in intravenous transplants of *KPfrt; Slc4a11*$^{MACD/+}$ LUAD reporter cells. Four weeks after transplantation, mice were administered saline or DT (25 µg/kg, q.o.d.). Tumors were harvested after 7 days post-treatment and miRFP670$^{+}$ cancer cells were sorted for scRNA-seq. **n,** Phenotypic volume of cells in saline or DT treated groups. N = 2-3 mice. Box plots are min-max. Welch's t test. Error bars are SEM.

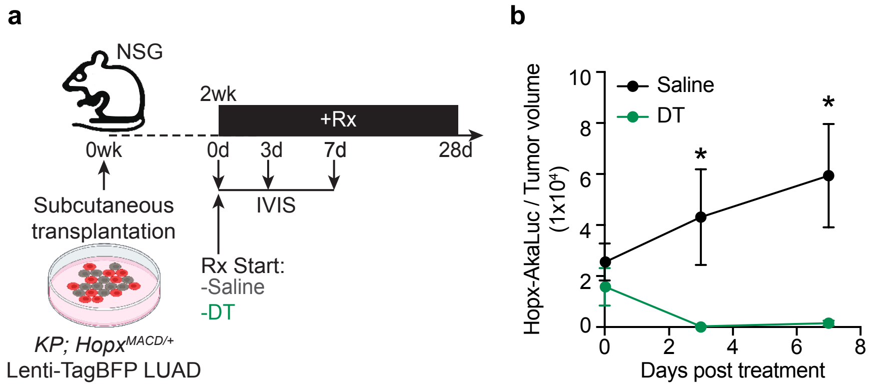




Chan, Pan et al.,
Extended Data Fig. 15

**Extended Data Figure 15. Cytoablation of *Hopx*$^{+}$ AT1-like cells in LUAD reporter transplant models. a,** Experimental design to test the effects of ablating *Hopx*$^{+}$ AT1-like cells on tumor growth in subcutaneous *KPfrt; Hopx*$^{MACD/+}$ LUAD reporter allografts. Two weeks after transplantation, allograft bearing mice were administered saline or DT (25 µg/kg, q.o.d.). Tumor volume measurement and AkaLuc bioluminescence signal detection were performed at the indicated time point. IVIS: *in vivo* imaging system. **b,** *Hopx*-AkaLuc bioluminescence signal intensity normalized to tumor volume in saline-treated (n = 10 tumors from 5 mice) vs. DT-treated (n = 8 tumors from 4 mice) allografts. Welch's test. **c,** Tumor volume at indicated timepoints in allografts in mice administered either saline or DT. N = 8 tumors from 4 mice per group. Welch's test. Error bars are SEM.

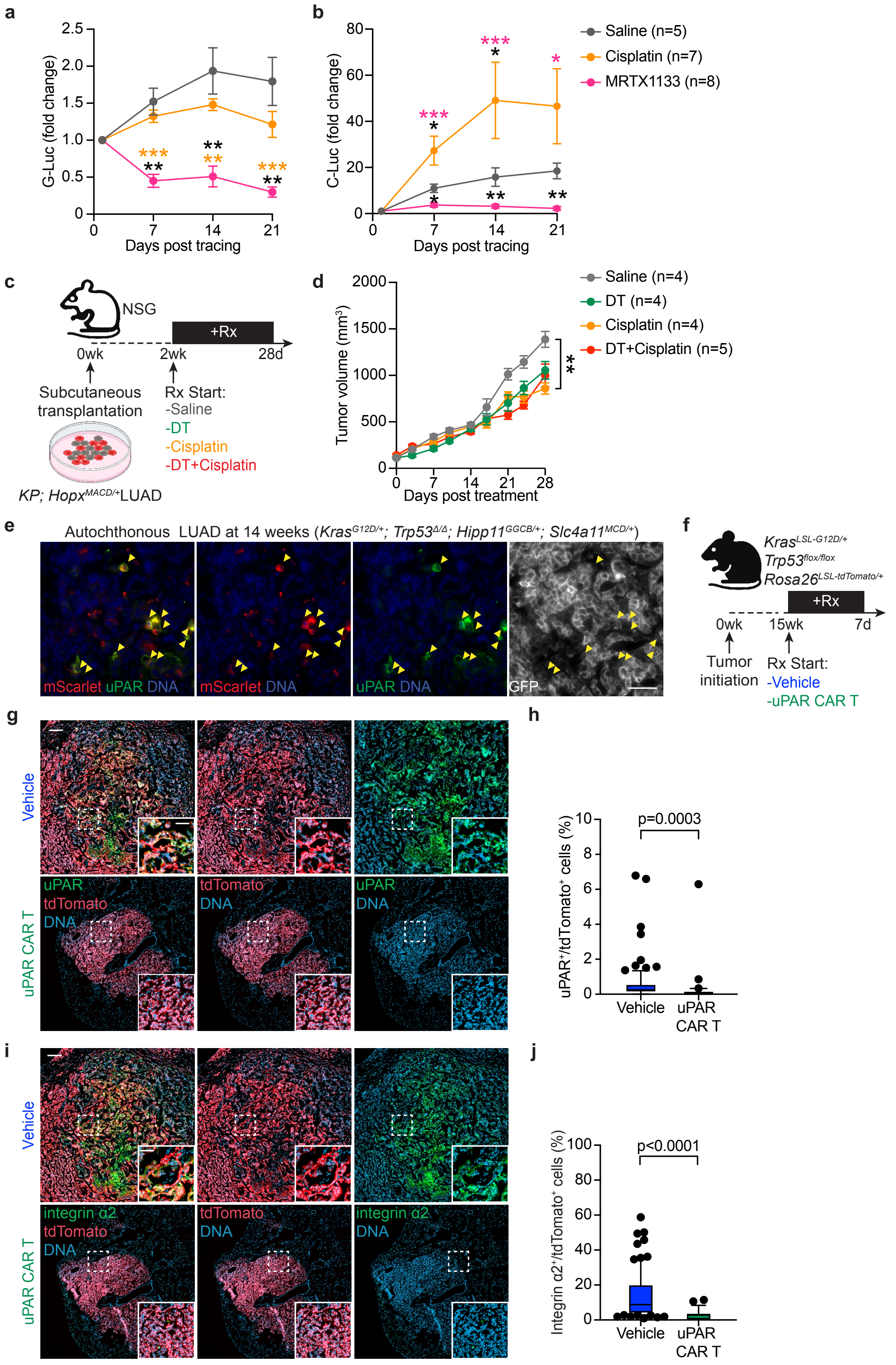




Chan, Pan et al.,
Extended Data Fig. 16

**Extended Data Figure 16. G-Luc and C-Luc activity following chemotherapy or targeted therapy treatment related to Fig. 4a-b; Combination of *Hopx*$^{+}$ cytoablation with cisplatin treatment related to Fig. 4e-g; uPAR expression and uPAR CAR-T cell-mediated HPCS eradication related to Fig. 4h-k. a-b,** Data relating to Fig. 4a-b. G-Luc (**a**) and C-Luc (**b**) activity at indicated time points (normalized to day 1) in saline (n = 5 mice), cisplatin (n = 7 mice), or MRTX1133 (n = 8 mice) treated groups. Note: Fold change of C-Luc under MRTX1133 therapy: 4x (7d), 3x (14d), and 2x (21d). Welch's t-test. Error bars are SEM. **c-d,** Data relating to Fig. 4e-g. **c,** Experimental design to test the effect of ablating *Hopx*$^{+}$ AT1-like LUAD cells by DT (25 µg/kg, q.o.d.) in the context of either saline control or cisplatin chemotherapy (1.5 mg/kg, every 3 days) in subcutaneous *KPfrt; Hopx*$^{MACD/+}$ LUAD reporter allografts. **d,** Tumor volume of subcutaneous *KPfrt; Hopx*$^{MACD/+}$ LUAD allografts subjected to saline control, DT, or cisplatin, alone, or in combination, as in (**c**). N = 8 tumors from 4 mice per group. Two-way ANOVA. Error bars are SEM. **e-j,** Data relating to Fig. 4h-k. **e,** mScarlet, uPAR, and GFP immunofluorescence. Yellow arrowheads indicate co-localization of mScarlet with uPAR (yellow arrowheads) in GFP$^{+}$ tumor cells. Scale bar: 20 µm. **f,** Outline of the experimental design to evaluate uPAR CAR-T cells (2x10$^{6}$ cells/mouse) in autochthonous *KP*; *Rosa26*$^{tdTomato/+}$ LUAD with 1 dose of either vehicle or uPAR CAR-T cells (2x10$^{6}$ cells/mouse). **g,** Representative IF images of autochthonous *KP; Rosa26*$^{tdTomato/+}$ LUAD tumors treated with vehicle versus uPAR CAR T cells. Cryosections are stained with anti-uPAR antibodies and depict endogenously expressed tdTomato. Inset depicts boxed regions. Large image scalebar: 200 µm, inset scalebar: 60 µm. **h,** Quantification of uPAR expressing tdTomato$^{+}$ tumor cells in vehicle (n = 93 primary tumors from one mouse) vs. uPAR CAR T cell (n = 31 primary tumors from one mouse) treated mice shown as a boxplot, with whiskers representing the 10$^{th}$ and 90$^{th}$ percentiles. Kruskal Wallis test. **i,** Representative IF images of integrin α2 and tdTomato in autochthonous *KP; Rosa26*$^{tdTomato/+}$ LUAD bearing mice treated with vehicle or uPAR CAR T cells. Inset depicts regions boxed with a dashed line. Large image scalebar: 200 μm, inset scalebar: 60 μm. **j,** Boxplot of integrin $\alpha2^{+}$/tdTomato$^{+}$ tumor cell percentages in vehicle (n = 83 primary tumors from one mouse) vs. uPAR CAR T (n = 25 primary tumors from one mouse) treated mice, with whiskers representing the $10^{th}$ and $90^{th}$ percentiles. Kruskal Wallis test.

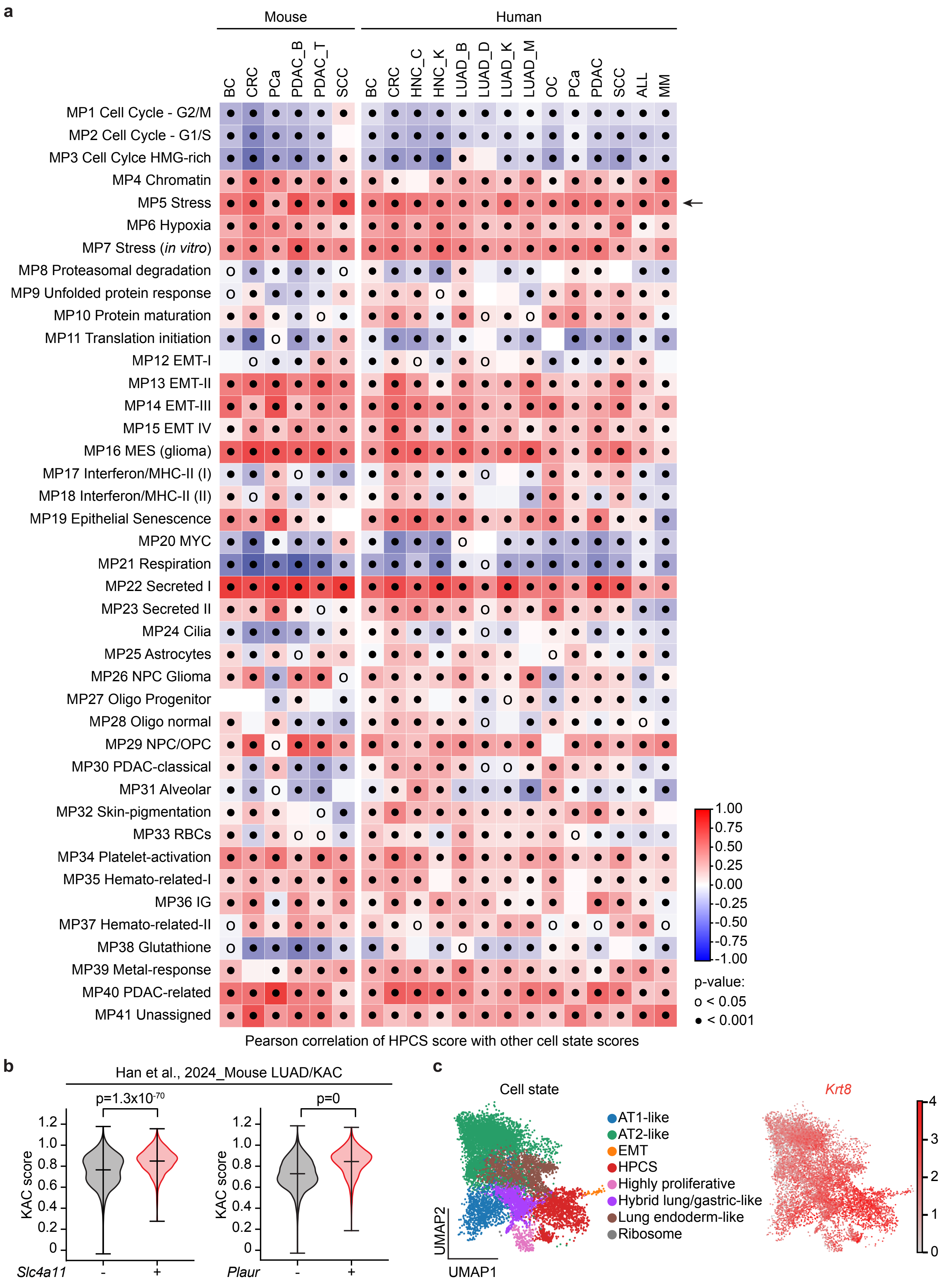




Chan, Pan et al.,
Extended Data Fig. 17

**Extended Data Figure 17. Analysis of a HPCS-like state across cancers. Related to Fig 5. a,** Heatmap of Pearson correlations calculated between the HPCS and recurrent pan-cancer cell states (RBC: red blood cells, IG: immunoglobulins)[51], divided by cancer type (BC: breast cancer, CRC: colorectal cancer, HNC: head and neck cancer, LUAD: lung adenocarcinoma, OC: ovarian cancer, PCa: prostate adenocarcinoma, PDAC: pancreatic adenocarcinoma, SCC: cutaneous squamous cell carcinoma, ALL: acute lymphoblastic leukemia, MM: multiple myeloma). Arrows indicate correlations to the *Stress*-associated cell state. A full list of studies is listed in **Supplementary Table 5**. **b,** Violin plot of the *Krt8*$^+$ alveolar intermediate cell state (KAC)[5] score in *Slc4a11* (*left*) and *Plaur* (*right*) negative and positive cells. t-test. **c,** Expression of *Krt8*, a marker of the "KAC" subset of LUAD cells, in *KP* lung adenocarcinoma cells. Note broad, near-uniform expression of *Krt8* throughout all the LUAD cell states.

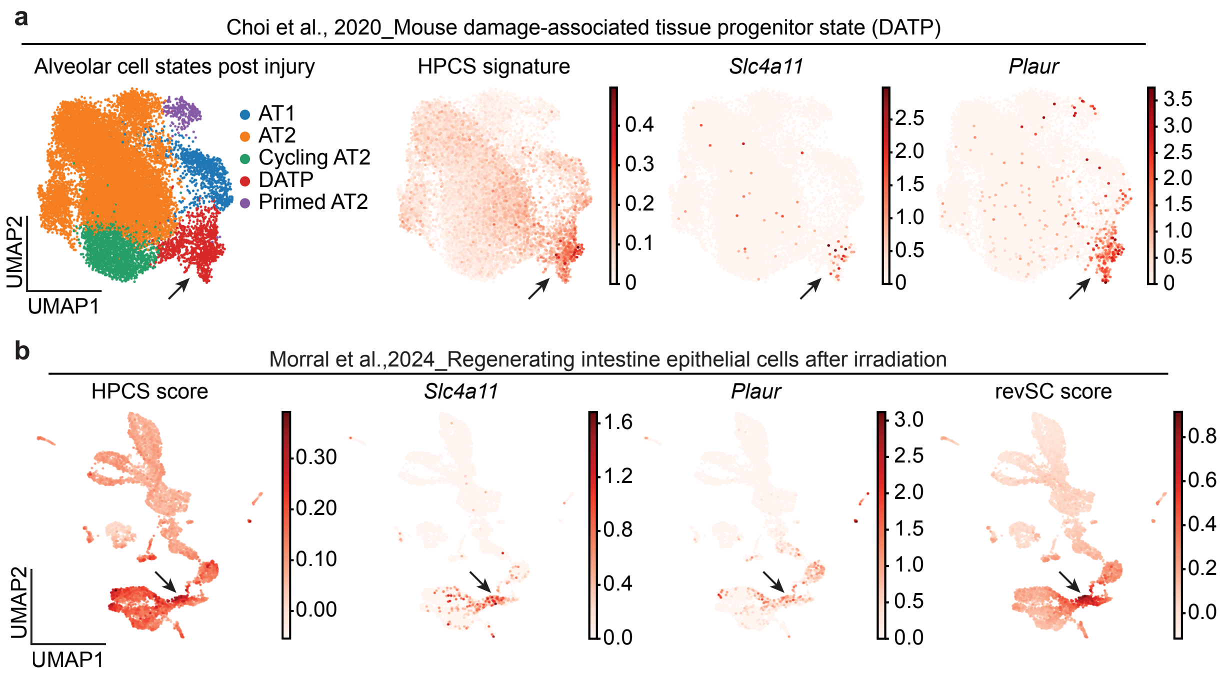




Chan, Pan et al.,
Extended Data Fig. 18

**Extended Data Figure 18. Analysis of scRNA-seq datasets from mouse injury models highlighting overlap between regenerative cell states, *Slc4a11*, and *Plaur.* Related to Fig. 5.** **a,** Classification of cell states (*far left*), expression of the HPCS program (*middle left*), *Slc4a11* (*middle right*) or *Plaur* (*far right*) in scRNA-seq data obtained from injured and non-injured primary lung tissue. Arrows point to the damage associated transient progenitor (DATP) cell state[55]. **b,** Distribution of the HPCS program (*far left*), expression of *Slc4a11* (*middle left*) or *Plaur* (*middle right*) in scRNA-seq data obtained from injured and non-injured primary mouse small intestine tissue[98]. Arrows point to the damage associated revival stem cell (revSC) cell state.
